# Four additional natural 7-deazaguanine derivatives in phages and how to make them

**DOI:** 10.1101/2023.04.13.536721

**Authors:** Liang Cui, Seetharamsingh Balalkundu, Chuanfa Liu, Hong Ye, Jacob Hourihan, Astrid Rausch, Christopher Hauß, Emelie Nilsson, Matthias Hoetzinger, Karin Holmfeldt, Weijia Zhang, Laura Martinez-Alvarez, Xu Peng, Denise Tremblay, Sylvain Moinau, Natalie Solonenko, Matthew B. Sullivan, Yan-Jiun Lee, Andrew Mulholland, Peter Weigele, Valérie de Crécy-Lagard, Peter C. Dedon, Geoffrey Hutinet

## Abstract

Bacteriophages and bacteria are engaged in a constant arms race, continually evolving new molecular tools to survive one another. To protect their genomic DNA from restriction enzymes, the most common bacterial defence systems, double-stranded DNA phages have evolved complex modifications that affect all four bases. This study focuses on modifications at position 7 of guanines. Eight derivatives of 7-deazaguanines were identified, including four previously unknown ones: 2’-deoxy-7-(methylamino)methyl-7-deazaguanine (mdPreQ_1_), 2’-deoxy-7-(formylamino)methyl-7-deazaguanine (fdPreQ_1_), 2’-deoxy-7-deazaguanine (dDG), and 2’-deoxy-7-carboxy-7-deazaguanine (dCDG). These modifications are inserted in DNA by a guanine transglycosylase named DpdA. Three subfamilies of DpdA had been previously characterized: bDpdA, DpdA1, and DpdA2. Two additional subfamilies were identified in this work: DpdA3, which allows for complete replacement of the guanines, and DpdA4, which is specific to archaeal viruses. Transglycosylases have now been identified in all phages and viruses carrying 7-deazaguanine modifications, indicating that the insertion of these modifications is a post-replication event. Three enzymes were predicted to be involved in the biosynthesis of these newly identified DNA modifications: 7-carboxy-7-deazaguanine decarboxylase (DpdL), dPreQ_1_ formyltransferase (DpdN), and dPreQ_1_ methyltransferase (DpdM), which was experimentally validated and harbors a unique fold not previously observed for nucleic acid methylases.

## INTRODUCTION

Because of their intrinsic properties, such as resistance to nucleases (*1*), or fluorescence quenching (*2*), 7-deazaguanine derivatives have long been employed in synthetic biology. Two of these derivatives are tRNA modifications, queuosine (Q) and archaeosine (G^+^). They are respectively involved in the avoidance of translational errors and in tRNA stabilization (*3*). Recently, 7-deazaguanine derivatives have been found in DNA as components of restriction/modification systems in bacteria (*4, 5*), and anti-restriction systems in phages (*4, 6, 7*). Epigenetic modifications are common among phages (*8–11*) to resists to various bacterial defense systems (*11–16*).

Members of a transglycosylase superfamily are responsible for the incorporation of 7-deazaguanine derivatives into both tRNA and DNA. Proteins of the Tgt subgroup modify tRNA, while DpdA subgroup proteins modify DNA (*3*), both by replacing the target guanine with a specific 7-deazaguanine derivative. Tgt enzyme 7-deazaguanine substrates differ between organisms. One of these substrates is queuine (q), which is inserted at position 34 of the GUN anticodon tRNAs in eukaryotes and in certain bacteria (*3*). 7-aminomethyl-7-deazaguanine (preQ_1_) is inserted at the same position in most bacteria. 7-cyano-7-deazaguanine (preQ_0_) is inserted at position 15 or 16 of many tRNAs in archaea (*3*). Similarly, all DpdA enzymes tested thus far insert preQ_0_ in DNA (*4, 5, 7*), and sequence specificity has been identified for one of them (*17*). DpdA homologs are divided in three groups: bacterial DpdA (bDpdA), and two phage DpdA (DpdA1 and DpdA2) (*7*). Of note, DpdA homologs have not been identified in some of the phages that contain modified 7-dezaguanine derivatives (*6, 7*).

PreQ_0_, the key intermediate in all experimentally validated pathways is synthesized from guanosine triphosphate (GTP) by a pathway involving four proteins (FolE, QueD, QueE and QueC, see Figure 1A) found in archaea, bacteria, and some phages (*3, 7*). The pathways then diverge, producing various modifications. PreQ_0_ is reduced by QueF into preQ_1_ in bacteria through a NADPH dependent reaction (*18*). QueF proteins can be categorized into two subgroups. Members of the unimodular subgroup harbor the NADPH binding site and the catalytic residues on the same domain. Members of the bimodular subgroup contain two repeating domains: the N-terminal domain with the NADPH binding site, and the C-terminal domain with the catalytic residues (*19*). PreQ_1_ is inserted in tRNA by the bacterial tRNA transglycosylase bTGT (*20*) and further modified in two steps to produce Q (*3*). PreQ_0_ is directly inserted in tRNA in archaea by arcTGT, where it is further modified into G^+^. The distant TGT paralog, ArcS (*21*), as well as Gat-QueC, a fusion protein of QueC and a glutamine amidotransferase (*22*), and QueF-L, a paralog of the unimodular QueF that lacks the NADPH-dependent reduction activity (*22, 23*), have been found as interchangeable proteins for this reaction.

**Fig. 1.**
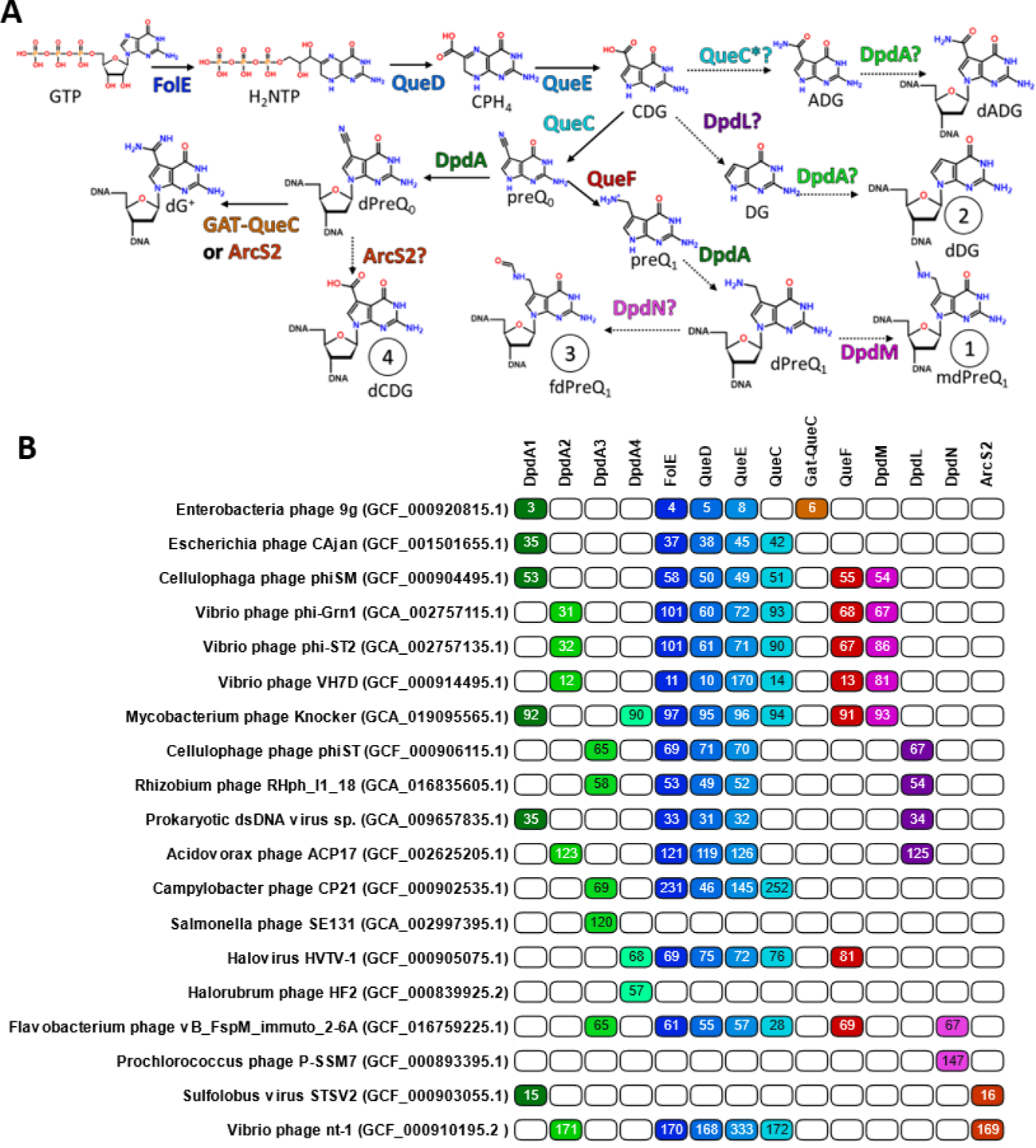
Proteins involved in the 7-deazaguanine DNA modifications proteins. (A) proposed pathway for the biosynthesis of all eight 7-deazaguanine derivatives in DNA and (B) detection of the gene encoding for these proteins in phage genomes (full table in sup. table). Numbering in the table correspond to the gene numbering in each phage. Coloring match with the proteins. In shades of green are the DpdA1 to 4 (7-deazaguanine derivative DNA transglycosidase). In shade of blues, the enzyme leading to preQ_0_: FolE (GTP cyclohydrolase I, EC 3.5.4.16), QueD (CPH_4_ synthase, EC 4.1.2.50), QueE (CDG synthase, EC 4.3.99.3), QueC (preQ_0_ synthase, EC 6.3.4.20), QueC* (proposed ADG synthase, homologue of QueC). In shades of orange, protein modifying preQ_0_ further: QueF (NADPH-dependent 7-cyano-7-deazaguanine reductase, EC 1.7.1.13), ArcS2 (core domain of the archaeosine synthase, PF5591), Gat-QueC, (glutamine amido-transferase class-II domain fused to QueC). In shades of purple, protein discovered and described in this study: DpdL (proposed CDG decarboxylase), DpdM (proposed preQ_1_ or dPreQ_1_ methylase), DpdN (proposed preQ_1_ or dPreQ_1_ formylase). Question marks are proposed reactions that have not be proven. Dashed arrows are previously unpublished reactions. Molecule abbreviations: guanosine tri-phosphate (GTP), dihydroneopterin triphosphate (H_2_NTP), 6-carboxy-5,6,7,8-tetrahydropterin (CPH_4_), 7-carboxy-7-deazaguanine (CDG), 7-amido-7-deazaguanine (ADG), 7-deazaguanine (DG), 7-cyano-7-deazaguanine (preQ_0_), 7-aminomethyl-7-deazaguanine (preQ_1_), 2’-deoxy-7-carboxy-7-deazaguanine (dCDG), 2’-deoxy-7-amido-7-deazaguanine (dADG), 2’-deoxy-7-deazaguanine (dDG), 2’-deoxy-7-cyano-7-deazaguanine (dPreQ_0_), 2’-deoxy-7-aminomethyl-7-deazaguanine (dPreQ_1_), 2’-deoxy-7-(aminomethyl)methyl-7-deazaguanine (mdPreQ_1_), 2’-deoxy-7-(aminoformyl)methyl-7-deazaguanine (fdPreQ_1_). Molecules 1 through 4 (circled numbers) were discovered in this study.

G^+^ was the first 7-deazaguanine derivative found in phages, replacing 25 % of the Gs in the dsDNA genome of *Enterobacteria* phage 9g. The enzymes FolE, QueD, QueE and Gat-QueC are all encoded by this phage, as is DpdA1 (*4*), which inserts preQ_0_ into DNA (*7*). A related phage, *Escherichia* phage CAjan, that encodes QueC rather than Gat-QueC, was found to replace 32% of its Gs with preQ_0_ (*7*). Furthermore, two other 7-deazaguanine derivatives were discovered in phage genomes: 7-amido-7-deazaguanine (ADG), that modifies *Campylobacter* phage CP220 DNA at 100 % (*6*), and preQ_1_ that modifies 30 % of the guanines in Halovirus HVTV-1, a virus that encodes a QueF (*7*). No DpdA was previously detected in these two last viruses. In bacteria, bDpdA, in complex with DpdB, inserts preQ_0_ into DNA, that is further modified into ADG by DpdC (*5*).

In our bioinformatics effort to expand the set of phages that contain 7-deazaguanine derivatives in their genome, we identified four unique 7-deazaguanine derivatives not previously observed in DNA: 7-deazaguanine (DG), 7-(methylamino)methyl-7-deazaguanine (mpreQ_1_), 7-(formylamino)methyl-7-deazaguanine (fpreQ_1_) and 7-carboxy-7-deazaguanine (CDG). We predicted and validated a preQ_1_ methyltransferase enzyme and predicted the involvement of five additional proteins in the synthesis of these modifications, including two additional subfamilies of DpdA, DpdA3 and DpdA4, and three enzymes with unprecedented chemistry.

## MATERIAL AND METHODS

### Strains and plasmids

All strains and plasmids used in this study are referenced in Table S1 and S2, respectively.

### Mass spectrometry analysis

DNA analysis followed our previous publication (*7*). Purified DNA (10 μg) was hydrolyzed in 10 mM Tris-HCl (pH 7.9) with 1 mM MgCl_2_ with Benzonase (20U), DNase I (4U), calf intestine phosphatase (17U) and phosphodiesterase (0.2U) for 16 h at ambient temperature. Following passage through a 10 kDa filter to remove proteins, the filtrate was analyzed by liquid chromatography-coupled triple quadrupole mass spectrometry (LC-MS/MS).

Quantification of the modified 2’-deoxynucleosides (dADG, dQ, dPreQ_0_, dPreQ_1_, mdPreQ_1_, dG^+^, dCDG and m6dA) and the four canonical deoxyribonucleosides (dA, dT, dG, and dC) was achieved by liquid chromatography-coupled triple quadrupole mass spectrometry (LC-MS/MS) and liquid chromatography-coupled diode array detector (LC-DAD), respectively. Aliquots of hydrolysed DNA were injected onto a Phenomenex Luna Omega Polar C18 column (2.1 x 100 mm, 1.6 μm particle size) equilibrated with 98% solvent A (0.1 % v/v formic acid in water) and 2 % solvent B (0.1 % v/v formic acid in acetonitrile) at a flow rate of 0.25 mL/min and eluted with the following solvent gradient: 2-12 % B in 10 min; 12-2 % B in 1 min; hold at 2 % B for 5 min. The HPLC column was coupled to an Agilent 1290 Infinity DAD and an Agilent 6490 triple quadruple mass spectrometer (Agilent, Santa Clara, CA). The column was kept at 40 °C and the auto-sampler was cooled at 4°C. The UV wavelength of the DAD was set at 260 nm and the electrospray ionization of the mass spectrometer was performed in positive ion mode with the following source parameters: drying gas temperature 200 °C with a flow of 14 L/min, nebulizer gas pressure 30 psi, sheath gas temperature 400 °C with a flow of 11 L/min, capillary voltage 3,000 V and nozzle voltage 500 V. Compounds were quantified in multiple reaction monitoring (MRM) mode with the following transitions: *m/z* 310.1è194.1, 310.1è177.1, 310.1è293.1 for dADG; *m/z* 394.1è163.1, 394.1è146.1, 394.1è121.1 for dQ; *m/z* 292.1è176.1, 176.1è159.1, 176.1è52.1 for dPreQ_0_; *m/z* 296.1è163.1, 296.1è121.1, 296.1è279.1 for dPreQ_1_; *m/z* 310.1è163.1, 310.1è121.1 for mdPreQ_1_; *m/z* 309.1è193.1, 309.1è176.1, 309.1è159.1 for dG^+^; *m/z m/z* 311.1è177.1, 311.1è78.9 for dCDG and 266.1è150.1, 266.1è108.1, 266.1è55.1 for m^6^dA. External calibration curves were used for the quantification of the modified 2’-deoxynucleosides and the four canonical deoxyribonucleosides. The calibration curves were constructed from replicate measurements of six concentrations of each standard. A linear regression with r^2^ > 0.995 was obtained in all relevant ranges. The limit of detection (LOD), defined by a signal-to-noise ratio (S/N) of 3, ranged from 0.1 to 1 fmol for the modified 2’-deoxynucleosides. Data acquisition and processing were performed using MassHunter software (Agilent, Santa Clara, CA).

Unknown DNA modification analysis was performed using Agilent 1290 ultrahigh pressure liquid chromatography system equipped with DAD and 6550 QTOF mass detector managed by a MassHunter workstation. The column used for the separation was a Waters ACQUITY HSS T3 column (2.1-150 mm, 1.8 μm). The oven temperature was set at 45 °C. The gradient elution involved a mobile phase consisting of (A) 0.1 % formic acid in water and (B) 0.1 % formic acid in acetonitrile. The initial condition was set at 2 % B. A 25 min linear gradient to 7 % B was applied, followed by a 15 min gradient to 100 % B which was held for 5 min, then returned to starting conditions over 0.1 min. Flow rate was set at 0.3 ml/min, and 2 μL of samples was injected. The electrospray ionization mass spectra were acquired in positive ion mode. Mass data were collected between m/z 100 and 1000 Da at a rate of two scans per second. The electrospray ionization of the mass spectrometer was performed in positive ion mode with the following source parameters: drying gas temperature 250 °C with a flow of 14 L/min, nebulizer gas pressure 40 psi, sheath gas temperature 350 °C with a flow of 11 L/min, capillary voltage 3,500 V and nozzle voltage 500 V. Two reference masses were continuously infused to the system to allow constant mass correction during the run: *m/z* 121.0509 (C_5_H_4_N_4_) and *m/z* 922.0098 (C_18_H_18_O_6_N_3_P_3_F_24_). Raw spectrometric data were analyzed by MassHunter Qualitative Analysis software (Agilent Technologies, US).

### Protein sequence detection in phages

HHpred online tool (https://toolkit.tuebingen.mpg.de/tools/hhpred) (*24, 25*) was used with default setting against the pfam database (Pfam-A_v35) (*26*) to investigate the deduced proteins encoded by genes flanking the 7-deazaguanine modification genes in *Cellulophaga* phage phiSM, *Cellulophaga* phage phiST, and Halovirus HVTV-1. DpdL, DpdM, DpdA3 and DpdA4 were predicted this way. DpdN was discovered by looking at the annotations of the genes in the vicinity of the 7-deazaguanine modification genes in *Flavobacterium* phage vB_FspM_immuto_2-6A. These proteins were then used as queries to retrieve homologs in the proteome of viruses publicly available in NCBI GenBank database (July 2022) using psiBLAST version 2.13.0 (*27*), with at most three iterations. Other previously discovered proteins involved in the 7-deazaguanine derivative DNA modifications (Data S1) (*7*) were used to identify homologs in viral genomes encoding for at least one of DpdL, DpdM, DpdN, DpdA3, or DpdA4 using BLASTp version 2.13.0 (*28*). HHpred and expert annotation were used to sort these proteins and curate false positives. All protein matches are summarized in Data S1.

### Alignments, trees and structures

Protein sequences were collected from the NCBI database, using the protein id collected from the detection. Multiple sequence alignments were generated using MAFFT (*29*) online server (version 7, https://mafft.cbrc.jp/alignment/server/), with default settings and then visualized using Jalview version 2.11.2.4. Clustering trees were generated using Graph Splitting (*30*) online server (version 2.0, http://gs.bs.s.u-tokyo.ac.jp/), with default settings. Protein structures were predicted using the multimer collab notebook of AlphaFold2 (version 2.2.4 (*31*), https://colab.research.google.com/github/deepmind/alphafold/blob/main/notebooks/AlphaFold.ipynb). Protein structures were visualized using ChimeraX version: 1.5rc202210241843 (*32*), and already published protein structure were imported from PDB (https://www.rcsb.org, (*33*)). Autodock Vina was used to predict the docking of chemicals in enzymes.

## RESULTS

### 7-(methylamino)-methyl-7-deazaguanine in Cellulophaga phage phiSM DNA

Cellulophaga phage phiSM encodes a complete set of dPreQ_1_ synthesis genes, including DpdA, FolE, QueD, QueE, QueC, and QueF (Figure 1, Data S1) and thus should harbor preQ_1_ in its genome, as previously observed for Halovirus HVTV-1 (*7*). To test this hypothesis, we used liquid chromatography coupled to diode array UV detection and a tandem mass spectrometer (LC-UV-MS/MS) to analyze of the nucleosides obtained from enzymatic digestion of phiSM genomic DNA, as we previously described (*4, 7*). A 2’-deoxynucleoside form of preQ_1_ (dPreQ_1_) was indeed detected (3,790 modifications per 10^6^ nucleotides, ∼ 1.1 % of the Gs, Table 1). In addition to the UV peaks for the four canonical nucleosides, dA, dC, dT and dG, an unknown UV peak with a mass of 310 Da was observed at a retention time of 6.5 min (Figure 2A, 212 modifications per 10^6^ nucleotides, ∼ 0.1 % of the Gs, Table 1). The collision-induced dissociation (CID) MS/MS spectra of the unknown peak revealed the protonated 2’-deoxyribose ion (*m/z* 117) and its further dehydration ions (*m/z* 99 and 81), confirming that the unknown peak corresponded to a non-canonical nucleoside (Figure 2B). The CID MS/MS spectra of dPreQ_1_ and the unknown modification showed very similar patterns. Both compounds showed fragment [M+H]^+^ ions at *m/z* 163 and *m/z* 279, indicating that the unknown modification could be a derivative of dPreQ_1_ (Figure 2B). The mass of the unknown modification is 14 Da greater than that of dPreQ_1_, implying that it is a methylated product of dPreQ_1_. The high-resolution mass spectrometry (HRMS) of the protonated [M+H]^+^ ion (*m/z* 310.1511) of the unknown modification matches the theoretical mass of protonated methylated-dPreQ_1_ very well (*m/z* 310.1515, mass error = 1.29 ppm). The MS/MS spectra of the unknown modification revealed a fragment ion with a loss of 31 Da (*m/z* 310 → *m/z* 279), corresponding to a methylamino group (Figure 2B). The loss of the methylamino group was observed in the MS/MS spectra at low CID energy, indicating that the methyl group is likely linked to the 7-amino group, which is less stable than the linkage to the 2-amino group in CID MS/MS experiment.

**Fig. 2.**
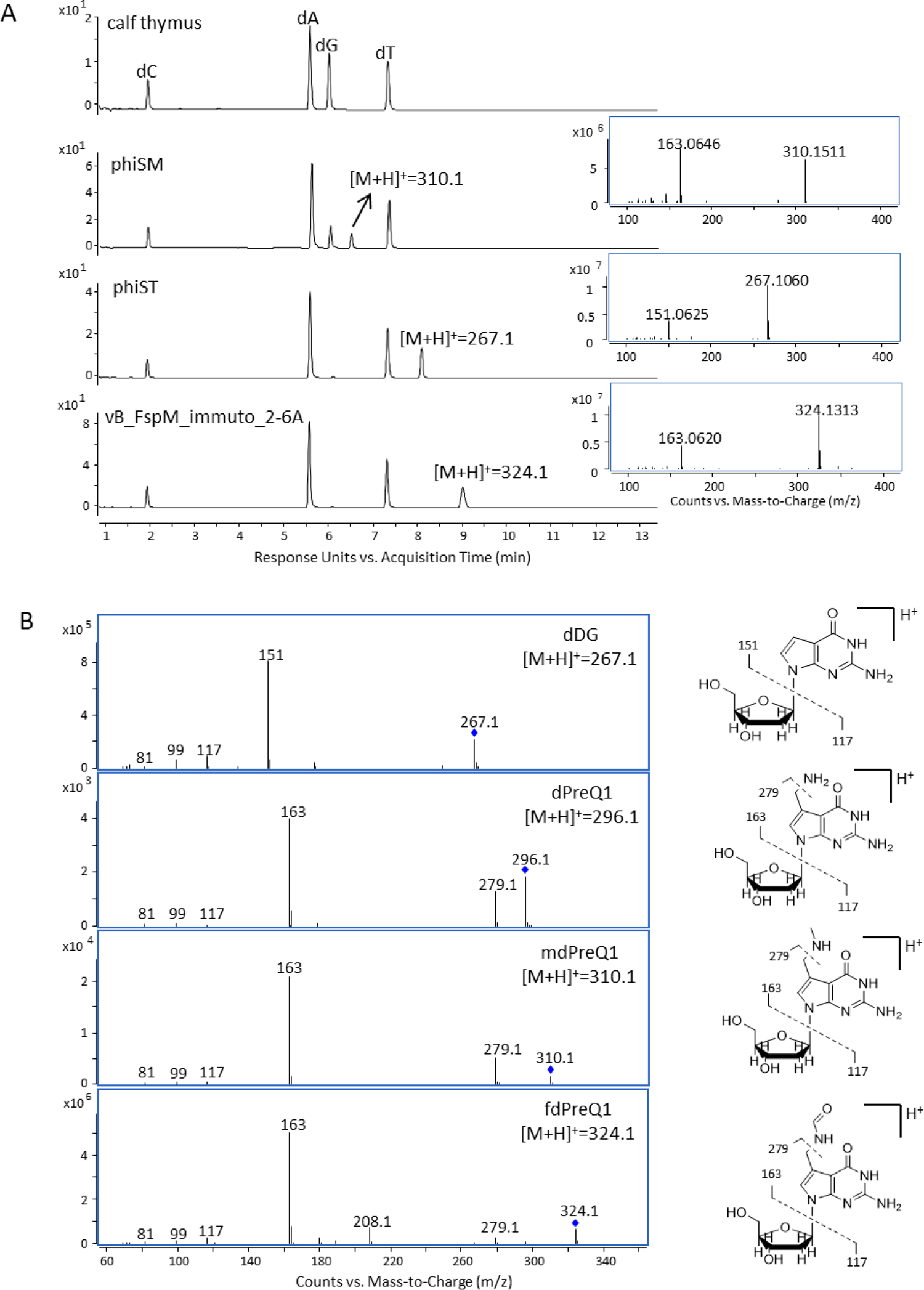
HPLC-UV-MS analysis of digested genomic DNA samples from bacteriaophage phiSM, phiST and vB_FspM_immuto_2-6A. (A) The HPLC-UV chromatogram on top was obtained from calf thymus DNA to show the retention of the canonical nucleosides. PhiSM shows a fifth peak with a mass of 310 Da. The dG peak disappeared, and a new peak was detected in phiST and vB_FspM_immuto_2-6A with a mass of 267 Da and 324 Da, respectively. (B) The MS/MS spectra and proposed CID fragmentation of dDG ([M+H]^+^=267.1), dpreQ1 ([M+H]^+^=296.1), mpreQ1 ([M+H]^+^=310.1), fdpreQ1 ([M+H]^+^=324.1). Molecule abbreviations: dC, 2’-deoxycytidine; dA, 2’-deoxyadenosine; dG, 2’-deoxyguanosine; dT, 2’-deoxythymidine.

**Table 1.** Quantification of 7-deazaguanine derivate DNA modifications per 10^6^ nucleotides in viruses.

| virus | dCDG | dADG | dDG | dPreQ<sub>0</sub> | dPreQ<sub>1</sub> | mdPreQ<sub>1</sub> | fdPreQ<sub>1</sub> | dG<sup>+</sup> |
| --- | --- | --- | --- | --- | --- | --- | --- | --- |
| Cellulophaga phage phiSM | 0 | 0 | 0 | 0 | 3790 | 212 | 0 | 0 |
| Vibrio phage phi-Grn1 | 0 | 0 | 0 | 0 | 816 | 35 | 0 | 0 |
| Vibrio phage phi-ST2 | 0 | 0 | 0 | 0 | 668 | 44 | 0 | 0 |
| Cellulophaga phage phiST | 0 | 0 | 302,000 | 0 | 0 | 0 | 0 | 0 |
| Flavobacterium phage<br>vB_FspM_immuto_2-6A | 0 | 0 | 0 | 0 | 0 | 0 | 345,000 | 0 |
| Sulpholobus virus SVST-2 | 149 | 0 | 0 | 0 | 0 | 0 | 0 | 0 |

We chemically synthesized 7-(methylamino)methyl-2’-deoxy-7-deazaguanine (mdPreQ_1_, Scheme 1), which was purified by HPLC and characterized by NMR and HRMS, to test whether methylation at the 7-amino position of dPreQ_1_ corresponds to the unknown DNA modification. The standard was then analyzed using LC-UV-MS/MS. The retention time and MS/MS spectra of mdPreQ_1_ standard were identical to those of the unknown non-canonical nucleoside (Figure S1), confirming that the unknown modification is mdPreQ_1_. The same modification was identified in Cellulophaga phages phi38:2 and phi47:1 (Figure S2), which are related to phage phiSM (*34*).

### Phage and bacterial QueF have similar function

Given the presence of dPreQ_1_ in DNA (*7*) and its subsequent modification to mdPreQ_1_, one must ask which precursor molecule (preQ_0_ or preQ_1_) is directly inserted in the genome. Indeed, to date, all characterized DpdAs insert preQ_0_ into DNA (*5, 7*). Hence, the phage QueF should behave like the Archaeal QueF-L (*22*) and generate preQ_1_ from preQ_0_ inserted into DNA. However, if phage QueF proteins are similar to the bacterial QueF (*35*) and form preQ_1_ base from preQ_0_ base, then the DpdA of these phages should have changed in substrate specificity to insert preQ_1_ in DNA. QueF family sequences were collected from phages (Data S1 and (*7*)) and compared to the sequences of three experimentally validated QueF proteins: the bimodular QueF of *Escherichia coli* (NP_417274), the unimodular QueF of *Bacillus subtilis* (NP_389258) and the QueF-L of *Pyrobaculum calidifontis* (WP_011848915). Surprisingly, no phage QueF sequences aligned with QueF-L (Alignment S1). The bimodular sequence aligned with half of the phage QueF sequences (Figure S3A and B), while the unimodular one aligned with the other half (Figure S3B). The NADPH binding motif, E(S/L)K(S/A)hK(L/Y)(Y/F/W), and most of the amino acids characteristic of the QueF family sequences (Figure S3B, stared conserved residues (*19, 36*)) were conserved in all phage sequences with the exception of a tyrosine (Y221 in *E. coli*, Y87 in *B. subtilis*, Y52 in *P. calidifontis*) in the unimodular phage sequences. This degree of conservation strongly suggests that the phage QueF proteins are NADPH-dependent preQ_0_ reductases.

To validate this prediction, an *E. coli ΔqueF* mutant was transformed with plasmids expressing *queF* genes from three phages/viruses, namely Cellulophaga phage phiSM, Vibrio phage VH7D, and Halovirus HVTV-1. We observed that expression of the phiSM and VH7D *queF* genes, but not of the HVTV-1 one, complemented the Δ*queF* strain’s Q-deficiency phenotype (Figure S3C). Because HVTV-1 is a virus infecting a hyper-saline archaeon, *Haloarcula valismortis*, expressing its *queF* gene in *E. coli* in a low salt environment may have been challenging. Nonetheless, these experiments confirmed that phage QueF, like its bacterial counterpart, catalyzed the reduction of preQ_0_ to preQ_1_.

To confirm that phage DpdA encoded in QueF-like reductase switched specificity to preQ_1_, we cloned phiSM *dpdA1* and VH7D *dpdA2* in pBAD24 vector and expressed them in several mutants of *E. coli*. In our experiments, phiSM DpdA1 was found to be inactive, while VH7D DpdA2 inserted preQ_1_ in DNA (2,765 modifications per 10^6^ nucleotides), proving that this DpdA substrate specificity indeed adapted to preQ_1_. Interestingly, VH7D DpdA2 also inserted preQ_0_ at a lower efficiency (712 modifications per 10^6^ nucleotides) in a strain that does not produce preQ_1_ and accumulates preQ_0_ (Δ*queF*, see pathway in Figure 1A), as well as CDG at a very low efficiency (67 modifications per 10^6^ nucleotides) in a strain that accumulates CDG (Δ*queC*, see Figure 1A).

### Prediction and validation of a preQ_1_ methyltransferase

Phages that harbor the mdPreQ_1_ modification should encode a methyltransferase that appends a methyl group onto the nitrogen of the methylamino group of preQ_1_ in genomic DNA. There are four genes coding for proteins of unknown function in the cluster of genes encoding the preQ_1_ pathway in phage phiSM, namely CEPG_00048, CEPG_00054, CEPG_00056 and CEPG_00057. The proteins CEPG_00048 and CEPG_00057 were ruled out as candidates because they encode short proteins (∼ 60 amino acids) and are not found by psiBLAST in other phages encoding a deazaguanine DNA modification pathway (Data S2 and S3). CEPG_00056 homologs were observed in closely related *Cellulophaga* phages and in eukaryotic herpes viruses (Data S4). This candidate was eliminated because no deazaguanine DNA modification was ever found in eukaryotic viruses (*7*) and eukaryotes do not produce any preQ_1_ (*3*). Finally, CEPG_00054 homologs were found in seven other phages, including Vibrio phages phi-Grn1, phi-ST2, and VH7D, which were predicted to encode preQ_1_ modification pathways (Figure 1B, Data S1 and S5). This protein belongs to the DUF3109 family (Data S6) and has an *E. coli* homolog, YkgJ, which is annotated as a zinc or iron binding protein, making CEPG_00054 the leading candidate for the missing preQ_1_ methyltransferase, and tentatively renaming it DpdM.

We found that the genome of Vibrio phages phi-Grn1 and phi-ST2 DNA, encoding DpdM homologs (Figure 1B, Data S1), were also modified with mdPreQ1 (Figure S4, Table 1) at a rate of 0.01 % of the Gs for both phages (35 and 44 modifications per 10^6^ nucleotides, respectively, Table 1). Finally, expressing the predicted VH7D *dpdM* gene in an *E. coli* strain already expressing the *dpdA2* of Vibrio phage VH7D resulted in the formation of low levels of mdPreQ1 in plasmid DNA (Table 2). Taken altogether, these data linked mdPreQ1 with the presence of DpdM (Figure 1B, Data S1).

**Table 2.** Quantification of 7-deazaguanine derivate DNA modifications per 10^6^ nucleotides in plasmids encoding for Vibrio phage VH7D system expressed in *E. coli* strains.

| background | pBAD24 | pBAD33 | dCDG | dPreQ<sub>0</sub> | dPreQ<sub>1</sub> | mdPreQ<sub>1</sub> |
| --- | --- | --- | --- | --- | --- | --- |
| WT | empty | empty | 0 | 0 | 0 | 0 |
|  | DpdA2 | empty | 0 | 7 | 2765 | 0 |
|  | empty | DpdM | 0 | 0 | 0 | 0 |
|  | DpdA2 | DpdM | 0 | 0 | 0 | 9 |
|  | DpdM | empty | 0 | 0 | 0 | 0 |
|  | empty | DpdA2 | 0 | 9 | 3666 | 0 |
|  | DpdM | DpdA2 | 0 | 7 | 97 | 40 |
| $\Delta queF$ | DpdA2 | empty | 0 | 630 | 0 | 0 |
|  | DpdA2 | DpdM | 0 | 696 | 0 | 0 |
|  | empty | DpdA2 | 0 | 616 | 0 | 0 |
|  | DpdM | DpdA2 | 0 | 672 | 0 | 0 |
| $\Delta queC$ | DpdA2 | empty | 82 | 0 | 0 | 0 |
|  | DpdA2 | DpdM | 52 | 0 | 0 | 0 |
|  | empty | DpdA2 | 95 | 0 | 0 | 0 |
|  | DpdM | DpdA2 | 54 | 0 | 0 | 0 |

As shown above with VH7D *dpdA2* expression alone, when both the VH7D *dpdA2* and *dpdM* genes were expressed in a Δ*queF* background, which does not produce preQ_1_ but accumulates preQ_0_, preQ_0_ was inserted into bacterial DNA at a ∼ 5-time lower efficiency. Similarly, when a Δ*queC* background that accumulates CDG was used, CDG was found in DNA with a ∼ 40-fold decrease in efficiency (Table 2).

### DpdM proteins likely bind two metals

Although the initial amino acid sequence analysis of DpdM from Cellulophaga phage phiSM revealed a CxxxCxxCC metal binding motif (Data S6), this motif was missing in the Vibrio phage phi-ST2 homolog. We found that the *orf* encoding this protein was miscalled and discovered that by selecting a start codon 171 nucleotides prior to the originally predicted one now resulted in a polypeptide containing the CxxxCxxCC motif (phi-ST2 corrected in Alignment S2).

The tertiary and quaternary structures of DpdM from both *Cellulophaga* phage phiSM and *Vibrio* phage VH7D were predicted using AlphaFold2. Both proteins were predicted to be monomeric, with only a few amino acids interacting between monomers (data not shown). The phiSM DpdM prediction (Figure S5A) had a higher confidence score than the VH7D prediction (Figure S5B). We found small domains around the VH7D predicted structure with unknown function as shown in the alignment. However, the core parts of the protein were well aligned (Figure S5C).

The phiSM DpdM structure contains a tunnel that has an electro-positively charged groove on one side (Figure 3A), which could be a candidate site for DNA binding, and a second groove on the opposite side (Figure 3B), which could be a site for a methyl donor binding. Surprisingly, majority of the conserved residues are clustered around this tunnel (Figures 3C and D). The CxxxCxxCC motif appears to be divided into two metal binding sites rather than one. The CxxxCxxC motif (orange in Figure S5H; representing C33, C38, and C41), is a known motif for a Fe_4_S_4_ cluster and SAM binding (*37*), but the presence of a fourth cysteine, C150 (red in Figure 3E), in the pocket would disrupt the Fe_4_S_4_ binding and may bind another metal instead, as well as a different methyl donor. It appears that the fourth cysteine in the CxxxCxxCC motif is involved in another metal binding pocket containing three other cysteine residues (yellow in Figure 3F; representing C42, C92, C102, and C112). Both these metal binding pockets are found in the DpdM tunnel implying that they both participate in the transfer of the methyl group from the methyl donor to preQ_1_ in DNA.

**Fig. 3.**
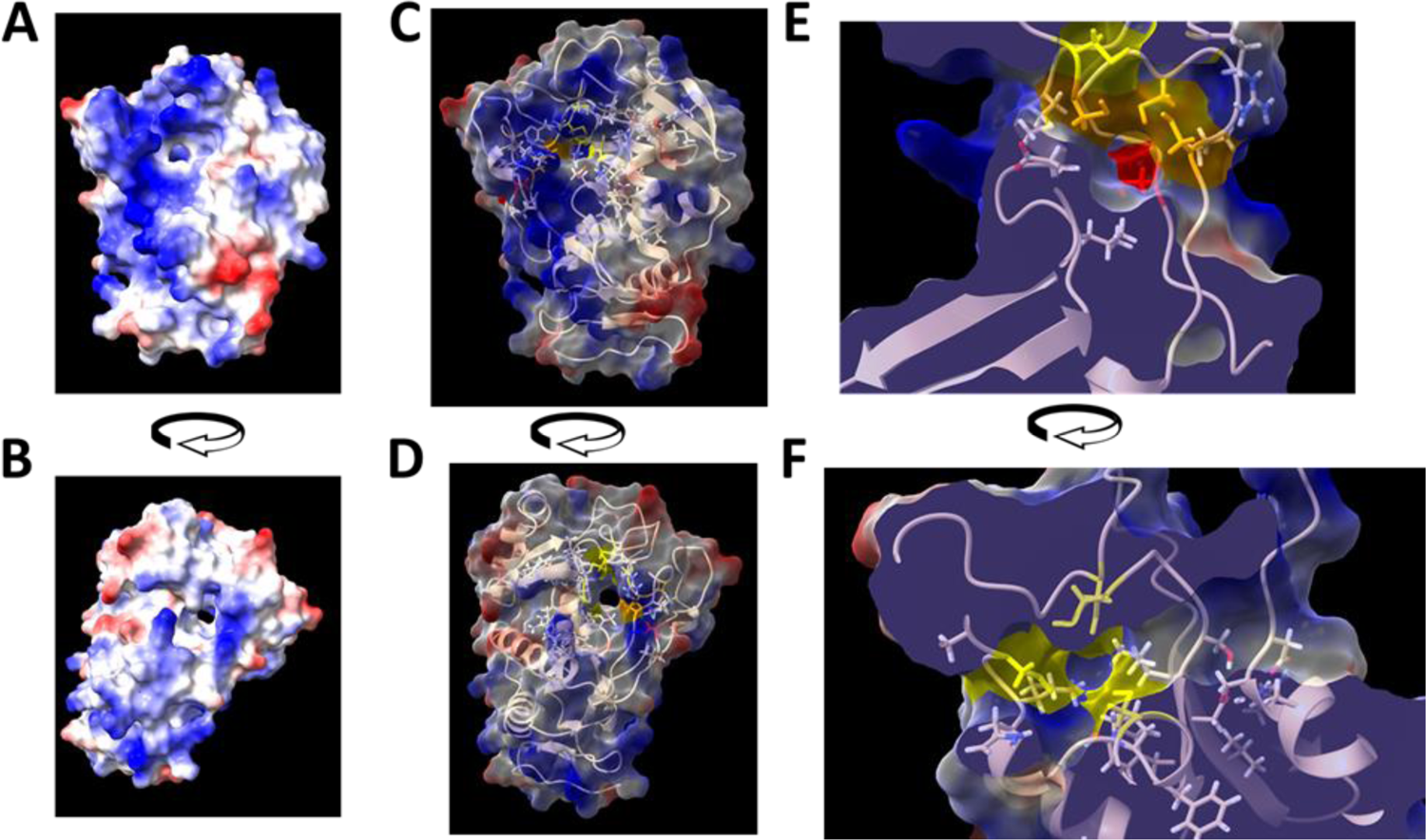
Analysis of Cellulophaga phage phiSM predicted structure. Electromagnetic surface charges of both proteins were visualized on two opposite faces of the proteins (A and B), blue are positive charges and red negative charges. Conserved residues were added to the structures with a 60 % transparency on the electromagnetic surface (C and D). Conserved cysteines predicted to bind metal are colored in yellow, orange, and red as described in the text. Visualization of these cysteines were zoomed in to view both metal pockets (E and F).

The tunnel observed in phiSM DpdM, the positively charged groove (Figure S3D), and the binding site on the opposite side of the protein (Figure S5E), are absent in the VH7D DpdM structure. If the conserved residues in both VH7D and phiSM DpdM proteins are mostly clustered at the same place (Figure S5F and G), the two metal binding sites are located in one side of the enzyme in VH7D DpdM (Figure S5H and I; in orange C45, C50, C52; in red C222; in yellow C53, C120, C129, C146).

### Discovery of 7-deazaguanine in *Cellulophaga* phage phiST DNA

Cellulophaga phage phiST encodes FolE, QueD and QueE but not QueC or DpdA (Figure 1, Data S1) (*7*). The product from the reaction catalyzed by QueE, 7-carboxyl-7-deazaguanine (CDG; see Figure 1A) (*3*) was not detected in this phage DNA using LC-UV-MS/MS analysis. Meanwhile, peaks corresponding to three of the canonical nucleosides, dA, dC, and dT were observed, along with an unknown peak with a mass of 267 Da and retention time of 8.2 min (Figure 2A). The CID MS/MS spectra of the unknown peak revealed the protonated nucleobase [B+H]^+^ ion at *m/z* 151, corresponding to the glycosidic bond cleavage with the loss of a neutral 2’-deoxyribose (*m/z* 116). The presence of a protonated 2’-deoxyribose ion (*m/z* 117) and its dehydration ions (*m/z* 99 and 81) confirmed that the unknown peak corresponds to a non-canonical nucleoside (Figure 2B). A signal for dG was not detected, suggesting that it had been completely replaced by the unknown non-canonical nucleoside. Although CDG was not detected, the unknown modification could have been a CDG derivative. The mass of the unknown modification is 1 Da less than dG and the HRMS of its protonated [M+H]^+^ ion (*m/z* 267.1094) matches well with the theoretical mass of the protonated [M+H]^+^ ion of the decarboxylated derivative of dCDG, 2’-deoxy-7-deazagunine (dDG, *m/z* 267.1093, mass error = 0.37 ppm). We analyzed a synthetic dDG standard by LC-UV-MS/MS and found that its retention time and CID MS/MS spectra matched those of the unknown non-canonical nucleoside (Figure S6), confirming that the unknown modification was dDG. The same modification was found in Cellulophaga phages phi19:2 and phi13:1 (Figure S7), which are related to phage phiST (*34*).

### Prediction of a decarboxylase leading to 7-deazaguanine

The discovery of dDG suggested that phiST encodes a CDG decarboxylase that could remove the carboxyl moiety of CDG to form DG. Between the CDG pathway and the polymerase genes of phiST lie five genes coding for protein of unknown function: CGPG_00064, CGPG_00065, CGPG_00066, CGPG_00067 and CGPG_00068. Other phages containing 7-deazaguanine modifications pathway do not encode CGPG_00064 and CGPG_00066 homologs (Data S7 and S8). CGPG_00068 encodes a dUTPase (99.15% probability matching to PF08761.14 by HHpred, Supplementary Data S9) or MazG (98.4% probability matching to PF12643.10 by HHpred, Data S9), which has been shown to hydrolyze dNTP in phages (*38*). CGPG_00065 is a distant homolog of a TGT/DpdA (98.83% probability matching to PF01702.21 by HHpred, Data S10) and its function is discussed in the sections below. CGPG_00067 is highly similar to QueD (99.89% probability matching to PF01242.22 by HHpred, Data S11). T-fold enzymes like QueD bind pterins or purines (*39*), and three of them are involved in preQ_0_ synthesis (*3*). This gene was also found in other phages encoding for a DpdA, FolE, QueE, and QueD but not QueC (Figure 1B; Data S1 and S12). Because of these findings, CGPG_00067 was chosen as the best candidate for the missing CDG decarboxylase and renamed DpdL.

To investigate structural differences between QueD and DpdL, we aligned the sequences of QueD from *E. coli* (NP_417245.1) and *B. subtilis* (NP_389256.1) with all the proposed decarboxylase phage protein sequences (Alignment S3). Both proteins share three histidines and two glutamic acids, but the position of the fourth histidine differs in the multiple alignment. The signature motif of QueD CxxxHGH (*40*) is also changed to LxxxHRHxF in DpdL. Both histidines of the motif coordinate the zinc ion in the active site, and the cysteine is required for the catalyzation of the reaction. Because the glycine residue is not involved in ligand binding or catalysis, changing it to arginine would not change any essential properties of the active site. The conversion of cysteine to leucine does, as QueD is inactive without this cysteine (*41*). The predicted structure of DpdM indicated that it would catalyze the reaction on the base (Supplementary Text, Figure S8) via an alkaline decarboxylation mechanism involving zinc or other bound metal. This would imply that the specificity of the co-encoded DpdA would be changed from preQ_0_ to DG.

We expressed *dpdL* genes from phage phiST and *Acidovorax* phage ACP17 in *E. coli* alongside their respective *dpdA* genes, but we were unable to detect any dDG in this heterologous system (data not shown). Proteins may be inactive in *E. coli* due to temperature, salt, or codon optimization differences with their host organisms, or other unknown enzymes may be required to complete the reaction.

### A DpdA is encoded in all phages that harbor 7-deazaguanine derivatives

As previously stated, CGPG_00065 is a distant homolog of a TGT/DpdA and is also found in Campylobacter phages (Figure 1B; Data S1 and S13), which have been previously shown to be modified by ADG (*6*). This DpdA3 family had not previously been identified (*6*) and is the most logical candidate for the enzyme inserting a 7-deazaguanine derivate in the DNA of both phiST and *Campylobacter* phages (*6*).

It is difficult to predict the substrate specificity of the DpdA3 family (Figure 1A). DpdA3 is unlikely to insert preQ_0_ as the full pathways are absent in phiST and the Campylobacter phages stop the synthesis at CDG (*6*). As a result, DpdA3 may insert CDG, a common precursor of dADG and dDG. Because the nucleoside form of ADG was detected in the cytoplasm of *Campylobacter jejuni* infected with phage CP220 (*6*), the DpdA3 might have shifted their substrate specificity to insert DG or ADG.

With the discovery of the DpdA3 subfamily, only a few of the phages/viruses identified in our previous study remained with no encoded DpdA (*7*). We reanalyzed the genome of Halovirus HVTV-1, which is modified with preQ_1_. HVTV1_69 gene product had a 100 % probability of matching with PF20314.1, a domain of unknown function (DUF6610), by HHpred, but also 92.5 % with PF01702.21, a tRNA-guanine transglycosylase (Data S14). Furthermore, homologs of this protein were found to be encoded in other archaeal viruses that also contain preQ_1_ synthesis genes, as well as a singleton modification gene in a few other viral genomes, including Halorubrum phage HF2 (Figure 1B; Data S1 and S15). With the discovery of this final DpdA subgroup, renamed DpdA4, all phages known to harbor a 7-deazaguanine in their DNA encode a DpdA family protein, which now could be considered a signature protein family for the presence of such DNA modifications.

### 7-(Formylamino)-methyl-7-deazaguanine in Flavobacterium phage vB_FspM_immuto_2-6A DNA

Flavobacterium phage vB_FspM_immuto_2-6A encodes DpdA3, FolE, QueD, QueE, QueC, and QueF (Figure 1, Data S1) and should thus have complete guanosine replacement to preQ_1_. However, dPreQ_1_ was not detected in this phage genome using LC-UV-MS/MS analysis. Meanwhile, peaks corresponding to three of the canonical nucleosides, dA, dC and dT, as well as an unknown peak at a retention time of 9 min with a mass of 324 Da were observed in the LC-UV-MS/MS analysis of this phage DNA (Figure 2A). The CID MS/MS spectra of the unknown peak revealed fragment ions at *m/z* 208, 117, 99, and 81, which could be attributed similarly to the loss of 2’-deoxyribose to form [B+H]^+^ ion and protonated 2’-deoxyribose ion and its further dehydration ions, respectively, confirming the unknown peak is a noncanonical nucleoside (Figure 2B). The dG peak was not detected, indicating that it has been completely replaced by the unknown non-canonical nucleoside. The CID MS/MS spectra of preQ_1_, mdPreQ_1_ and the unknown modification showed very similar pattern, with fragment [M+H]^+^ ions observed at *m/z* 163 and *m/z* 279 for all three compounds, indicating that the unknown modification could also be a dPreQ_1_ derivative (Figure 2B). The unknown modification had a mass of 28 Da greater than dPreQ_1_, corresponding to one additional carbon and one oxygen (formyl group) or two additional carbons and four hydrogens (ethyl or dimethyl group). The unknown modification mass (*m/z* 324.1313) matched well with the theoretical mass of protonated [M+H]^+^ ion of formyl-dPreQ_1_ (*m/z* 324.1308, mass error = 1.56 ppm, Figure S9) but not with the theoretical mass of protonated ethyl- or dimethyl-dPreQ_1_ (*m/z* 324.1672, mass error = 112.32 ppm). The MS/MS spectra of the unknown modification at low CID energy revealed a fragment ion with a loss of 45 Da (*m/z* 324 → *m/z* 279), corresponding to a formylamino group. This suggested that the formyl group was most likely linked to the 7-amino group, which is less stable than the 2-amino group in CID MS/MS experiment. To test our hypothesis, we chemically synthesized fdPreQ_1_, which was then purified using HPLC and characterized using NMR and HRMS. (Scheme 2, Figure S9). The standard was then analyzed using LC-UV-MS/MS. The standard retention time and MS/MS spectra were identical to those of the unknow noncanonical nucleoside, confirming that the unknown modification is fdPreQ_1_ (Figure S10). This finding suggested that the vB_FspM_immuto_2-6A genome may encode a formytransferase that adds a formyl group to dPreQ_1_.

### Prediction of a preQ_1_ formyltransferase

A protein annotated as PF00551 formyltransferase is encoded close to the 7-deazaguanine insertion gene cluster of phage vB_FspM_immuto_2-6A (locus tag KNV73_gp067, Figure 1B, Data S1). This protein was used to identify similar proteins in other viral genomes (Figure 1B; Supplementary Data S1, S16). We found six phages that encode a similar protein and shared the entire pathway from FolE to QueF, including a DpdA, and 15 other phages that encode a similar protein but lacked any 7-deazaguanine modification genes. These sequences could be divided into three groups, according to a multiple sequence alignment (Alignment S4) and a clustering cladogram (Figure S11). One of them include four proteins that are co-encoded with the modification pathway (DpdA and FolE to QueF, Data S1): YP_010114479.1, of phage vB_FspM_immuto_2-6A, as well as CAB5226463.1, CAB4142580.1 and CAB5221950.1, all three encoded by uncultured *Caudoviral* phages and renamed DpdN. The two other groups appear to be unrelated to the fdPreQ_1_ modification because group 2 is encoded by phage that do not encode the proteins involved in the modification pathway and group 3 contains members that are longer forms of the formylase, which are likely to be involved in other reactions (Supplementary Data S1). As DpdN is a member of the same superfamily as the enzyme PurN, which catalyzed the formylation of 5-phospho-ribosyl-glycinamide in the purine synthesis pathway (*42*), it likely uses the same formyl donor 5-methyl-5,6,7,8-tetrahydrofolate (Supplementary Text and Figure S12).

### 7-Carboxy-7-deazaguanine in Sulfolobus virus STSV-2

Because Sulfolobus virus STSV-2 encodes DpdA and ArcS (Figure 1, Supplementary Data S1), it should harbor dG^+^ in its genome. As we previously described (*4, 7*), we used LC-UV-MS/MS to analyze the nucleosides obtained from enzymatic digestion of STSV-2 genomic DNA. dCDG, but not dG^+^, was detected at a rate of 0.04 % of the Gs (149 modifications per 10^6^ nucleotides, Figure S13, Table1).

There was no other neighboring gene that was clearly shared with other phages or viruses (data not shown). Surprisingly, the host archaeon, *Sulfolobus tengchongensis*, does not encode any proteins involved in the Q or G^+^ biosynthesis pathway (data not shown). We believed that its ArcS evolved to revert preQ_0_ into CDG. To investigate this, we aligned STSV-2 ArcS sequence with canonical ArcS proteins (*21*) and with homologs previously identified in other viruses (*7*) (Alignment S5). The phage/virus ArcS corresponds to only the core catalytic domain of the canonical ArcS (PF17884.4 annotated as DUF5591, 99.9 % similar for STSV2_16 encoded by Sulfolobus virus STSV2, Supplementary Data S17, and 99.9 % for VPFG_00169 encoded by *Vibrio* phage nt-1, Data S18). It has previously been demonstrated that the ArcS have a high degree of diversity (*21*). Initially, four domains were identified in ArcS (Nt, C1, C2 and PUA). The PUA domain is specific to RNA binding, the Nt domain is similar to the TGT catalytic domain and the C1 domain is specific to ArcS and contains the catalytic core of the functions. These four domains are found in others, but in some organisms, the Nt domain is separated from the other three domains. In some archaea, the C1 domain is encoded independently, as in the phages. The C1 domain’s specific motif, PC-X3-KPY-X2-S-X2-H (*21*), was conserved in STSV-2 ArcS but slightly degenerated in *Vibrio* phage nt-1 ArcS (Figure S14, Alignment S5).

We decided to test the ArcS of phage nt-1 because the ArcS from a hyperthermophile organism might be inactive in our *E. coli* double plasmid test system, as hypothesized previously (*7*). Both nt-1 *dpdA2* and *arcS* were cloned in pBAD24 and pBAD33 vectors, expressed in *E. coli*, and the plasmids were extracted. We found that dPreQ_0_ is inserted into DNA when nt-1 DpdA2 is expressed alone, and dG^+^ is present when nt-1 ArcS is co-expressed (Table 3). This suggested that STSV-2 ArcS, which is less degenerate than nt-1 ArcS, may have the same function, generating dG^+^. Therefore, additional STSV-2 proteins yet to be identified are required to catalyze the insertion of CDG in DNA in this virus.

**Table 3.** Quantification of 7-deazaguanine derivate DNA modifications per 10^6^ nucleotides in plasmids encoding for Vibrio phage nt-1 system expressed in *E. coli* strains.

| background | pBAD24 | pBAD33 | dPreQ<sub>0</sub> | dG<sup>+</sup> |
| --- | --- | --- | --- | --- |
| WT | empty | empty | 0 | 0 |
|  | DpdA2 | empty | 926 | 0 |
|  | empty | ArcS | 0 | 0 |
|  | DpdA2 | ArcS | 0 | 4 |
|  | ArcS | empty | 0 | 0 |
|  | empty | DpdA2 | 1607 | 0 |
|  | ArcS | DpdA2 | 11 | 7 |
| *ΔqueD* | empty | empty | 0 | 0 |
|  | DpdA2 | empty | 0 | 0 |
|  | empty | ArcS | 0 | 0 |
|  | DpdA2 | ArcS | 0 | 0 |
|  | ArcS | empty | 0 | 0 |
|  | empty | DpdA2 | 0 | 0 |
|  | ArcS | DpdA2 | 0 | 0 |

## DISCUSSION

In this current era of active discoveries of new bacterial defence systems against phages driven by genomic data mining (*12–14*), the identification of phage counter defences (*11*), including DNA modifications, is also rising. This study focused on a group of guanine modifications, known as 7-deazaguanines, where the nitrogen in position 7 of guanine is replaced by a carbon allowing an easier addition of various side chains at this position. In a previous study, we presented four side chains, namely dPreQ_1_ and dG^+^ (*7*) in the genome of some viruses, and dADG and dPreQ_0_ in both phage and bacterial DNA (*4, 7*). Here, we have doubled the number of 7-deazaguanine derivatives identified in DNA, with the description of four new epigenetic marks (a) two modifications that represent further modification of dPreQ_1_, namely mdPreQ_1_ and fdPreQ_1_; (b) one precursor of preQ_0_, namely dCDG; (c) and one unprecedented natural 7-deazaguanine, dDG. In addition, we identified five previously undescribed families of viral enzymes involved in the synthesis of these modified bases (Figure 1).

Phage genomic DNAs encoding QueF homologs always contain dPreQ_1_, or derivatives. Indeed, viral QueF proteins are preQ_0_ reductases (Figure S3), like the bacterial ones (*7*). Two hypermodified dPreQ_1_ that each require an additional enzymatic step for their synthesis were identified. One of the enzymes involved in this step is the dPreQ_1_ methytransferase, now named DpdM, and found in Cellulophaga phage phiSM. We showed that the Vibrio phage VH7D DpdM homolog methylated dPreQ_1_ into mdPreQ_1_ *in vivo* (Table 2). Based on the analysis of the protein structures, we propose that DpdM methylates preQ_1_ already inserted in DNA using two metal groups (Figure 3 and S5). We also identified a potential preQ_1_ or dPreQ_1_ formyltransferase, DpdN, leading to fdPreQ_1_, in the genome of Flavobacterium phage vB_FspM_immuto_2-6A. Finally, we identified a candidate protein, Cellulophaga phage phiST DpdL, that most certainly promotes alkaline decarboxylation of CDG to lead to dDG in phage genomes. Unfortunately, we were unable to demonstrate its activity.

The presence of dCDG in Sulfolobus virus SVST-2 is puzzling because this virus only encodes discernible DpdA and ArcS homologs. Furthermore, its host does not modify its tRNA with 7-deazaguanines. The proteins encoded in the vicinity of *dpdA* and *arcS* were not found in any other phage or virus encoding a 7-deazaguanine modification pathway. We do not know what the source of 7-deazaguanine for this virus is, nor how it ends up on dCDG. Indeed, we expected dG^+^ instead because Vibrio phage nt-1 ArcS produces dG^+^ (Table 3). Hence, we renamed this enzyme ArcS2 to differentiate from its tRNA-acting homolog (*21*).

Thus far, the DNA transglycosylases that insert preQ_0_ into DNA have been classified into three subfamilies: bDpdA (*4*), DpdA1, and DpdA2 (*7*). We identified two other subfamilies in viruses, DpdA3 and DpdA4. All viruses encoding 7-deazaguanine synthesis genes now encode a member a DpdA subgroup. Thus, we propose that all 7-deazaguanine DNA modifications reported to date are post-replication modifications. Interestingly, the efficiency of insertion by DpdA members varied between subgroups. bDpdA appears to have a low insertion rate, less than 0.1 % of the Gs (*4*), similar ot what was observed for DpdA2 (*7*) (Table 1). However, modification levels vary from 0.1 to 30 % of the Gs for DpdA1 (*7*). The genome of the only DpdA4 encoding virus tested (Halovirus HVTV-1) was modified at 30 % (*7*). DpdA3 is the most efficient, completely modifying the genomes of Campylobacter phage CP220 (*6*), Cellulophaga phage phiST, and Flavobacterium phage vB_FspM_immuto_2-6A (Table 1). Our attempts to test members of the DpdA3 and DpdA4 families in our *E. coli* model were unsuccessful.

In this study, we showed that the substrate specificity of some DpdA has shifted toward other 7-deazaguanines. For example, the change in substrate specificity between dPreQ_0_ and dPreQ_1_ seems to have occurred several times in evolution, as phages acquired both unimodular and bimodular QueF (Figure 3), and almost all DpdA sub-families have a member that may insert preQ_1_ into DNA (Data S1). We predict that dPreQ_0_ was the first 7-deazaguanine DNA modification, as it is the modification that requires the fewest enzymes. Interestingly, Vibrio phage VH7D DpdA2 inserted various 7-deazaguanine derivatives in its DNA with different efficiencies (Table 2). Vibrio phage nt-1 DpdA2 did not insert preQ_1_ in DNA in our assay but could insert preQ_0_ and possibly G^+^ (Table 3). It was previously reported that this phage harbored three 7-deazaguanine DNA modifications, at various levels (*7*). DpdA2 family exhibits promiscuity for substrate specificity.

We previously showed that 7-deazaguanine protect DNA from restriction enzymes at various levels depending on the modification (*7*). Phages have likely evolved different DNA modification strategies, including the addition of deazapurines in their genome, to counteract nucleic acid-based defence systems (*11–16*). It is also tempting to speculate that bacteria likely have evolved anti-phage systems targeting 7-deazaguanine. Consequently, in this “arms race” with their hosts, phages may have been driven to diversify their 7-deazaguanine into various derivatives in their genome, thereby explaining the presence of various deazapurine in viral DNA.

## SUPPLEMENTARY DATA

Supplementary Materials:

- Supplementary Text
- Supplementary Material and Methods
- Supplementary Alignments S1 to S5
- Supplementary Figure S1 to S14
- Supplementary Schemes S1 and S2
- Supplementary Tables S1 and S2
- Legends of Supplementary Data

Supplementary Data S1 to S18

Supplementary Data are available at NAR online.

## ACKNOWLEDGEMENT FUNDING

The project had partial support from the Human Frontier Science Program (grant number HFSP-RGP0024/2018) to VdCL and SM and from National Institutes of Health (GM70641 to VdCL and ES031576 to PCD) and from the National Research Foundation of Singapore under the Singapore-MIT Alliance for Research and Technology Antimicrobial Resistance IRG. SM holds the Canada Research Chair in Bacteriophages. Funding for open access charge: National Institutes of Health (GM70641).

## DATA AVAILIBILITY

Protein model predicted by AlphaFold2 were deposited in ModelArchive:

- *Cellulophaga* phage phiSM DpdM DOI: (not active) 10.5452/ma-tqgw7
- Vibrio phage VH7D DpdM DOI: (not active) 10.5452/ma-1l7yh
- *Cellulophaga* phage phiST DpdL DOI: (not active) 10.5452/ma-bxwuk
- *Flavobacterium* phage vB_FspM_immuto_2-6A DpdN DOI: (not active) 10.5452/ma-t6vzw

## REFERENCES

1. M. Mačková, S. Boháčová, P. Perlíková, L. Poštováslavětínská, M. Hocek, Polymerase Synthesis and Restriction Enzyme Cleavage of DNA Containing 7-Substituted 7-Deazaguanine Nucleobases. ChemBioChem. 16, 2225–2236 (2015).

2. P. Ménová, D. Dziuba, P. Güixens-Gallardo, P. Jurkiewicz, M. Hof, M. Hocek, Fluorescence Quenching in Oligonucleotides Containing 7-Substituted 7-Deazaguanine Bases Prepared by the Nicking Enzyme Amplification Reaction. Bioconjug. Chem. 26, 150129153326001 (2015).

3. G. Hutinet, M. A. Swarjo, V. de Crécy-Lagard, Deazaguanine derivatives, examples of crosstalk between RNA and DNA modification pathways. RNA Biol. 14, 1175–1184 (2017).

4. J. J. Thiaville, S. M. Kellner, Y. Yuan, G. Hutinet, P. C. Thiaville, W. Jumpathong, S. Mohapatra, C. Brochier-Armanet, A. V. Letarov, R. Hillebrand, C. K. Malik, C. J. Rizzo, P. C. Dedon, V. de Crécy-Lagard, Novel genomic island modifies DNA with 7-deazaguanine derivatives. Proc. Natl. Acad. Sci. U. S. A. 113, E1452–9 (2016).

5. Y. Yuan, G. Hutinet, J. G. Valera, J. Hu, R. Hillebrand, A. Gustafson, D. Iwata-Reuyl, P. C. Dedon, V. de Crécy-Lagard, Identification of the minimal bacterial 2′-deoxy-7-amido-7-deazaguanine synthesis machinery. Mol. Microbiol. 110, 469–483 (2018).

6. C. S. Crippen, Y.-J. Lee, G. Hutinet, A. Shajahan, J. C. Sacher, P. Azadi, V. de Crécy-Lagard, P. R. Weigele, C. M. Szymanski, Deoxyinosine and 7-Deaza-2-Deoxyguanosine as Carriers of Genetic Information in the DNA of Campylobacter Viruses. J. Virol. 93, 1–14 (2019).

7. G. Hutinet, W. Kot, L. Cui, R. Hillebrand, S. Balamkundu, S. Gnanakalai, R. Neelakandan, A. B. Carstens, C. F. Lui, D. Tremblay, D. Jacobs-sera, M. Sassanfar, Y. Lee, P. Weigele, S. Moineau, G. F. Hatfull, P. C. Dedon, L. H. Hansen, V. de Crécy-Lagard, 7-Deazaguanine modifications protect phage DNA from host restriction systems. Nat. Commun. 10, 1–12 (2019).

8. P. Weigele, E. A. Raleigh, Biosynthesis and Function of Modified Bases in Bacteria and Their Viruses. Chem. Rev. 116, 12655–12687 (2016).

9. G. Hutinet, Y. Lee, Hypermodified DNA in Viruses of E. coli and Salmonella. EcoSal Plus. 9 (2021), doi:https://doi.org/10.1128/ecosalplus.ESP-0028-2019.

10. Y. Lee, N. Dai, I. M. Stephanie, C. Guan, M. J. Parker, M. E. Fraser, S. E. Walsh, J. Sridar, A. Mulholland, K. Nayak, Z. Sun, Y. Lin, D. G. Comb, K. Marks, R. Gonzalez, D. P. Dowling, V. Bandarian, L. Saleh, I. R. Corr, P. R. Weigele, NAR Breakthrough Article Pathways of thymidine hypermodification ^ Jr, 1–17 (2021).

11. Y. Wang, H. Fan, Y. Tong, Unveil the Secret of the Bacteria and Phage Arms Race. Int. J. Mol. Sci. 24, 1–24 (2023).

12. L. Gao, H. Altae-Tran, F. Böhning, K. S. Makarova, M. Segel, J. L. Schmid-Burgk, J. Koob, Y. I. Wolf, E. V Koonin, F. Zhang, Diverse enzymatic activities mediate antiviral immunity in prokaryotes. Science. 369, 1077–1084 (2020).

13. A. B. Isaev, O. S. Musharova, K. V. Severinov, Microbial Arsenal of Antiviral Defenses – Part I. Biochem. 86, 319–337 (2021).

14. A. B. Isaev, O. S. Musharova, K. V. Severinov, Microbial Arsenal of Antiviral Defenses. Part II. Biochem. 86, 449–470 (2021).

15. S. Doron, S. Melamed, G. Ofir, A. Leavitt, A. Lopatina, M. Keren, G. Amitai, R. Sorek, Systematic discovery of antiphage defense systems in the microbial pangenome. Science (80-.). 359, 0–12 (2018).

16. A. Millman, S. Melamed, A. Leavitt, S. Doron, A. Bernheim, J. Hör, J. Garb, N. Bechon, A. Brandis, A. Lopatina, G. Ofir, D. Hochhauser, A. Stokar-Avihail, N. Tal, S. Sharir, M. Voichek, Z. Erez, J. L. M. Ferrer, D. Dar, A. Kacen, G. Amitai, R. Sorek, An expanded arsenal of immune systems that protect bacteria from phages. Cell Host Microbe. 30, 1556–1569.e5 (2022).

17. W. Kot, N. S. Olsen, T. K. Nielsen, G. Hutinet, V. De Crécy-Lagard, L. Cui, P. C. Dedon, A. B. Carstens, S. Moineau, M. A. Swairjo, L. H. Hansen, Detection of preQ0 deazaguanine modifications in bacteriophage CAjan DNA using Nanopore sequencing reveals same hypermodification at two distinct DNA motifs. Nucleic Acids Res. 48, 10383–10396 (2020).

18. K. Moeller, G. S. Nguyen, F. Hollmann, U. Hanefeld, Expression and characterization of the nitrile reductase queF from E. coli. Enzyme Microb. Technol. 52, 129–133 (2013).

19. S. G. Van Lanen, J. S. Reader, M. A. Swairjo, V. de Crécy-Lagard, B. Lee, D. Iwata-Reuyl, From cyclohydrolase to oxidoreductase: Discovery of nitrile reductase activity in a common fold. Proc. Natl. Acad. Sci. U. S. A. 102, 4264–4269 (2005).

20. B. Stengl, K. Reuter, G. Klebe, Mechanism and substrate specificity of tRNA-guanine transglycosylases (TGTs): tRNA-modifying enzymes from the three different kingdoms of life share a common catalytic mechanism. ChemBioChem. 6, 1926–1939 (2005).

21. G. Phillips, V. M. Chikwana, A. Maxwell, B. El-Yacoubi, M. A. Swairjo, D. Iwata-Reuyl, V. De Crécy-Lagard, Discovery and characterization of an amidinotransferase involved in the modification of archaeal tRNA. J. Biol. Chem. 285, 12706–12713 (2010).

22. G. Phillips, M. a. Swairjo, K. W. Gaston, M. Bailly, P. a. Limbach, D. Iwata-Reuyl, V. De Crécy-Lagard, Diversity of archaeosine synthesis in crenarchaeota. ACS Chem. Biol. 7, 300–305 (2012).

23. A. Bon Ramos, L. Bao, B. Turner, V. de Crécy-Lagard, D. Iwata-Reuyl, QueF-Like, a Non-Homologous Archaeosine Synthase from the Crenarchaeota. Biomolecules. 7, 1–14 (2017).

24. L. Zimmermann, A. Stephens, S. Z. Nam, D. Rau, J. Kübler, M. Lozajic, F. Gabler, J. Söding, A. N. Lupas, V. Alva, A Completely Reimplemented MPI Bioinformatics Toolkit with a New HHpred Server at its Core. J. Mol. Biol. 430, 2237–2243 (2018).

25. A. Hildebrand, M. Remmert, A. Biegert, J. Söding, Fast and accurate automatic structure prediction with HHpred. Proteins Struct. Funct. Bioinforma. 77, 128–132 (2009).

26. J. Mistry, S. Chuguransky, L. Williams, M. Qureshi, G. A. Salazar, E. L. L. Sonnhammer, S. C. E. Tosatto, L. Paladin, S. Raj, L. J. Richardson, R. D. Finn, A. Bateman, Pfam: The protein families database in 2021. Nucleic Acids Res. 49 (2021), doi:10.1093/nar/gkaa913.

27. S. F. Altschul, T. L. Madden, A. A. Schäffer, J. Zhang, Z. Zhang, W. Miller, D. J. Lipman, Gapped BLAST and PSI-BLAST: A new generation of protein database search programs. Nucleic Acids Res. 25, 3389–3402 (1997).

28. C. Camacho, G. Coulouris, V. Avagyan, N. Ma, J. Papadopoulos, K. Bealer, T. L. Madden, BLAST+: Architecture and applications. BMC Bioinformatics. 10, 1–9 (2009).

29. K. Katoh, J. Rozewicki, K. D. Yamada, MAFFT online service: Multiple sequence alignment, interactive sequence choice and visualization. Brief. Bioinform. 20, 1160–1166 (2018).

30. M. Matsui, W. Iwasaki, Graph Splitting: A Graph-Based Approach for Superfamily-Scale Phylogenetic Tree Reconstruction. Syst. Biol. 69, 265–279 (2020).

31. J. Jumper, R. Evans, A. Pritzel, T. Green, M. Figurnov, O. Ronneberger, K. Tunyasuvunakool, R. Bates, A. Žídek, A. Potapenko, A. Bridgland, C. Meyer, S. A. A. Kohl, A. J. Ballard, A. Cowie, B. Romera-Paredes, S. Nikolov, R. Jain, J. Adler, T. Back, S. Petersen, D. Reiman, E. Clancy, M. Zielinski, M. Steinegger, M. Pacholska, T. Berghammer, S. Bodenstein, D. Silver, O. Vinyals, A. W. Senior, K. Kavukcuoglu, P. Kohli, D. Hassabis, Highly accurate protein structure prediction with AlphaFold. Nature. 596, 583–589 (2021).

32. E. F. Pettersen, T. D. Goddard, C. C. Huang, E. C. Meng, G. S. Couch, T. I. Croll, J. H. Morris, T. E. Ferrin, C. E. Thomas Ferrin, UCSF ChimeraX: Structure visualization for researchers, educators, and developers. Protein Sci. 30, 70–82 (2021).

33. H. M. Berman, J. Westbrook, Z. Feng, G. Gilliland, T. N. Bhat, H. Weissig, I. N. Shindyalov, P. E. Bourne, The Protein Data Bank. Nucleic Acids Res. 28, 235–242 (2000).

34. K. Holmfeldt, N. Solonenko, M. Shah, K. Corrier, L. Riemann, N. C. VerBerkmoes, M. B. Sullivan, Twelve previously unknown phage genera are ubiquitous in global oceans. Proc. Natl. Acad. Sci. U. S. A. 110, 12798–12803 (2013).

35. B. W. K. Lee, S. G. Van Lanen, D. Iwata-Reuyl, Mechanistic studies of Bacillus subtilis QueF, the nitrile oxidoreductase involved in queuosine biosynthesis. Biochemistry. 46, 12844–12854 (2007).

36. X. Mei, J. Alvarez, A. Bon Ramos, U. Samanta, D. Iwata-Reuyl, M. A. Swairjo, Crystal Structure of the Archaeosine Synthase QueF-Like– Insights into Amidino Transfer and tRNA Recognition by the Tunnel Fold. Proteins. 165, 255–269 (2016).

37. T.-Q. Nguyen, Y. Nicolet, Structure and Catalytic Mechanism of Radical SAM Methylases. Life. 12, 1732 (2022).

38. B. Rihtman, S. Bowman-Grahl, A. Millard, R. M. Corrigan, M. R. J. Clokie, D. J. Scanlan, Cyanophage MazG is a pyrophosphohydrolase but unable to hydrolyse magic spot nucleotides. Environ. Microbiol. Rep. 11, 448–455 (2019).

39. N. Colloc’h, A. Poupon, J. P. Mornon, Sequence and structural features of the T-fold, an original tunnelling building unit. Proteins Struct. Funct. Genet. 39, 142–154 (2000).

40. G. Phillips, L. L. Grochowski, S. Bonnett, H. Xu, M. Bailly, C. Blaby-Haas, B. El Yacoubi, D. Iwata-Reuyl, R. H. White, V. De Crécy-Lagard, Functional promiscuity of the COG0720 family. ACS Chem. Biol. 7, 197–209 (2012).

41. Z. D. Miles, S. A. Roberts, R. M. McCarty, V. Bandarian, Biochemical and structural studies of 6-carboxy-5,6,7,8-tetrahydropterin synthase reveal the molecular basis of catalytic promiscuity within the tunnel-fold superfamily. J. Biol. Chem. 289, 23641–23652 (2014).

42. Z. Zhang, T. T. Caradoc-Davies, J. M. Dickson, E. N. Baker, C. J. Squire, Structures of Glycinamide Ribonucleotide Transformylase (PurN) from Mycobacterium tuberculosis Reveal a Novel Dimer with Relevance to Drug Discovery. J. Mol. Biol. 389, 722–733 (2009).

