## Supplementary Material for "Four additional natural 7-deazaguanine derivatives in phages and how to make them"

**sSupplemental material for**

### Contents

Supplementary Text ... 3
DpdL proteins are likely a homohexamer that bind CDG ... 3
DpdN uses methyltetrahydrofolate as formyl donor ... 3
Supplemental Materials and Methods ... 4
Cloning of phage genes ... 4
QueF mutant ... 4
Phage DNA purification ... 4
Deazaguanine synthesis ... 5
Northern blot with APB-acrylamide gel ... 9
Structure flexibility prediction ... 10
Supplementary Alignments ... 11
Alignment S1 : QueF ... 11
Alignment S2: DpdM ... 14
Alignment S3: DpdL ... 15
Alignment S4: DpdN ... 20
Alignment S5: ArcS ... 24
Supplementary References ... 26
Supplementary Figures ... 27
Fig. S1. LC-MS analysis of digested Cellulophaga phage phiSM DNA sample and methyl-dPreQ1 (mdPreQ1) standard ... 28
Fig. S2. LC-MS nucleoside analysis of genomic DNA isolated from *Cellulophaga* phages phi47:1 and phi38:2. ... 29
Fig. S3. Phage QueF are NADPH-dependent 7-cyano-7-deazaguanine reductase. ... 30
Fig. S4. LC-MS analysis of digested *Vibrio* phage phi-Grn1 and phi-ST2 DNA samples and methyl-dPreQ1 standard ... 31
Fig. S5. DpdM structure prediction ... 32
Fig. S6. LC-MS analysis of digested *Cellulophaga* phage phiST DNA sample and methyl-2'-deoxy-7-deazagunine (dDG) standard ... 33
Fig. S7. LC-MS nucleoside analysis of genomic DNA isolated from *Cellulophaga* phages phi19:2 and phi13:1. ... 34
Fig. S8. DpdL structure prediction ... 35
Fig. S9. 2D NMR spectra of formyl-dPreQ1 (fdPreQ1) standard ... 36

#### Supplementary Text

##### DpdL proteins are likely a homohexamer that bind CDG

Cellulophaga phage phiST DpdL, like other T-Fold enzymes such as QueD (Data S11), was best modeled as a homohexamer (Figure S8A) composed of two rings of trimers (Figure S8B and C). The protein's core ring structure aligned well with the core structure of *E. coli* QueD (Figure S8D and E). The three conserved histidine residues known to bind zinc in QueD are also conserved and found in the same location in the DpdL protein (Figure S8F), implying that DpdL binds a metal, most likely zinc. The predicted pocket contains almost all the conserved residues in the alignment. As observed for other T-fold enzymes (ref), the pocket is at the interface of two subunits (Figure S8G), one of which participates with the three conserved histidines H17, H37, and H39, as well as other residues (G15, L33, F41, S83, E85 and E104), and the other with R54 and E57. R38 from a third subunit closes the pocket, an amino acid that is replaced by a phenylalanine in the DpdL encoded in *Pseudomonas* phage vB_PaeM_PA5oct and *Aeromonas* phage Ahp1_CNU-2021 (see alignment above). Interestingly, the three molecules, CPH4 (Figure S8H), CDG (Figure S8I), and DG (Figure S8J) can all fit in this pocket. However, only a base can fit, ruling out the possibility that the enzyme's substrate is a 7-deazaguanine inserted in DNA. A prediction of the flexibility of the protein (Figure S8K) revealed that the residue R54 is most likely involved in the pocket's opened/closed configuration (Figure S8L).

##### DpdN uses methyltetrahydrofolate as formyl donor

Flavobacterium phage vB_FspM_immuto_2-6A DpdN is predicted to fold into an L-shape with a mildly positively charged cross-section where DNA could bind (Figure S12A and B). The conserved residues are mostly found inside of the L (Figure S12C and D). The conserved residues specific to the four proteins predicted to be true DpdN (in green in Figure S12, corresponding to) do not cluster in a specific region, but instead split in two groups: one around the positively charged groove and one across what could be a binding pocket. Indeed, when overlayed with *Mycobacterium tuberculosis* PurN (FigureS12E and F), the formylase's general fold matched, as predicted. The formyl donor of PurN, 5-methyl-5,6,7,8-tetrahydrofolate, does fit in the pocket described above and is thus almost certainly also the formyl donor for DpdN.

#### Supplemental Materials and Methods

##### Cloning of phage genes

Phage genes were synthesized by TWIST bioscience (https://www.twistbioscience.com, South San Francisco, CA) and introduced into the pTwist Kan High Copy vector. These genes were coding for: Cellulophaga phage phiSM QueF (YP_007675731), DpdM (YP_007675730), and DpdA1 (YP_007675729), Vibrio phage VH7D QueF (YP_009006087), DpdM (YP_009006155), and DpdA2 (YP_009006086), Acidovorax phage ACP17 DpdL (YP_009609725), an dDpdA2 (YP_009609723), and an uncultured phage DpdL (AGH13910), and DpdA1 (AGH13911). A KpnI site and a strong RBS sequence (5'-AGGAATTCACC-3') were added at the 5' end of the genes and a SalI site at the 3' end. Each gene was then cloned in between the KpnI and SalI sites of both pBAD24 and pBAD33 *E. coli* vector. Halovirus HVTV-1 QueF (YP_007378987) encoding gene was amplified by PCR using primers GO442 (5'-GCACGCCATGGGAGACAACGAAGTCAAC-3') and GO443 (5'-GCACGCCTGCAGGTCACTGGATACCTCCGACC-3') and then cloned in between the NcoI and SbfI sites of pBAD24 vector. Vibrio phage nt-1 DpdA2 (YP_008125322) and ArcS (YP_008125320) encoding genes were amplified by PCR using primers GO488 (5'-ATGATGGTACCAGGAGGAATTCACCTTGATTATTGAATACGTAATGTCAGCG-3') and GO489 (5'-ATGATGCATGCTTATGATTTAAAATCTGGGTATTCAGCAG-3'), as well as GO490 (5'-ATGATGGTACCAGGAGGAATTCACCATGAATAGCGTACAAGATAAATCAGTTC-3') and GO491 (5'-ATGATGCATGCCTAAAAGAAGTTAGCTAAGCTATTTTCTTTC-3'), respectively, then cloned in between the KpnI and SphI sites of pBAD24 and pBAD33 vectors. Each plasmid is listed in Table S2. Plasmids were transformed in appropriate *E. coli* strains for testing, as described in the text and referenced in Table S1.

##### QueF mutant

The deletion of queF was obtained from the keio collection (1) and transduced using P1vir into MG1655 (2).

##### Phage DNA purification

*All Cellulophaga phages and Flavobacterium phage vB_FspM_immuto_2-6A*

All Cellulophaga phages and Flavobacterium phage vB_FspM_immuto_2-6A were replicated using the top-agar plating technique as described previously (3) Briefly, viral dilutions and overnight bacterial cultures were mixed with top agar (450 mM NaCl, 50 mM $MgSO_4*7H_2O$, 50 mM Trizma base, pH 8, 0.5 % low melting point agarose) and poured out onto Zobell-agar plates (1 g yeast extract, 5 g Bacto peptone, and 15 g Bacto Agar in 800 mL of Baltic Sea water and 200 mL of Milli-Q water). To create high-titer phage-stocks, 5 mL marine sodium magnesium buffer (MSM: 450 mM NaCl, 50 mM $MgSO_4*7H_2O$, 50 mM Trizma base, pH 8) was added to fully lysed plates. The top-agar layer was shredded with an inoculation loop and incubated on a shaker (50 rpm for > 30 min). The phage mix was collected into a falcon tube, centrifuged at 2860 x g for 15 min, and filtered through a 0.2 μm syringe filter. Insufficiently concentrated phage lysates were collected as above from 20 plates and concentrated with polyethylene glycol (PEG). Sodium chloride was added to the resulting lysate at 1 M, and PEG8000 was added at 10 % w/v. The resulting mixture was incubated at 4 °C for 1 hour - overnight, then centrifuged for 10 min at 10,000 g to collect the pellet. The pellet was resuspended in 1 mL MSM. One ml of each phage stock was used for DNA extraction using the Wizard PCR DNA Purification kit (Promega, Madison, WI, USA) as previously described (4, 5). Briefly, the phages were mixed with 1 mL wizard resin and pushed through a mini column using a sterile syringe. The mini column was washed twice with 1 ml 80% isopropanol, followed by centrifugation at 10,000 x g for 2 min to remove the isopropanol, and the DNA was eluted by adding 100 μL 80°C 10 mM Tris (pH 8) and centrifuging at 10,000 x g for 30 s.

*Sulfolobus virus STSV-2*

Two liters of STSV2-infected culture of *Sulfolobus tengchongensis* HB52 (6) were grown until late exponential phase in TYS medium (Sulfolobus medium supplemented with 0.2 % tryptone, 0.1 % yeast and 0.2 % sucrose (7) at 78 °C under agitation. Virions were separated from cell debris by centrifugation at 11 000 x g for 10 min. Polyethylene glycol and NaCl were added to a final concentration of 10 % and 1 M, respectively, the virus preparation was incubated overnight at 4°C and pelleted through centrifugation at 11 000 x g for 10 min. Virus was then recovered by resuspending the pellet in 10 mM Tris-acetate pH 6 and the remaining cell debris was removed through centrifugation at 4,085 x g for 5 min. The resulting virus preparation was treated with benzonase (final concentration 1.5 mM) at 37 °C for 1 h to remove extracellular nucleic acids. Finally, STSV2 DNA was extracted using 25:24:1 phenol:chloroform:isoamyl alcohol pH 8, followed by isopropanol precipitation and resuspended in TE buffer (10 mM Tris-HCl pH 8, 1 mM EDTA).

*Vibrio phages*

The bacterial strain *Vibrio alginolyticus* VI was grown in LB at 25°C with agitation (8). For phages Grn1 and St2 amplification, $MgSO_4$ and $CaCl_2$ were added to LB at a final concentration of 1 mM each. DNA was extracted from phage lysates using the phenol-chloroform method previously described (9).

**Deazaguanine synthesis**

All reactions were carried out in oven-dried glassware under a positive pressure of argon or nitrogen, unless otherwise mentioned, with magnetic stirring. Air-sensitive reagents and solutions were transferred via syringe or cannula and were introduced to the apparatus via rubber septa. All reagents, starting materials, and solvents were obtained from commercial suppliers and used as such without further purification. Reactions were monitored by thin-layer chromatography (TLC) with 0.25 mm precoated silica gel plates (60 F254). Visualization was accomplished either with UV light or by immersion in an ethanolic solution of phosphomolybdic acid (PMA), para-anisaldehyde, 2,4-DNP, $KMnO_4$, Ninhydrin solution, or iodine adsorbed on silica gel, followed by heating with a heat gun for ~15 s. Column chromatography was performed on silica gel (230−400 mesh size). Deuterated solvents for NMR spectroscopic analyses were used as received. All $^{1}$H NMR spectra were obtained using a 400 MHz spectrometer. Coupling constants were measured in Hertz. All chemical shifts were quoted in parts per million, relative to TMS, using the residual solvent peak as a reference standard. The following abbreviations were used to explain the multiplicities: s = singlet, d = doublet, t = triplet, q = quartet, m = multiplet, b = broad. Chemical nomenclature was generated using Chem Bio Draw Ultra 13.0.

***Synthesis of Methyl-dPreQ$_1$***

The synthesis (Scheme S1) was commenced from commercially available Hoffer's chloro sugar (**1**) and 7-deazapurine derivative (**2**), glycosylation of **2** using KOH afforded compound **3**, the exocyclic $NH_2$ was protected using isobutyryl chloride in pyridine to afford compound **4**, which on treatment with N-Iodosuccinimide in DMF at 80 ºC afforded iodo derivative **5**, which on treatment with pyrine-2-carboldoxime and tetramethylguanidine afforded compound **6**. The toluly protection in **6** was removed using NaOMe to get compound **7**. The iodo in compound **7** was converted to aldehyde **8** using carbon monoxide (CO), $Pd_2(dba)_3$ and $Bu_3SnH$ in DMF. The aldehyde group in compound **8** converted to aminomethyl using reductive amination with methylamine to afford compound **9**, during which some percentage of isobutyryl protecting group was deprotected, aq. $NH_3$ was added and heated at 50 ºC to completely remove to afford Methyl-dPreQ$_1$. Experimental procedures for each step as follow:

*(2R,3S,5R)-5-(2-Amino-4-chloro-7H-pyrrolo[2,3-d]pyrimidin-7-yl)-2-(((4-methylbenzoyl)oxy)methyl)tetrahydrofuran-3-yl 4-methylbenzoate* ***(3)***

KOH (3.5 g, 62.5 mmol) was added portion wise at room temperature to a suspension of **2** (3.0 g, 17.8 mmol) and stirred at RT for 10 min, Hoffer's chloro sugar (9.0 g, 23.14 mmol) was added portion wise and stirred at RT for 30 min. Reaction mixture was filtered and the filtrate was evaporated under reduced pressure and the crude was triturated with diethyl ether -Hexanes to get compound **3** (7.2 g, 78%) as pale yellow solid. $^{1}$H NMR (DMSO-D$_6$, 400 MHz) δ 7.87-7.95 (m, 4H), 7.31-7.39 (m, 5H), 6.80 (s, 2H), 6.55 (dd, *J* = 5.7, 8.7 Hz, 1H), 6.38 (d, *J* = 3.9 Hz, 1H), 5.67 (d, *J* = 6.0 Hz, 1H), 4.47-4.63 (m, 3H), 2.94-3.04 (m, 1H), 2.61-2.68 (m, 1H), 2.37, 2.40 (2s, 6H); LCMS (ESI): *m/z* calculated for $C_{27}H_{26}ClN_4O_5$ $[M + H]^+$ 521.16 found 521.18.

*(2R,3S,5R)-5-(4-Chloro-2-isobutyramido-7H-pyrrolo[2,3-d]pyrimidin-7-yl)-2-(((4-methylbenzoyl)oxy)methyl)tetrahydrofuran-3-yl 4-methylbenzoate* ***(4)***

Isobutyryl chloride (1.32 mL, 13.1 mmol) was added dropwise to a solution of **3** in anhydrous pyridine (40 mL) at 0 °C, and RM was allowed to stir at RT for 1h, RM was quenched with MeOH (2.0 mL), evaporated, the residue was dissolved in EtOAc (100 mL), washed with 1 N HCl (50 mL), water (50 mL) dried over anhydrous $Na_2SO_4$, evaporated under reduced pressure. The crude was triturated with diethyl ether to get compound **4** (6.1 g, 86%) as off-white solid. $^{1}$H NMR ($CDCl_3$) δ 8.04 (s, 1H), 7.98 (dt, *J* = 8.3, 1.7 Hz, 2H), 7.90 (dt, *J* = 8.3, 1.7 Hz, 2H), 7.28 (d, *J* = 8.7 Hz, 2H), 7.27 (d, *J* = 3.7 Hz, 1H), 7.22 (d, *J* = 8.0 Hz, 2H), 6.69 (dd, *J* = 8.1, 6.0 Hz, 1H), 6.51 (d, *J* = 3.8 Hz, 1H), 5.78 (dt, *J* = 6.3, 2.3 Hz, 1H), 4.76 (dd, *J* = 11.5, 3.8 Hz, 1H), 4.63 (dd, *J* = 11.5, 4.4 Hz, 1H), 4.59 (ddd, *J* = 4.2, 4.1, 2.7 Hz, 1H), 3.06– 2.92 (m, 1H) 2.97 (ddd, *J* = 14.3, 8.1,

6.6Hz, 1H), 2.76 (ddd, $J$ = 14.3, 6.0, 2.4 Hz, 1H), 2.44 (s, 3H), 2.41 (s, 3H), 1.28 (d, $J$ = 6.9 Hz, 6H); LCMS (ESI): $m/z$ calculated for $C_{31}H_{32}ClN_4O_6$ $[M + H]^+$ 591.20 found 591.23.

*(2R,3S,5R)-5-(4-Chloro-5-iodo-2-isobutyramido-7H-pyrrolo[2,3-d]pyrimidin-7-yl)-2-(((4-methylbenzoyl)oxy)methyl)tetrahydrofuran-3-yl 4-methylbenzoate **(5)***

N-Iodosuccinimide (15.9 g, 71.06 mmol) was added to a solution of **4** (6.0 g, 10.15 mmol) in anhydrous DMF (300 mL) and stirred at 80 °C for 8h, RM was quenched with Sat. $NaHCO_3$ solution (150 mL), extracted with EtOAc (3 x 150 mL), the combined organic layer was washed with Sat. $Na_2S_2O_3$ solution (100 mL), followed by water (100 mL) and brine (100 mL). The crude was triturated with diethyl ether to afford compound **5** (5.9 g, 82%) as pale-yellow solid. $^1$H NMR ($CDCl_3$) δ 8.14 (s, 1H), 7.96 (dt, $J$ = 8.2, 1.5 Hz, 2H), 7.90 (dt, $J$ = 8.2, 1.5 Hz, 2H), 7.41 (s, 1H), 7.27 (d, $J$ = 7.9 Hz, 2H), 7.24 (d, $J$ = 7.9 Hz, 2H), 6.67 (dd, $J$ = 7.8, 6.2 Hz, 1H), 5.76 (dt, $J$ = 6.3, 2.5 Hz, 1H), 4.77 (dd, $J$ = 12.0, 3.9 Hz, 1H), 4.65 (dd, $J$ = 12.0, 3.9 Hz, 1H), 4.59 (dd, $J$ = 6.5, 3.8 Hz, 1H), 2.98 (m, 1H), 2.88 (ddd, $J$ = 14.4, 7.7, 6.6 Hz, 1H), 2.77 (ddd, $J$ = 14.3, 6.1, 2.6 Hz, 1H), 2.43 (s, 3H), 2.41 (s, 3H), 1.28 (d, $J$ = 6.9 Hz, 6H); LCMS (ESI): $m/z$ calculated for $C_{31}H_{31}ClIN_4O_6$ $[M + H]^+$ 717.10 found 717.18.

*(2R,3S,5R)-5-(5-Iodo-2-isobutyramido-4-oxo-3,4-dihydro-7H-pyrrolo[2,3-d]pyrimidin-7-yl)-2-(((4-methylbenzoyl)oxy)methyl)tetrahydrofuran-3-yl 4-methylbenzoate **(6)***

Tetremethylguanidine (1.1 mL, 8.37 mmol) was added to a solution of compound **5** (3.0 g, 4.19 mmol) and pyridine 2-Pyridinecarbaldehyde oxime (1.03 g, 8.37 mmol) in anhydrous DMF (50 mL), anhydrous 1,4-dioxane (50 mL) at RT and allowed to stir for 24h, RM was diluted with EtOAc (200 mL) and washed with Sat. $NaHCO_3$ (100 mL), 1N HCl (100 mL) followed by $H_2O$ (100 mL) combined organic layers dried over dried over anhydrous $Na_2SO_4$, evaporated under reduced pressure. Crude was purified by silica gel column chromatography using (0-50% EtOAc in hexanes to afford compound **6** (2.20 g, 68 %) as off-white solid. $^1$H NMR ($CDCl_3$, 400 MHz) δ 11.68 (s, 1H), 8.79 (s, 1H), 7.94 (d, $J$ = 8.20 Hz, 2H), 7.89 (d, $J$ = 8.20 Hz, 2H), 7.26 (m, 4H), 6.94 (s, 1H), 6.27 (t, $J$ = 6.94 Hz, 1H), 5.81 (m, 1H), 5.01 (q, $J$ = 6.66 Hz, 1H), 4.61 (m, 2H), 3.01 (m, 1H), 2.71 (m, 1H), 2.59 (m, 1H), 2.45 (s, 3H), 2.43 (s, 3H), 1.31 (d, $J$ = 3.84 Hz, 3H), 1.30 (d, $J$ = 3.84 Hz, 3H); LCMS (ESI): $m/z$ calculated for $C_{31}H_{31}IN_4O_7$ $[M + H]^+$ 698.12 found 698.16.

*N-(7-((2R,4S,5R)-4-Hydroxy-5-(hydroxymethyl)tetrahydrofuran-2-yl)-5-iodo-4-oxo-4,7-dihydro-3H-pyrrolo[2,3-d]pyrimidin-2-yl)isobutyramide **(7)***

1 M NaOMe (5.73 mL, 5.73 mmol) was added dropwise to a solution of compound **6** (2.00 g, 1.86 mmol) in THF (20 mL) and MeOH (0.5 mL) at 0 ºC and allowed to stir at RT for 2 h, RM was neutralized with AcOH (0.5 mL) and evaporated and purified by silica gel column chromatography using (0-5 % MeOH in DCM to afford compound **7** (920 mg, 69 %) as off-white solid. $^1$H NMR (DMSO-$D_6$, 400 MHz) δ 11.78 (s, 1H), 11.55 (s, 1H), 7.45 (s, 1H), 6.36 (dt, $J$ = 5.7, 8.4 Hz, 1H), 5.24 (d, $J$ = 3.0 Hz, 1H), 4.93 (t, $J$ = 5.4 Hz, 1H), 4.31 (m, 1H), 3.78 (m, 1H), 3.50 (m, 2H), 2.74 (m, 1H), 2.38 (m, 1H), 2.12 (m, 1H), 1.12 (d, $J$ = 3.84 Hz, 3H), 1.10 (d, $J$ = 3.84 Hz, 3H); LCMS (ESI): $m/z$ calculated for $C_{15}H_{19}IN_4O_5$ $[M + H]^+$ 462.04 found 462.09.

*N-(5-Formyl-7-((2R,4S,5R)-4-hydroxy-5-(hydroxymethyl)tetrahydrofuran-2-yl)-4-oxo-4,7-dihydro-3H-pyrrolo[2,3-d]pyrimidin-2-yl)isobutyramide* ***(8)***

$Pd_2(dba)_3$ (20 mg, 0.022 mmol) was added to a degassed solution of compound **7** (100 mg, 0.22 mmol), $PPh_3$ (35 mg, 0.132 mmol) in anhydrous THF. The RM was purged with carbon monoxide gas and heated to 60 ºC, $Bu_3SnH$ was added dropwise over 1h and the reaction further stirred for 18h under CO gas (balloon was used), RM was evaporated and purified by silica gel column chromatography using (0-50% EtOAc in hexanes to afford compound **8** (65 mg, 83%) as off-white solid. LCMS (ESI): *m/z* calculated for $C_{16}H_{20}N_4O_6$ $[M + H]^+$ 364.14 found 364.18.

*N-(7-((2R,4S,5R)-4-Hydroxy-5-(hydroxymethyl)tetrahydrofuran-2-yl)-5-((methylamino)methyl)-4-oxo-4,7-dihydro-3H-pyrrolo[2,3-d]pyrimidin-2-yl)isobutyramide* ***(9)***

$NaCNBH_3$ (9.0 mg, 0.15 mmol) was added to a solution of compound **8** (18 mg, 0.05mmol) and Methylamine (40% in Water, 1.0 mmol) and stirred at RT for 18h, rm was evaporated and crude carried to next step without further purification.

*2-Amino-7-((2R,4S,5R)-4-hydroxy-5-(hydroxymethyl)tetrahydrofuran-2-yl)-5-((methylamino)methyl)-3,7-dihydro-4H-pyrrolo[2,3-d]pyrimidin-4-one* ***(10)***

Aqueous $NH_3$ (2.0 mL) was added to crude compound **11** from above reaction and stirred at 50 °C for 2h, rm was evaporated and crude was purified by HPLC, fractions containing product were pooled and lyophilized to afford **Methyl-dPreQ$_1$** (5.0 mg, 32%) as off-white solid. $^1$H NMR ($CD_3OD$, 400 MHz) δ 7.16 (s, 1H), 6.40 (q, *J* = 4.66 Hz, 1H), 4.49 (t, *J* = 2.89 Hz, 1H), 4.24 (s, 2H), 3.96 (m, 1H), 3.74 (m, 2H), 2.74 (s, 3H), 2.28 (m, 1H), 2.52 (m, 1H); HRMS (ESI): *m/z* calculated for $C_{13}H_{20}N_5O_4$ $[M + H]^+$ 310.1515 found 310.1509.

***Synthesis of dPreQ$_1$ and Formyl-dPreQ$_1$***

The aldehyde **8** was synthesized according Scheme 1, which was converted to dPreQ$_1$ using reductive amination with ammonia in MeOH/$NaBH_4$, during which some percentage of isobutyryl protecting group was deprotected, aq. $NH_3$ was added and heated at 50 ºC to completely remove to afford dPreQ$_1$. Selective formylation of dPreQ1 using acetic anhydride/formic acid gave Formyl-dPreQ$_1$ (Scheme S2). The position of formyl group was unambiguously confirmed by 2D NMR analysis. Experimental procedures for each step as follow:

*2-Amino-5-(aminomethyl)-7-((2R,4S,5R)-4-hydroxy-5-(hydroxymethyl)tetrahydrofuran-2-yl)-3,7-dihydro-4H-pyrrolo[2,3-d]pyrimidin-4-one* ***(11)***

Compound **8** (30 mg, 0.083 mmol) was dissolved in 7N $NH_3$ in MeOH and stirred at RT for 4h, RM was evaporated and dissolved in MeOH cooled to 0 ºC and $NaBH_4$ was added and stirred at RT for 30 min, RM was evaporated and re-dissolved in aq.$NH_3$ and heated at 50 ºC for 2h. RM was evaporated and purified by reverse phase HPLC to afford compound **11, dPreQ$_1$** (9.0 mg,

37%) as off-white solid. HRMS (ESI): *m/z* calculated for $C_{12}H_{17}N_5O_4$ $[M + H]^+$ 295.1281 found 295.1281.

*N-((2-Amino-7-((2R,4S,5R)-4-hydroxy-5-(hydroxymethyl)tetrahydrofuran-2-yl)-4-oxo-4,7-dihydro-3H-pyrrolo[2,3-d]pyrimidin-5-yl)methyl)formamide* ***(12)***

Formic acid (0.25 mmol) was added dropwise to Acetic anhydride (0.25 mmol) at 0 ºC and this mixture was added compound **9** (9.0 mg) at 0 ºC and stirred for 30 min followed by 18h at RT. RM was diluted with MeOH (5.0 mL) evaporated and purified by reverse phase HPLC to afford compound **12, Formyl-dPreQ$_1$** (2.1 mg) as off-white solid. $^1$H NMR ($CD_3OD$, 400 MHz) δ 8.10 (s, 1H), 6.89 (s, 1H), 6.37 (dd, *J* = 6.12, 2.12 Hz, 1H), 4.49 (s, 2H), 4.47-4.45 (m, 1H), 3.94 (q, *J* = 2.76 Hz, 1H), 3.77-3.67 (m, 2H), 2.56-2.49 (m, 1H), 2.26-2.22 (m, 1H);$^{13}$C NMR ($CD_3OD$, 100 MHz) δ 161.9, 160.5, 152.7, 151.4, 115.8, 115.3, 99.2, 87.1, 83.9, 71.6, 62.3, 39.7, 33.6; HRMS (ESI): *m/z* calculated for $C_{13}H_{18}N_5O_5$ $[M + H]^+$ 324.1308 found 324.1311.

**Nucleoside analysis of Cellulophaga phage genomic DNA.**

Genomic DNA from Cellulophaga phages phi19:2, phi13:1, phi38:2, and phi47:1 were enzymatically digested to free nucleosides and the resulting mixture subjected to liquid chromatography-mass spectrometry (LC-MS) analysis. Approximately 1-2 µg phage DNA was treated with the Nucleoside Digestion Mix (New England Biolabs, cat# M0649) following the manufacturer's protocol at 37°C for >1h. The resulting mixture of nucleosides were filtered through a hydrophilic PTFE 0.2 µm centrifugal filter, and the filtrate was subjected to the reverse phase LC-MS for nucleoside separation and MS detection.

LC-MS was performed on an Agilent 1290 Infinity II UHPLC-MS system equipped with a G7117 Diode Array Detector and a LC/MSD XT G6135 Single Quadrupole Mass Detector. Chromatography performed with the instrument was on a Waters Atlantis T3 C18 column (4.6 × 150 mm, 3 µm particle size) and operated at a flow rate of 0.5 mL/min with a binary gradient mobile phase consisting of 10 mM ammonium acetate (pH 4.5) and methanol. The course of chromatography was monitored at 260 nm. Mass spectrometry was operated in both positive (+ESI) and negative (-ESI) electrospray ionization modes. MS was performed with a capillary voltage of 2500 V at both modes, a fragmentor voltage of 70 V, and a mass range of m/z 100 to 1000. Agilent ChemStation software was used for the primary LC-MS data processing. ChemStation produced chromatograms were further annotated in Adobe Illustrator.

**Northern blot with APB-acrylamide gel**

Overnight *E. coli* cultures were diluted 1/100-fold into 5 mL of LB supplemented with 0.4 % arabinose and 100 µg/mL ampicillin and grown for 2 h at 37 °C. Cells were harvested by centrifugation at 16,000 x g for 2 min at 4 °C. Cell pellets were immediately resuspended in 1 mL of Trizol (Life technologies, Carlsbad, CA). Small RNAs were extracted using PureLinkTM miRNA Isolation kit from Invitrogen (Carlsbad, CA) according to manufacturer protocol. Purified small RNAs were eluted in 50 µL of RNase-free water and concentrations were measured by NanoDrop® ND-1000 Spectrophotometer (Thermo Fisher Scientific, Waltham, MA). Then, 200 µg were boiled in 2X RNA Loading Dye (NEB, Ipswich, MA), then migrated on a 0.5 % 3-(Acrylamido)-phenylboronic acid (APB from Sigma-Aldrich, St. Louis, MO), 10 % acrylamide/bisacrylamide (29:1), 8 M Urea, 1X TAE (40 mM Tris-HCl, 20 mM EDTA, 1 mM acetate, pH 7.5). Migrated samples were transferred onto a ByodyneTM B Pre-Cut Modified Nylon Membrane, 0.45 µm (Thermo Fisher Scientific, Waltham, MA). The RNAs were cross-linked using UV and the Chemiluminescent Nucleic Acid Detection Module (Thermo Fisher Scientific, Waltham, MA) was used, replacing the first blocking buffer by the DIG Easy Hyb buffer (Sigma-Aldrich, St. Louis, MO), with the (5'-biotin-CCCTCGGTGACAGGCAGG-3') probe that detects $tRNA_{Asp}$(GUC) at final concentration of 0.3 µM. Revelation was performed using the iBright FL1000 device (Invitrogen, Carlsbad, CA) in automatic detection mode.

#### Structure flexibility prediction

Medusa (10) online server (https://www.dsimb.inserm.fr/MEDUSA/) was used to predict the flexibility of proteins. The strict threshold result was used and transformed into a confidence score of flexibility from 0, confidently rigid, to 100, confidently flexible, following this calculation: if the residue is predicted rigid, the confidence score was inverted and multiply by 50 ((1 - score) x 50); if the residue is predicted flexible, the confidence score was multiplied by 50 to which 50 was added (50 + score x 50). The result was viewed on ChimeraX version: 1.5rc202210241843 (11) and colored by attribute on a Blue-Red scale.

**Supplementary Alignments**

**Alignment S1 : QueF**

```
E.coli             -MS------SYANHQALAGLT-LG-KSTDY--------RDTYDASLLQGVPRSLNRDPLG
MD-2021a           -MS------VEQSL--------LG-KETQY--------PTSYQPDVLFPIARAQSREKYA
vB_FspM_immuto_    -MAEL----NQAEVVKIAGKH-LG-KVGGEGY------KDTYDPSLLVEIPRYLNREAYG
pf16               -MS------NQSEIRKLAGVH-LG-KAGDGSVVNPYQTPDAIDPSLLVAVPRHLNRTDYN
phage2             -MN------NQEELNKLVGVH-LG-KAGDGSAVKPYVTPDEVDPSLLVAVPRHLNRTGYG
phage3             -MT---------KIEDIASVH-LG-KAGDGSVVKPYVTPDSVDPSLLVGVPRVLNRTAYE
1.081.O._10N.28    -MSE-----NYDKVSAIAAEH-LGKKAGDYSD---FVDIKDVKSDLLVAIPRELNRVDYN
KVP40              -MS------NYDQVSAIAAEH-LGKKAGDYSD---FVDIVEVRPDLLVPIPRELNRTDYG
phi-pp2            -MS------NYDQVSAIAAEH-LGKKAGDYSD---FVDIVEVRPDLLVPIPRELNRTDYG
phi-Grn1           -MS------NYDEVSAIAAEH-LGKKAGDYSD---FVNIVDVRPDLLVPIPRELNRTDYG
ValKK3             -MS------NYDEVSAIAAEH-LGKKAGDYSD---FVNIVDVRPDLLVPIPRELNRTDYG
phi-ST2            MTS------NYDEVSAIAAEH-LGKKAGDYSD---FVNIVDVRPDLLVPIPRELNRTDYG
Va3                -MS------NYDEVSAIAAEH-LGKKAGDYSD---FVNIVDVRPDLLVPIPRELNRTDYG
vB_ValM_R10Z       -MS------NYDEVSAIAAEH-LGKKAGDYSD---FVNIVDVRPDLLVPIPRELNRTDYG
VH7D               -MS------NYDEVSAIAAEH-LGKKAGDYSD---FVNIVDVRPDLLVPIPRELNRTDYG
phage1             -MS--------TNLTDIAAKT-LG-SSASYAV-----YTEQFDPSLLNPMPRHLARDGWG
B.subtilis         -MT-------TRKESELEGVTLLGNQGTNYLF--------EYAPDVLESFPN--------
Dp-1               -MS----QNTTRTDAELTGVTLLGNQDTKYDY--------DYNPDVLETFPN--------
vB_DshS-R5C        -MT--------TEYSKPDDLEKLGSGSADQRE-WP---KLPPTPEVLERFPNPMK-----
phi3:1             -MDPAVNQEASGKVFAFDGVEKIMTQE---------------------------------
phi3ST:2           -MDPAVNQEASGKVFAFDGVEKIMTQE---------------------------------
phi47:1            -MDPAVNQEASGKVFAFDGVEKIMTQE---------------------------------
phiSM              -MDPAVNQEASGKVFAFDGVEKIMTQE---------------------------------
Knocker            -MSDHNGVLGRTHTGTLESVETLAISVVN-------------TPDVVSA-----------
P.calidifontis     -ML-----------------------------------KVSKSPSLVRLKTR--------
HCTV-16            -MAD---------------------NEVTFRF-NP--DYDPIDSDVLQAVPN--------
HCTV-1             -MAD---------------------NEVTFRF-NP--DYDPIDSDVLQAVPN--------
HCTV-5             -MGD---------------------NEVTFRF-NP--DYDPINGEVLQAVPN--------
HVTV-1             -MGD---------------------NEVNFRF-NP--DYDPINSEVLQAVPN--------

E.coli             LKADNLPFHGTDIWTLYELSWLNAKGLPQVAVGHVELDYTSVNLIESKSFKLYLNSFNQT
MD-2021a           HIEGI--TQGKDWWHVFEISWLNAHGIPQVAIGRITLPASSPNLIESKSLKLYFNSLNFT
vB_FspM_immuto_    IDDNNLPFVGGDVWNAYEVSAITTKGLPVVGMLKIYYPADSRLHVESKSIKLYLNSFNMT
pf16               IAEDNLPFIGFDVWNCYEVSALTTNGYPVSGVLKIKYPANSPSIVESKSLKLFLNSFNMD
phage2             IQEEELPFVGIDAWNSYEFSTLQKNGFPISGWLKFTYSSSTPNIVESKSVKLYLNSYNMA
phage3             IDEENLPFEGSDTWNCYEFSTLANNGFPVSGLLRIVYPSDSANIVESKSLKLYLNSFNMM
1.081.O._10N.28    IDNDT--FVGIDAWHGFEVSALTKNGLPVSGMVKVVYPSNSPNIVESKSMKLYWNSFNMA
KVP40              ITNED--FAGFDTWHGFEVSALTEQGLPVSGMAKVVYPSDSPNIVESKSMKLYWNSFNMA
phi-pp2            ITNED--FAGFDTWHGFEVSALTEQGLPVSGMAKVVYPSDSPNIVESKSMKLYWNSFNMA
phi-Grn1           ITNED--FAGFDTWHGFEVSALTEQGLPVSGMAKVVYPSDSPNIVESKSMKLYWNSFNMA
ValKK3             ITNED--FAGFDTWHGFEVSALTEQGLPVSGMAKVVYPSDSPNIVESKSMKLYWNSFNMA
phi-ST2            ITNED--FAGFDTWHGFEVSALTEQGLPVSGMAKVVYPSDSPNIVESKSMKLYWNSFNMA
Va3                ITNED--FAGFDTWHGFEVSALTEQGLPVSGMAKVVYPSDSPNIVESKSMKLYWNSFNMA
vB_ValM_R10Z       ITNED--FAGFDTWHGFEVSALTEQGLPVSGMAKVVYPSDSPNIVESKSMKLYWNSFNMA
VH7D               ITNED--FAGFDTWHGFEVSALTEQGLPVSGMAKVVYPSDSPNIVESKSMKLYWNSFNMA
phage1             IKSDM--FVGFDTWHCHEATFLLNNGVPVAGTVKYVYPADSEFMVESKSAKLYMNSFDMC
B.subtilis         ------------------------------------------------------------
Dp-1               ------------------------------------------------------------
vB_DshS-R5C        ------------------------------------------------------------
phi3:1             ------------------------------------------------------------
phi3ST:2           ------------------------------------------------------------
phi47:1            ------------------------------------------------------------
phiSM              ------------------------------------------------------------
```

```
Knocker         ------------------------------------------------------------
P.calidifontis  ------------------------------------------------------------
HCTV-16         ------------------------------------------------------------
HCTV-1          ------------------------------------------------------------
HCTV-5          ------------------------------------------------------------
HVTV-1          ------------------------------------------------------------

E.coli          RFNNWDE-----VRQTLERDLSTCAQGK-ISVALYRL-DELEGQPIGHFNG------TCI
MD-2021a        QFDSTQS-----FIKTVEKDLSAAAGAK-VELTLFQV-DDLEISK---PQG------ICI
vB_FspM_immuto_ KMGDTAAECIAILKDRVKRDLSEKLETE-VGVEMFTS-DFGPAYA---FKG-YAQLDQMV
pf16            RLADSVDDVIEVIQQTVETHLGELLDCA-VSVKFFAYWDSASSKKN-HLHEEYRRIEHGI
phage2          RLIESKEELWK-IEEQVERDIAKAVGGE-VGVYIAIG-DVDTVKP---MKGDFMSLENYC
phage3          KNGDTIDEILINIEERIYEDLSKCLETESVDVCLLFE-DGITRKP---VEGDFVQVERFV
1.081.O._10N.28 KIADNSRDIRAEIERIATADLSECMQAP-VQVTFFTE-RVGQPHG---MNG-FAEMSKYL
KVP40           KVAATRAEVKDAIEAIASRDLSEVIGAP-VQVSFFTE-KLTQPHG---MTG-FAELDKYV
phi-pp2         KVAATRAEVKDAIEAIASRDLSEVIGAP-VQVSFFTE-KLTQPHG---MTG-FAELDKYV
phi-Grn1        KVAATRAEVQGAIEAIASRDLSEVIGAE-VQVSFFTI-RSTQPHG---MTG-YVELDKFV
ValKK3          KVAATRAEVQGAIEAIASRDLSEVIGAE-VQVSFFTI-RSTQPHG---MTG-YVELDKFV
phi-ST2         KVAATRAEVQGAIEAIASRDLSEVIGAE-VQVSFFTV-RSTQPHG---MTG-YAELDKYV
Va3             KVAATRAEVQGAIEAIASRDLSEVIGAE-VQVSFFTV-RSTQPHG---MTG-YAELDKYV
vB_ValM_R10Z    KVAATRAEVQGAIEAIASRDLSEVIGAE-VQVSFFTV-RSTQPHG---MTG-YAELDKYV
VH7D            KVAATRAEVQGAIEAIASRDLSEVIGAE-VQVSFFTV-RSTQPHG---MTG-YAELDKYV
phage1          KMGDTVDEAIKNYENQIATDLTNVIGKE-VKVKFFKS-GSVGLFP---LND-YTDLYDIV
B.subtilis      ------------------------------------------------------------
Dp-1            ------------------------------------------------------------
vB_DshS-R5C     ------------------------------------------------------------
phi3:1          ------------------------------------------------------------
phi3ST:2        ------------------------------------------------------------
phi47:1         ------------------------------------------------------------
phiSM           ------------------------------------------------------------
Knocker         ------------------------------------------------------------
P.calidifontis  --------------------------------------------------G---------
HCTV-16         ------------------------------------------------------------
HCTV-1          ------------------------------------------------------------
HCTV-5          ------------------------------------------------------------
HVTV-1          ------------------------------------------------------------

E.coli          D-DQDITIDNYEFTTDYLENA---TCGEKVVEETLVSHLLKSNCLI--------------
MD-2021a        D-DLMPERLERHPDATLLKLDE----SGEEIEVELYSHLLRSNCPV--------------
vB_FspM_immuto_ D-LDAVEFTSYHSDASQLQCEE--TDDDTQFEIKFQSNLLRSNCRV--------------
pf16            H-MQSVVFNDFNENPGLLEVSPGVSTSSVQ---RITTNALRSNCQI--------------
phage2          N-VDKMSFDRYNESSDILEVVP--SIGRYE---RWRSHSLRSNCRV--------------
phage3          D-PATITFDQFNESPDTLQVVD--SDGYPK---LWRSAVLRSNCRV--------------
1.081.O._10N.28 S-PDLVFDADGADTGLLVSA----GRGGQL---KVMTNNLRSNCRV--------------
KVP40           S-ESTVFDSESGADADLLQE----SHIKKL---KVFTPNLRSNCRV--------------
phi-pp2         S-ESTVFDSESGADADLLQE----SHIKKL---KVFTPNLRSNCRV--------------
phi-Grn1        S-EDTVFDSESGADADLLQE----SHVKRM---KVFTPNLRSNCRV--------------
ValKK3          S-EDTVFDSESGADADLLQE----SHVKRM---KVFTPNLRSNCRV--------------
phi-ST2         S-EDTVFDSESGADADLLQE----SHVKRM---KVFTPNLRSNCRV--------------
Va3             S-EDTVFDSESGADADLLQE----SHVKRM---KVFTPNLRSNCRV--------------
vB_ValM_R10Z    S-EDTVFDSESGADADLLQE----SHVKRM---KVFTPNLRSNCRV--------------
VH7D            S-EDTVFDSESGADADLLQE----SHVKRM---KVFTPNLRSNCRV--------------
phage1          NKTRNIEITDYSAKENHLKFVSR-PDGGTLCDNKYFTNALRSRCRH--------------
B.subtilis      ----------KHVNRDYF--------------VKFNCPEFTSLCPK--------------
Dp-1            ----------KHPENNYL--------------VTFDGYEFTSLCPK--------------
```

```
vB_DshS-R5C     --------FSRHPSPNPIQL------GGYD-----DPFEFTSLCPK--------------
phi3:1          -----LKTFDFDSSEQLIET---------------FTPEFSAVCPF--------------
phi3ST:2        -----LKTFDFDSSEQLIET---------------FTPEFSAVCPF--------------
phi47:1         -----LKTFDFDSSEQLIET---------------FTPEFSAVCPF--------------
phiSM           -----LKTFDFDSSEQLIET---------------FTPEFSAVCPF--------------
Knocker         -----------------------------------RCTEVQACCPV--------------
P.calidifontis  ----------------------------------------ESVCPI--------------
HCTV-16         ----------PRPDTNRTDRH--------------VAKEFSTNCPVDYGVDPDADDDEVE
HCTV-1          ----------PRPDTNRTDRH--------------VAKEFSTNCPVDYGVDPDADDDEVE
HCTV-5          ----------PRPDTDRTDRH--------------VAKEFSTNCPVDYGVDPDADEDEVE
HVTV-1          ----------PRPDTERTDRH--------------VAKEFSTNCPVDYGVDPDAEDDEVE
                                                         : *

E.coli          -THQPDWGSLQIQYRGR---QIDREKLLRYLVSFRHHNEFHEQCVERIFNDLLRFC-QPE
MD-2021a        -TGQPDWGTVFIRFKGK---KPCYRSLLAYIISYRQHNGFHEQCVEQIFADIWQNL-QPE
vB_FspM_immuto_ -TNQPDWGDVFVHIKPAAGKIPNLESIARYIVSHRQVSHFHEEICEMIYTHLTQAY-QPE
pf16            -TNQPDWGDIYIFVKGP--SQLVERSVLQYIVSMRNESHFHEEIVECVFKRLYDLLPRGT
phage2          -TNQPDWGDVYIHIKGE--KAVTPESLLQYIVSMRKENHFHEEIAECIYKRLWDLL-EPE
phage3          -TNQPDWGDVYIAIKGD--KTVTPESLLQYIVSMRKENHFHEEICECIYKRLHDLI-NPD
1.081.O._10N.28 -THQPDFGDLYIDMIGT--DVPTIDSLMQYIVSFRKENHFHEECVEMIYKALQDTF-EPS
KVP40           -THQPDFGDLYVYMSGE--KTPTVDSLMQYIVSFRKENHFHEECVEMIYKALLDKF-DPT
phi-pp2         -THQPDFGDLYVYMSGE--KTPTVDSLMQYIVSFRKENHFHEECVEMIYKALLDKF-DPT
phi-Grn1        -THQPDFGDLYVYMKGD--KTPTIDSLMQYIVSFRKENHFHEECVEMIYKALLDKF-DPQ
ValKK3          -THQPDFGDLYVYMKGD--KTPTIDSLMQYIVSFRKENHFHEECVEMIYKALLDKF-DPQ
phi-ST2         -THQPDFGDLYVYMKGE--KTPTIDSLMQYIVSFRKENHFHEECVEMIYKALLDKF-DPQ
Va3             -THQPDFGDLYVYMKGE--KTPTIDSLMQYIVSFRKENHFHEECVEMIYKALLDKF-DPQ
vB_ValM_R10Z    -THQPDFGDLYVYMKGE--KTPTIDSLMQYIVSFRKENHFHEECVEMIYKALLDKF-DPQ
VH7D            -THQPDFGDLYVYMKGE--KTPTIDSLMQYIVSFRKENHFHEECVEMIYKALLDKF-DPQ
phage1          -TKQKDTGAAYINIITK-GTSVDPISLFKQIVSLREVNEFHEFCAEKLLTSIMEHE-EVV
B.subtilis      -TGQPDFATIYISYIPD-EKMVESKSLKLYLFSFRNHGDFHEDCMNIIMNDLIELM-DPR
Dp-1            -TGQPDFANVFISYIPN-EKMVESKSLKLYLFSFRNHGDFHEDCMNIILNDLYELM-EPK
vB_DshS-R5C     -TGQPDHARIVVNYQPK-DWCVESKSWKIYLQSFRLHGEFHESCIQRIKDDLVDLL-DPL
phi3:1          -SGLPDIAELTIVYRPRGGKCIELKSLKYYLTSFRNVGLYQEGCTKTIYQDLAKIL-ETD
phi3ST:2        -SGLPDIAELTIVYRPRGGKCIELKSLKYYLTSFRNVGLYQEGCTKTIYQDLAKIL-ETD
phi47:1         -SGLPDIAELTIVYRPRGGKCIELKSLKYYLTSFRNVGLYQEGCTKTIYQDLAKIL-ETD
phiSM           -SGLPDIAELTIVYRPRGGKCIELKSLKYYLTSFRNVGLYQEGCTKTIYQDLAKIL-ETD
Knocker         -TLQPDLYTVVIEYRPTTGEVIESKSLKLYLWKFRDRGISCEELAATIANELSRAV-QSP
P.calidifontis  -SKTVDSFEVSVEYIPR-GAVLAIEEFKKMVDSYRGREILHEELAVDLLEKVKAAV-NPP
HCTV-16         SEGVRDYGEIEIEYRPD-DFIVELKSLKYYLMSFEDARISHEEVTAKVFQDLAGIL-YPD
HCTV-1          SEGVRDYGEIEIEYRPD-DFIVELKSLKYYLMSFEDARISHEEVTAKVFQDLAEIL-YPD
HCTV-5          SEGVRDYGEIEIEYVPD-DFIVELKSLKYYLMSFEDARISHEEVTAKVWQDLAEIL-YPD
HVTV-1          SEGVRDYGEIEIEYVPE-DYIVELKSLKYYLMSFEDARISHEEVTAKVWQDLAEIL-YPD
                     *     :             .    : . .      *     :   :

E.coli          K--------LSVYARYTRRGGLDINPWRSNSDF-VPST------------TRLVRQ----
MD-2021a        K--------LMVYATYTRRGGLDINPCRVSDLTWMPKP------------IRLARQ----
vB_FspM_immuto_ E--------LMVACLYTRRGGLDINPVRASHKHLIPSF----FADPKCRMEKTLRQ----
pf16            D--------IAVGALYTRRGGIDINPVRATSFNALDEF--AQLTDINTMQFKTARQ----
phage2          E--------LLVTCLYTRRGGIDINPTRASNYPLLNEA---PIIDAYNFCEKTARQ----
phage3          E--------LFVACLYTRRGGIDINPLRASSREVMYKY-GSGLTDPYSINFKTLRQ----
1.081.O._10N.28 G--------LMVTALYTRRGGWNICPSRASHESLLPDT-LKSASVSHTNGSCLFRQ----
KVP40           E--------LMVTALYTRRGGWNICPARATHEELLPEA-LGSKSVSHTNGSCLFRQ----
phi-pp2         E--------LMVTALYTRRGGWNICPARATHEELLPEA-LGSKSVSHTNGSCLFRQ----
phi-Grn1        E--------LMVTALYTRRGGWNICPARATHEELLPEA-LGSEVVSHTNGSCLFRQ----
ValKK3          E--------LMVTALYTRRGGWNICPARATHEELLPEA-LGSEVVSHTNGSCLFRQ----
phi-ST2         E--------LMVTALYTRRGGWNICPARATHEELLPEA-LGSEVVSHTNGSCLFRQ----
Va3             E--------LMVTALYTRRGGWNICPARATHEELLPEA-LGSEVVSHTNGSCLFRQ----
```

```
vB_ValM_R10Z    E--------LMVTALYTRRGGWNICPARATHEELLPEA-LGSEVVSHTNGSCLFRQ----
VH7D            E--------LMVTALYTRRGGWNICPARATHEELLPEA-LGSEVVSHTNGSCLFRQ----
phage1          D--------CVVTLLYSRRGSLDINPSRASNKHLLPKL----LKDTNVYTEKAMGQ----
B.subtilis      Y--------IEVWGKFTPRGGISIDPYTNYGKP--GTK-YEKMAEYRMMNHDLYPETIDN
Dp-1            Y--------IEVMGLFTPRGGISIYPFVNKVNPQFATPELEQLQLQRKLNF------LGN
vB_DshS-R5C     W--------IEVTGEFTPRGGISINPTARWEYPMKGVT-FEQMTDTRSPQDRLNEM-LAN
phi3:1          A--------IIVKTKYNVRGGFSTTCQEGK------------------------------
phi3ST:2        A--------IIVKTKYNVRGGFSTTCQEGK------------------------------
phi47:1         A--------IIVKTKYNVRGGFSTTCQEGK------------------------------
phiSM           A--------IIVKTKYNVRGGFSTTCQEGK------------------------------
Knocker         ---------VTVTAHQASRGGIELSATRKGSVLE--------------------------
P.calidifontis  Y--------VKVTVK-SYYIGVEVEVVAESGGVP--------------------------
HCTV-16         RTEAQAKELLHVRFDVNPRGGISSDTWVGGAR----------------------------
HCTV-1          RTEAQAKELLHVRFDVNPRGGISSDTWVGGAR----------------------------
HCTV-5          LDEDEVQKRLHVAFDVNPRGGISSQTWVGGIQ----------------------------
HVTV-1          LDEDEAQKRLHVAFDVNPRGGISSQTWVGGIQ----------------------------
                           *        . .

E.coli          -------------------
MD-2021a        -------------------
vB_FspM_immuto_ -------------------
pf16            -------------------
phage2          -------------------
phage3          -------------------
1.081.O._10N.28 -------------------
KVP40           -------------------
phi-pp2         -------------------
phi-Grn1        -------------------
ValKK3          -------------------
phi-ST2         -------------------
Va3             -------------------
vB_ValM_R10Z    -------------------
VH7D            -------------------
phage1          -------------------
B.subtilis      --------------R----
Dp-1            VQGLGRAI------R----
vB_DshS-R5C     GSHVGERIGEGEMPDPT-M
phi3:1          ------------------L
phi3ST:2        ------------------L
phi47:1         ------------------L
phiSM           ------------------L
Knocker         ---------------P---
P.calidifontis  ---------------PVYI
HCTV-16         -------------------
HCTV-1          -------------------
HCTV-5          -------------------
HVTV-1          -------------------
```

**Alignment S2: DpdM**

```
phiSM           MAQRDSLNKI-DDYIKAVNNE------------QLILSPMFFRDYSCPSHCGGCCPKFSL
Va3             M-KVDSVDKI-AMYMSMVSKTPFRYKGNLYVPKTMRIQENIRKGFLCPASCGSCCGRWSL
VH7D            M-KVDSVDKI-AMYMSMVSKTPFRYKGNLYVPKTMRIQENIRKGFLCPASCGSCCGRWSL
vB_ValM_R10Z    M-KVDSVDKI-AMYMSMVSKTPFRYKGNLYVPKTMRIQENIRKGFLCPASCGSCCGRWSL
phi-ST2         -----------------------------------------------------------M
phi-ST2_correct V-KVDSVDKI-AMYMSMVSKTPFRYKGNLYVPKTMRIQENIRKGFLCPASCGSCCGRWSL
phi-Grn1        M-KVDSVDKI-AMYMSMVSKNPFRYKGNLYVPKTMRIQENIRKGFLCPASCGSCCGRWSL
```

```
Knocker         M-KNDSIDKVVESYFATVTAEPFTYKGKRFDVPDLIVSPLILRGFTCPAVCGGCCRNWSL
                                                           :

phiSM           DYFEGERWEKFK----ETYPHLVHK---FKKRI--VNGATVYTNYQTENPD--------S
Va3             DYLPNDYLDEHDTHGVETRAPLEAQSDLYEKRVINVNGKDVVLLSDTQEKDYTNRTLKPN
VH7D            DYLPNDYLDEHDTHGVETRAPLEAQSDLYEKRVINVNGKDVVLLSDTQEKDYTNRTLKPN
vB_ValM_R10Z    DYLPDDYLDEHDTHGVETRAPLEAQSDLYEKRVINVNGKDVVLLSDTQEKDYTNRTLKPN
phi-ST2         DYLPNDYLDEHDTYGVETRAPLEAQSDLYEKRVINVNGKDVVLLSDTQEKDYTNRTLKPN
phi-ST2_correct DYLPNDYLDEHDTYGVETRAPLEAQSDLYEKRVINVNGKDVVLLSDTQEKDYTNRTLKPN
phi-Grn1        DYLPNDYLDEHDTHGVETRAPLETQSDLYEKRVINVNGKDVVLLSDTQEKDYTNRTLKPN
Knocker         DYLPGE-----------------PQPPTTVPREVEFDGRAVLVRSIAADGD------PAD
                **: .:                  :      *   .:*  *     : : *        .

phiSM           KCEFLNKEDGRCGIHVS-------NPFSCEFELNKFLV------------------KKDK
Va3             ACRNVSM-TGRCDIHNFVLQGDYGQPFSCDFEIIRFISPAGHAQGHVTEKQFKEGIRYSI
VH7D            ACRNVSM-TGRCDIHNFVLQGDYGQPFSCDFEIIRFISPAGHAQGHVTEKQFKEGIRYSI
vB_ValM_R10Z    ACRNVSM-TGRCDIHNFVLQGDYGQPFSCDFEIIRFISPAGHAQGHVTEKQFKEGIRYSI
phi-ST2         ACRNVSM-TGRCDIHNFVLQGDYGQPFSCDFEIIRFISPAGHAQGHVTEKQFKEGIRYSI
phi-ST2_correct ACRNVSM-TGRCDIHNFVLQGDYGQPFSCDFEIIRFISPAGHAQGHVTEKQFKEGIRYSI
phi-Grn1        ACRNVSM-TGRCDIHNFVLQGDYGQPFSCDFEIIRFISPAGHAQGHVTEKQFKEGIRYSI
Knocker         FCRQLDMATGRCGIHGV-------HPFTCDFELIRSMR------------------RGSQ
                 *. :.   ***.**         :**:*:**: : :                   : .

phiSM           TYLMNKLFGRGWAMMRIDGE--------------------RGALCEMKDF------NLKK
Va3             GRITTDLYANDRHMRDVHGKWREMKSLSHPDFIDVKHVDLPGPKCKLTDVDEDSVADAIR
VH7D            GRITTDLYANDRHMRDVHGKWREMKSLSHPDFIDVKHVDLPGPKCKLTDVDEDSVADAIR
vB_ValM_R10Z    GRITTDLYANDRHMRDVHGKWREMKSLSHPDFIDVKHVDLPGPKCKLTDVDEDSVADAIR
phi-ST2         GRITTDLYANDRHMRDVHGKWREMKSLSHPDFIDVKHVDLPGPKCKLTDVDEDSVADAIR
phi-ST2_correct GRITTDLYANDRHMRDVHGKWREMKSLSHPDFIDVKHVDLPGPKCKLTDVDEDSVADAIR
phi-Grn1        GRITTDLYANDRHMRDVHGKWREMKSLSHPDFIDVKHVDLPGPKCKLTDVDEDSVADAIR
Knocker         ARLNQQLFTRGWAMRRVDGE--------------------RGARCSMTPPTPETVADVRR
                  :  .*: ..  *  :.*:                     *. *.:.        :  :

phiSM           YLRDVELLQELLDIVSKFNKKAPLLEK------VV-----SFLKKTNRPIPKQKVFTINE
Va3             KFRRLQVWSDYFGIETWLDEIIDWLETKKFGSFVVNAQPQTKYKESDTGYTNMKPYELSD
VH7D            KFRRLQVWSDYFGIETWLDEIIDWLETKKFGSFVVNAQPQTKFKESDTGYTNMKPYELSD
vB_ValM_R10Z    KFRRLQVWSDYFGIETWLDEIIDWLETKKFGSFVVNAQPQTKYKESDTGYTNMKPYNLSD
phi-ST2         KFRRLQVWSDYFGIETWLDEIIDWLETKKFGSFVVNAQPQTKYKESDTGYTNMKPYNLSD
phi-ST2_correct KFRRLQVWSDYFGIETWLDEIIDWLETKKFGSFVVNAQPQTKYKESDTGYTNMKPYNLSD
phi-Grn1        KFRRLQVWSDYFGIETWLDEIIDWLETKKFGSFVVNAQPQTKYKESDTGYTNMKAYNLSD
Knocker         KLRRLKQWADHFGLRTRVPTILAWIDV-------------TAIDFADSAALRLPAS----
                 :* ::   : :.: : .      ::              :  . ::    .

phiSM           EKDNQVNLF
Va3             YEKNEDN--
VH7D            YEKNEDN--
vB_ValM_R10Z    YEKNEDN--
phi-ST2         YEKNEDN--
phi-ST2_correct YEKNEDN--
phi-Grn1        YEKNEDK--
Knocker         ---------
```

**Alignment S3: DpdL**

```
NP_417245.1     ------------------------------------------------------------
NP_389256.1     M-----------------------------------------------------------
YP_007673448.1  ------------------------------------------------------------
```

```
CAB4159940.1     ------------------------------------------------------------
CAB4155266.1     ------------------------------------------------------------
CAB4125606.1     ------------------------------------------------------------
CAB4142608.1     ------------------------------------------------------------
QDP63658.1       ------------------------------------------------------------
QDP49444.1       M-------------------------------------------------------ISAH
CAB4128316.1     ------------------------------------------------------------
CAB5215170.1     ------------------------------------------------------------
CAB4137873.1     ------------------------------------------------------------
CAB4241859.1     ------------------------------------------------------------
CAB4125953.1     ------------------------------------------------------------
CAB4221099.1     M-----------------------------------------------------------
CAB4163928.1     ------------------------------------------------------------
CAB4123241.1     ------------------------------------------------------------
CAB4140455.1     ------------------------------------------------------------
CAB5220926.1     ------------------------------------------------------------
QIG70625.1       ------------------------------------------------------------
YP_009609725.1   ------------------------------------------------------------
QCG76199.1       ------------------------------------------------------------
CAB4130199.1     MKVRVLQTTGNGSFTEVEWEKPEITADEIEVASVMTGVCRSDIDMMNGKFGPLPVAMQGH
USV40849.1       ------------------------------------------------------------
QRE00068.1       ------------------------------------------------------------
QIW86898.1       ------------------------------------------------------------
CAB4149958.1     ------------------------------------------------------------
CAB4127312.1     ------------------------------------------------------------
AGH13910.1       ------------------------------------------------------------

NP_417245.1      ------------------------------------------------------------
NP_389256.1      ------------------------------------------------------------
YP_007673448.1   ------------------------------------------------------------
CAB4159940.1     ------------------------------------------------------------
CAB4155266.1     ------------------------------------------------------------
CAB4125606.1     ------------------------------------------------------------
CAB4142608.1     ------------------------------------------------------------
QDP63658.1       ------------------------------------------------------------
QDP49444.1       LG----------------------------------------------------------
CAB4128316.1     ------------------------------------------------------------
CAB5215170.1     ------------------------------------------------------------
CAB4137873.1     ------------------------------------------------------------
CAB4241859.1     ------------------------------------------------------------
CAB4125953.1     ------------------------------------------------------------
CAB4221099.1     ------------------------------------------------------------
CAB4163928.1     ------------------------------------------------------------
CAB4123241.1     ------------------------------------------------------------
CAB4140455.1     ------------------------------------------------------------
CAB5220926.1     ------------------------------------------------------------
QIG70625.1       ------------------------------------------------------------
YP_009609725.1   ------------------------------------------------------------
QCG76199.1       ------------------------------------------------------------
CAB4130199.1     EGLGKVTKVGANITNITVGAYVATRGEPAYADHYNVRRGEFVKVPSAEPKFILEPVACGI
USV40849.1       ------------------------------------------------------------
QRE00068.1       ------------------------------------------------------------
QIW86898.1       ------------------------------------------------------------
CAB4149958.1     ------------------------------------------------------------
CAB4127312.1     ------------------------------------------------------------
AGH13910.1       ------------------------------------------------------------
```

```
NP_417245.1         ------------------------------------------------------------
NP_389256.1         ------------------------------------------------------------
YP_007673448.1      ------------------------------------------------------------
CAB4159940.1        ------------------------------------------------------------
CAB4155266.1        ------------------------------------------------------------
CAB4125606.1        ------------------------------------------------------------
CAB4142608.1        ------------------------------------------------------------
QDP63658.1          ------------------------------------------------------------
QDP49444.1          ------------------------------------------------------------
CAB4128316.1        ------------------------------------------------------------
CAB5215170.1        ------------------------------------------------------------
CAB4137873.1        ------------------------------------------------------------
CAB4241859.1        ------------------------------------------------------------
CAB4125953.1        ------------------------------------------------------------
CAB4221099.1        ------------------------------------------------------------
CAB4163928.1        ------------------------------------------------------------
CAB4123241.1        ------------------------------------------------------------
CAB4140455.1        ------------------------------------------------------------
CAB5220926.1        ------------------------------------------------------------
QIG70625.1          ------------------------------------------------------------
YP_009609725.1      ------------------------------------------------------------
QCG76199.1          ------------------------------------------------------------
CAB4130199.1        NLIFQNKSAIAERAGAGKRMLILGSGFLAWTAYNTVKLSNYHYSIDVVGSSNTAMWLGAG
USV40849.1          ------------------------------------------------------------
QRE00068.1          ------------------------------------------------------------
QIW86898.1          ------------------------------------------------------------
CAB4149958.1        ------------------------------------------------------------
CAB4127312.1        ------------------------------------------------------------
AGH13910.1          ------------------------------------------------------------

NP_417245.1         ------------------------------------------------------------
NP_389256.1         ------------------------------------------------------------
YP_007673448.1      ------------------------------------------------------------
CAB4159940.1        ------------------------------------------------------------
CAB4155266.1        ------------------------------------------------------------
CAB4125606.1        ------------------------------------------------------------
CAB4142608.1        ------------------------------------------------------------
QDP63658.1          ------------------------------------------------------------
QDP49444.1          ------------------------------------------------------------
CAB4128316.1        ------------------------------------------------------------
CAB5215170.1        ------------------------------------------------------------
CAB4137873.1        ------------------------------------------------------------
CAB4241859.1        ------------------------------------------------------------
CAB4125953.1        ------------------------------------------------------------
CAB4221099.1        ------------------------------------------------------------
CAB4163928.1        ------------------------------------------------------------
CAB4123241.1        ------------------------------------------------------------
CAB4140455.1        ------------------------------------------------------------
CAB5220926.1        ------------------------------------------------------------
QIG70625.1          ------------------------------------------------------------
YP_009609725.1      ------------------------------------------------------------
QCG76199.1          ------------------------------------------------------------
CAB4130199.1        ESLFKEPNGSYDVVIDLSNSDIVFRGDCLNNNALVILAAEKHPKIVTDFSKLLWKSCTIS
USV40849.1          ------------------------------------------------------------
```

```
QRE00068.1      ------------------------------------------------------------
QIW86898.1      ------------------------------------------------------------
CAB4149958.1    ------------------------------------------------------------
CAB4127312.1    ------------------------------------------------------------
AGH13910.1      ------------------------------------------------------------

NP_417245.1     ------------------------------------------------------------
NP_389256.1     ------------------------------------------------------------
YP_007673448.1  ------------------------------------------------------------
CAB4159940.1    ------------------------------------------------------------
CAB4155266.1    ------------------------------------------------------------
CAB4125606.1    ------------------------------------------------------------
CAB4142608.1    ------------------------------------------------------------
QDP63658.1      ------------------------------------------------------------
QDP49444.1      ------------------------------------------------------------
CAB4128316.1    ------------------------------------------------------------
CAB5215170.1    ------------------------------------------------------------
CAB4137873.1    ------------------------------------------------------------
CAB4241859.1    ------------------------------------------------------------
CAB4125953.1    ------------------------------------------------------------
CAB4221099.1    ------------------------------------------------------------
CAB4163928.1    ------------------------------------------------------------
CAB4123241.1    ------------------------------------------------------------
CAB4140455.1    ------------------------------------------------------------
CAB5220926.1    ------------------------------------------------------------
QIG70625.1      ------------------------------------------------------------
YP_009609725.1  ------------------------------------------------------------
QCG76199.1      ------------------------------------------------------------
CAB4130199.1    CPSPRNSQFIESMEQAVTWVSSGQLVVDRFWSKGYNRDKDWQAAFSDGNLRIPDYGRGYI
USV40849.1      ------------------------------------------------------------
QRE00068.1      ------------------------------------------------------------
QIW86898.1      ------------------------------------------------------------
CAB4149958.1    ------------------------------------------------------------
CAB4127312.1    ------------------------------------------------------------
AGH13910.1      ------------------------------------------------------------

NP_417245.1     -----------------------MMSTTLFKDFTFEAAHRLPHVP-----------EGHK
NP_389256.1     ----------HKLLSQIYPQAQHPYSFELNKDMHISAAHFIPRE------------SAGA
YP_007673448.1  ----------------------MSVKKVVGITTKVEGFHNFPTAVEM-FGK-----KVEF
CAB4159940.1    ------------------------MKYSVVVTFSIEGFHNWPDAKD--IFP-----EVAF
CAB4155266.1    ------------------------MKYSVVVTFSIEGFHNWPDAKD--VFP-----EVAF
CAB4125606.1    ----------------------MSTKTTVIVKLAVDGCHNFPKAAE--LFP-----EVDF
CAB4142608.1    ------------------------MKKYIEVKLDIEGLHHWPDCN----LP-----HVDY
QDP63658.1      -------------------------MTNIIVKLQYAGVHCWADCP----LE-----EVKY
QDP49444.1      -----------------LFANLFAMKKLIILHFEIEGHHAWKDAPE----------NIWF
CAB4128316.1    ---------MNQEQRETIERVKDAAKRQIWVTFRKEGIHRYPAAATDPNLCTAGEYDVAF
CAB5215170.1    ---------MNQEQREQIERIKSSAHRQIWITFRKEGIHCYPAAATDPKLNTAGEYDVSF
CAB4137873.1    ---------MIQAEREQIERIKESAERKIWITFRKEGIHKYPAALTDPQLATNDEYDVSF
CAB4241859.1    -----------------------MSQDMIWVKFQKEGIHRYPQALTDPKLATGDEYDVSF
CAB4125953.1    ----------------MIIRQDVRPNKMVWCTFRREGIHKYPAALTDPTLATGDEYDVSF
CAB4221099.1    ------FTQTETALAEKRNRISDRAKRLIWVTFRKEGIHMYPAAATDPTLATGDEYDVSF
CAB4163928.1    ------MNQRELALLEQRTKTMNTAIRYIWVTFQKEGIHCYPAAATDPNLATGDEYDVSF
CAB4123241.1    -------------------MTTEPTARMIYVTFQKEGIHCYPAALEDPALA-----DVSF
CAB4140455.1    ---------------------MFNSGKSIWVTFQKEGIHLYPQAATDPALA-----SVSF
CAB5220926.1    ------------------------MDKRIWVKFAKEGIHKYPAALTDPNLE-----DVSF
```

```
QIG70625.1      ----------------MPLITPEAIYRQIWVKFQRAGIHRYPAAATDPKLA-----DVAF
YP_009609725.1  -------------------------MKKIFVTFQKAGIHKYPAAAHDPKLS-----DVSY
QCG76199.1      ----------------------MGIKRWIEVPFQKEGIHMYPGADNNPALATNDWKDVSF
CAB4130199.1    YWEKKMEEANKKQVRDVITKNIKSARRWIWVTFQKVGFHKYPAAASEASLH-----DVRY
USV40849.1      -------------------------MKKIWVTFRKAGFHRYPAAGTQ-GLE-----DVAY
QRE00068.1      --------------------MLAHNPKFIYVTFQRKGYHFYPNAPE----------DVAY
QIW86898.1      ------------------MNQVFKKKAFVYCSFQLEGYHMYPGADVNPSTATGDYLDVSH
CAB4149958.1    -----------------MSPSRLNVKNFIEVRFQFEGIHHWPEAPR----------EVDF
CAB4127312.1    -----------------------MFSTFIKVRAEFEGFHFYPNASS--IDP-----RIKF
AGH13910.1      ------------------------MDNFIVCNVTVEGFHYWEDAPA----------QYAY
                                            :       . *

NP_417245.1     CGRLHGHSFMVRLEITGEVDPHTGWIIDFAELKAAFKPTYERLDHHYLND------IPGL
NP_389256.1     CSRVHGHTYTVNITVAGDELDDSGFLVNFSVLKKLVHGNYD---HTLLNDHEDFSQDDRY
YP_007673448.1  LSLNHRHLFGINLEIPVEHNERD---KEFILLKREVEEFIE---QRWGR-------PAQF
CAB4159940.1    LSDRHRHMFGFRCYATVTHTDRD---EEFILLNRKIQKALR---IAFVKAQPN---VLEF
CAB4155266.1    LSDRHRHMFGFRCYATVTHTDRD---EEFILLNRKIQKSLR---IGFSKESTN---VLEF
CAB4125606.1    LADRHRHMFHFTVACAVRHDDRD---KEFIMLKRDIITYIN---TYYFDTQTR---TCEF
CAB4142608.1    LAHLHRHTFVINCRAEVSHGDRD---IEFIDFKHKIKQYVA---KTWYDPAYG---CCNF
QDP63658.1      LKDMHRHTFYITCKKEVSHDDRD---IEIIMFKNKILQWLD---KTY---------KSNF
QDP49444.1      LSKEHRHLFTIRIGMNVEHNDRE---KEIFLQQQLFQEYLK---TKYEKTHLNDYTYHDF
CAB4128316.1    LASPHRHIFHFRVSIDVFHNDRD---IEFIQFKRWLENLYS---GTGPYNENR---VLEL
CAB5215170.1    LASPHRHIFHFRVSIDVFHNDRD---IEFIQFKRWCESLYS---SNSS-SQGS---VLEL
CAB4137873.1    LGYPHRHIFHFRVWIDVFHNDRD---IEFIQFKRWLENLYK----------DS---VLTL
CAB4241859.1    LGHPHRHIFHFKVYLEVFHDDRD---VEFIQFKRWLENLYN----------KG---TLEL
CAB4125953.1    LGYPHRHVFHFKVWISVEHNDRD---IEFIQFKRWLKSLYG-----------T---TLDL
CAB4221099.1    LGFPHRHIFHFKIAIQVFHNDRD---IEFIQFKRWIENLYK----------DS----LQL
CAB4163928.1    LASKHRHMFHFKVSISVTHNDRD---LEFIQAKRWLENLYK----------DS---ILAL
CAB4123241.1    LGYPHRHMFHFRVAISVLHHDRE---IEFILFKRWLESLYS----------DK---VLEL
CAB4140455.1    LGYPHRHIFHFRVELEVFHDDRD---VEFIMFKRELEQLYG----------DG---ILDL
CAB5220926.1    LGYPHRHMFHFYVEIEVVHNDRD---IEFIQFKRWLESLYD----------EK---VIEL
QIG70625.1      LAHPHRHLFHFNVAIQVVHNDRD---IEFLQFSKWLNSLYE-----------G---TLEL
YP_009609725.1  LGSPHRHLFKFRVTIQVFHDDRE---IEFHQFLNYVEGMYH----------TG---TLSL
QCG76199.1      LGYPHFHYFYFTVKIEVFHNERD---IEFIQFRRFLERLYS----------ES---LLKL
CAB4130199.1    LASRHRHLFKFKVQMEIFHSDRE---LEFHQVLNYCESLFT----------TQ---SIDI
USV40849.1      LAHPHRHLFHFKVAIEVFHDDRE---IEFHQFLNFLEAAVA---------------LNGL
QRE00068.1      LRDRHRHLFKFRIQIEVHHDNRE---IEFHQFLNMVEGWYE----------DG---TLEL
QIW86898.1      LGDRHFHYFNVRVWIEVKHEERD---IEFIQLRRRIVAEYQ----------KG---GLEM
CAB4149958.1    LRDKHRHIFHVRARMNVSHDDRE---LEFILVQHKLKALVS----------CF---GYDL
CAB4127312.1    LESEHRHIFKIEVTISVNYADRE---LEFFLVKWKLQEFLT---------------TTNQ
AGH13910.1      LRNNHRHMFNIELHIPVMDLNRE---IEFIDEQRIIKDRIL---KEFGDS-LG---YAQF
                    * * : .                ::

NP_417245.1     ENPTSEVLAKWIWDQVKPVV------P-----LLSAVMVKETCTAGCIYRGE--------
NP_389256.1     SLPTTEVVAKTIYDNVQAYLDTLENKP-----TCVQVFVRETPTSYCVYRPKKGGLNG--
YP_007673448.1  EGLSCESIAEELMIQFKATM----------------VKVDEDGENYAKITKQ--------
CAB4159940.1    GSMSCEMIGEWLLNEFPALY---------------KVEVWEDFENGAIIER---------
CAB4155266.1    GSMSCEMIGEWLLEAFSSLY---------------KVEVWEDWENGAIIER---------
CAB4125606.1    GPRSCEMLAKEILEEFDAEW----------------VEVWEDMENGAKVEKV--------
CAB4142608.1    GAMSCEMIAEDLFNYFGLCR----------------CSVSEDGEFFGIIEA---------
QDP63658.1      GTMSCEMIARVLMDQFKLNY----------------CKVLEDNENGAEIKL---------
QDP49444.1      GNMSCEDIAEEIIKLDDSIH---------------WVEVLEDNKGGARIER---------
CAB4128316.1    DWKSCEMIADDLYLQIAGRY------PG----RAITIEVSEDGENGCTITYNLTRPSQSI
CAB5215170.1    DWKSCEMIADDLYLQIAAAY------PG----RNVIIEVSEDGENGCSINYPLTRPNLSV
CAB4137873.1    DYKSCEMIADDIYIKIAERY------PN----RAVWIEVAEDGENGCLIKYELSRPNLSI
CAB4241859.1    DYKSCEMIADDLYKQISATY------TD----RKVWIEVSEDNENGCIKQY---------
CAB4125953.1    DYKSCEMMSDDLYDVISKKY------PG----REVWIEVSEDGENGSLIKY---------
```

```
CAB4221099.1    DHKSCEMISDELYQVIATRY------PD----RDIEIEVSEDGENGCNISYHRYSAKQIS
CAB4163928.1    DYKSVEMMADDLYQEIAVKY------PY----RHVVIEVSEDGENGCCISYLNN------
CAB4123241.1    DFKSCEMIAEDLYTVINDRY------PG----RCVNIDVSEDGENGAWMMWKP-------
CAB4140455.1    NRKSCEMISDELAVYIKDKY------PG----RDLQITVSEDGENGATCRYLSS------
CAB5220926.1    DFQSCEMISDALAERIKDKY------PG----RDIKISISEDDENGSVCTYLAD------
QIG70625.1      DYKSCEMLSEELINTISERY------PG----RNVTVTVSEDDENGATLIYQA-------
YP_009609725.1  DHRSCEMIADELAFTLRKRY------PN----RWMQIEVSEDGECGAVIDYEPATAWDGE
QCG76199.1      DGKSCETMAEELYEQINIVY------PN----RDVVITVAEDNINKAVLEFFK-------
CAB4130199.1    DSKSVEMLADDLYTKLAEKY------PG----RGIKINVSEDGECGCWIEYESPNDNILF
USV40849.1      DYKSCEMIADDIAKIVAEKY------PAL--YRTLEVEVSEDGECGVHAIYGISNGEVFG
QRE00068.1      NHKSCEMIADDLAKRLDQHY------QGDRVKRDLIIEVSEDGECGAVCHYIIQ------
QIW86898.1      NGKSCETLSDELFKTLNEWY------PN----RDIKIDISEEGINGAYVEYSVA------
CAB4149958.1    GRLSCEQIAYEILKYCVQDL------GR----SEIEVEVSEDGENGAVVRYG--------
CAB4127312.1    NHKSCEMIAFDILENHLFDLY----GDD----RTYEVVVSEDGESDGIVRYHPDLLHNGD
AGH13910.1      EGMSCEHIAEWLMAEYPTAI---------------YCKVIEDTNGGAVLVRKQH------
                   : * :.  :                          : *

NP_417245.1     -------------
NP_389256.1     -------------
YP_007673448.1  -------------
CAB4159940.1    -------------
CAB4155266.1    -------------
CAB4125606.1    -------------
CAB4142608.1    -------------
QDP63658.1      -------------
QDP49444.1      -------------
CAB4128316.1    VI-----------
CAB5215170.1    VI-----------
CAB4137873.1    KI-----------
CAB4241859.1    -------------
CAB4125953.1    -------------
CAB4221099.1    V------------
CAB4163928.1    -----------LT
CAB4123241.1    -------------
CAB4140455.1    -------------
CAB5220926.1    -------------
QIG70625.1      -------------
YP_009609725.1  EGV----------
QCG76199.1      -------------
CAB4130199.1    TGTRVQTT---RR
USV40849.1      -------------
QRE00068.1      -------------
QIW86898.1      -------------
CAB4149958.1    -------------
CAB4127312.1    L------------
AGH13910.1      -----QNPFCWFR
```

**Alignment S4: DpdN**

```
YP_010114479.1  MS----------------------------------------------------------
CAB5221950.1    ------------------------------------------------------------
CAB4142580.1    M-----------------------------------------------------------
CAB5226463.1    MQVD--------------------------------------------------------
CAH1079710.1    M-----------------------------------------------------------
BCU93200.1      MT-------TMHVQVDAKTGL------WLEPQSVEVP------------KYL--------
AGN12222.1      ------------------------------------------------------------
```

```
YP_214283.1      ------------------------------------------------------------
YP_004324978.1   ------------------------------------------------------------
YP_007674480.1   ------------------------------------------------------------
AYV82304.1       ------------------------------------------------------------
CAH1078398.1     MNMTTANTARLLITCEDKPGIVQAVSSFLYHQGANITALDQYATEAQGGRYFMRVEFELD
CAH1082857.1     ------------------------------------------------------------
UJE15658.1       M-----------------------------------------------------------
QDF17535.1       M-----------------------------------------------------------
URM86103.1       M-----------------------------------------------------------
QNL30130.1       M-----------------------------------------------------------
QGH80014.1       M-----------------------------------------------------------
CAB4130405.1     MR----------------------------------------------------------
YP_001111230.1   ------------------------------------------------------------
AYV83854.1       ME----------------------------------------------------------
AYV80424.1       M-----------------------------------------------------------

YP_010114479.1   ---------------------------------RPWVAFFSQT---GSEIVEVSKL--LG
CAB5221950.1     ---------------------------------MKWVAFFSQT---GSEIVELSKK--LG
CAB4142580.1     -----------------------------------WVALFSQT---GSEIYNIAQR--LR
CAB5226463.1     ---------------------------------ENWVAFFSQS---GTELNNIIYH--TG
CAH1079710.1     ---------------------------------MKIAVLVSGN---GSNLQALIDA----
BCU93200.1       -------------------------------KKLRIGVMCSGN---GTNFENIVRS----
AGN12222.1       ---------------------------------MRLGIMCSGN---GTNFENIVTN--PL
YP_214283.1      ---------------------------------MRLGIMCSGN---GTNFENIVTN--PL
YP_004324978.1   ---------------------------------MKLVVLCSGN---GTNFENIVTN--PL
YP_007674480.1   ---------------------------------MRIGVMCSGN---GTNFENIVEN----
AYV82304.1       -----------------------------------------------TNLQAIIDACHNK
CAH1078398.1     HLQSRKDALIQTFAANVAERYGMQWRLAFVNDIKKVGILVSKV---DHALLELLWRHARG
CAH1082857.1     -----------------------------------MKIIFAGTPEFAATALAALLK----
UJE15658.1       -----------------------------------STVFLAGSGWFGAEAARALHA----
QDF17535.1       -----------------------------------STVYLAGSGWFGAEAARRLAD----
URM86103.1       -----------------------------------STVFLAGSGWFGAEAAGRLAD----
QNL30130.1       -----------------------------------STVFLAGSGWFGAEAARRLAD----
QGH80014.1       -----------------------------------TTIYLSGSGWFGAEAAKALTD----
CAB4130405.1     -------------------------------EPHSLRIVVAGQKHFGQKVAETLHA----
YP_001111230.1   -----------------------------------MRLMIVGQKWLGAELLKQCVR----
AYV83854.1       ---------------------------------KPWKILVLSA-----------------
AYV80424.1       -----------------------------------WRILVLSA-----------------

YP_010114479.1   RWPDMIVTNERPEHLRKIHPA--------LESK------HLVFVENKPT-------DEEL
CAB5221950.1     CKPALIVTNNFEEKI-KFHPG--------IREL------DTTILSAK-H-------DAIM
CAB4142580.1     THPKYILTNNADRE--TWHPG--------LEEL------DSTIITGK-H-------AMLM
CAB5226463.1     KIPAAIITNRQSDD--DVNPV--------LKQVKDKGLINWITLPKTPE-------SKDY
CAH1079710.1     NLSGQIVGVLSNKA--DAYA---------LERAQNANI-ATAVISHK-DFPSREDFDEAM
BCU93200.1       CRQDEVVLMIYNKK--HCGA---------KKRATKLGI-PNCHVKSS-H-------EDDI
AGN12222.1       CSKHEVVLMIHNTK--KCGA---------VARAAKYGI-PHIRIPHK-D-------EDKM
YP_214283.1      CSKHEVVLMIHNTK--KCGA---------VARAAKYGI-PHIRIPHK-D-------EDKM
YP_004324978.1   SNKHEVVLMIHNKE--KCNA---------VKRAAKFGI-PHIHIPHK-N-------EDLM
YP_007674480.1   CPDHEVVVMIYNIK--GCGA---------QERAERLGI-PNCRIKSI-D-------EQKI
AYV82304.1       VIHADIVLLISNRP--DVGA---------LTKAQNHNI-PTYCSVYKPNMIARTEYDKQL
CAH1078398.1     SLPCEITHVISNHE--DL-----------REAVENFGI-PFTVIKVT-K-------DNKA
CAH1082857.1     -TSHEIIAVYTQPDR-KAGRGQKLTPSPVKQLALEHNI-P--VYQPL-HFKASTEEGLAA
UJE15658.1       -DGHKIVGVSSPRA--DRRNAER--PD-LLALWAELGHVPWVGREAL-R-------PDDV
QDF17535.1       -AGHKIVGVSSPRA--DRRNEDR--PD-LLALWAELGHVPWVGRDAL-R-------PEDV
URM86103.1       -AGHKIVGVSSPRA--DRRNEDR--PD-LLALWAELGHVPWVGRESL-R-------PDDV
```

```
QNL30130.1        -AGHKIVGVSSPRD--DRRNADR--PD-LLALWAELGHVPWVGRDSL-R-------PDDV
QGH80014.1        -AGHKIVGVSSPRE--DRRNAER--PD-LLALWAELGHYPWVGRDSL-R-------PDDV
CAB4130405.1      LAGIEIAAVAAPAD--DRLAD--------FAESRRIHV-----ITAL-R-------AHNM
YP_001111230.1    -DGHEIAAVAAPAA--TGEEYDR-----LYATAQQLAVPASIVGRWL-D-------PDSI
AYV83854.1        -YPQLLLPGLITKEC------------------------DEIIFQNESN-------PVLR
AYV80424.1        -YPELLVPGLISNNC------------------------DEIIFQNERH-------PALT
                       :

YP_010114479.1    SMILGQY----GNPLVTLHGWLRIMPPYICNRF--EIYNGHPGLITEYPELKGKDPQQKA
CAB5221950.1      NYFRNQRIFDVSETFISLHGYLRILPADICEKY--EIYNGHPGAIDLYPELKGKDPQEKV
CAB4142580.1      DLLIGMR----NNPLVTLHGYLRILPEEVVTKH--EVINGHPGDIVKYPELKGKDPQQKA
CAB5226463.1      KKALKPF----KDPLITLHGYLRIIPKDICKKYK-QIYNLHPGLITEYPDLKGKDPQIRA
CAH1079710.1      HQQLIAW----QADVVILAGFMRILTADFVNKWQGKMLNIHPSLL---PAYKGINTHQRV
BCU93200.1        IAFFKAY----NVDLIVLAGWMRIIKNP--ESFPAPMINVHPSLL---PKYKGLHAIEQA
AGN12222.1        IELFKTW----RVDLIILAGYMRVIKNP--SDFPCPIINIHPSLL---PKYKGLNVVQRA
YP_214283.1       IELFKTW----RVDLIILAGYMRVIKNP--SDFPCPIINIHPSLL---PKYKGLNVVQRA
YP_004324978.1    IRTIRAF----APDLIVLAGYMRILSPRFVGSFE-NIINVHPSLL---PKFKGAHAIEQA
YP_007674480.1    IDKLNRH----KVDLVVLAGWMRIVTPGLINAFPNKIINIHPSLL---PKYKGLNAVKQA
AYV82304.1        AEYINTV----DYDLIVCAGWMHLLSKHFLNLISKPIINLHPALP---GQFPGKNSIVDA
CAH1078398.1      EAYAQIHEMMQGNDLLVLARYMQILSEDFVSKWEMKIINIHHSFL---PAFVGANPYKQA
CAH1082857.1      QQELAAL----GADVMVVAAYGLILPQAVLDTPKYGCLNIHGSLL---PRWRGAAPIQRA
UJE15658.1        PA---------NTDVILAAHSHAFIGRRTRNRAQ-YAMGYHPSLL---PLHRGRDAVKWQ
QDF17535.1        PA---------NTDVILAAHSHAFIGRRTRNRAA-YALGYHPSLL---PLHRGRDAVKWQ
URM86103.1        PA---------NTDVILAAHSHAFIGRRTRNRAA-YALGYHPSLL---PLHRGRDAVKWQ
QNL30130.1        PA---------NTDVILAAHSHAFIGRRTRNRAQ-FALGYHPSLL---PLHRGRDAVRWQ
QGH80014.1        PA---------NTDVILTAHSHAFIGRRTRNRAT-YALGFHPSLL---PLHRGRDAVKWQ
CAB4130405.1      PD---------EIDLIIAAHSHDFIGERTRLRARFGGVGYHPSLL---PLHRGRDAVRWA
YP_001111230.1    PD---------EVDLILAAHAHAFLPRAARERARLGALGYHPSLL---PRHRGRDAIRWA
AYV83854.1        PEWILER----GINFIVVFVHNRIIKKNIIDLV--PVINIHGSFL---PLNRGPNPILWA
AYV80424.1        PEWILER----KIDFIIVFVHNRIIKKNILDLV--PAINVHGSLL---PLNRGPNPILWA
                                .:       .:             . * .         *

YP_010114479.1    FDLGL--ESS--GCVIHKVTEGVDEGEILR-SRKVSIKGLEIGELFHIL----HSISVSL
CAB5221950.1      WQEND--KYTIIGSVVHKCTAELDAGDILK-AVHLRNRNYSREELYASL----KMTSLSA
CAB4142580.1      IDLGL--PST--GVVLHKAVAEVDAGEIIKHATRDIFAGTTTEQLILDL----KAIQLDL
CAB5226463.1      VKAGY--NTA--GVVIHKVIPAVDEGEIVS-SHAINIKSLTEEEVITQL----HSLGSIM
CAH1079710.1      LNTGD--RLH--GCTVHFVTSELDAGQAIAQSAIEVKEHDNVASLAERV----HKLEHFI
BCU93200.1        MKAGE--EYT--GCTVHYVNEELDGGEIIIQKEVPILAADDVESLTKAV----QRMEYSI
AGN12222.1        MEAGE--LVT--GCTVHYVNEELDGGEIIMQGEVPILPNDDVESLTKAI----QRKEYAI
YP_214283.1       MEAGE--LVT--GCTVHYVNEELDGGEIIMQGEVPILPNDDVDSLTKAI----QRKEYAI
YP_004324978.1    LESGD--TET--GVTVHYVTEELDSGEVILQTKVPILPNDDVKSLTKAI----QRVEYGI
YP_007674480.1    LDSGD--KIT--GCTVHYVTEELDSGGCIDSSSVPICVGDTEETLHHRV----QRAEHRL
AYV82304.1        YNGSSYCRGD--GCRTSY------------------------------------------
CAH1078398.1      YEKGV--KLI--GATAHYVTADLDQGPIIEQDVERVSHDYNVEQLRELG----EDVERNV
CAH1082857.1      IATGD--DET--GITIMQMAAGLDTGDMMYKTYCPITSEDTSATLHDKL----AAQGATA
UJE15658.1        ARLRE--RVV--GGSIYHLTDRVDGGPLHSQVHTVVPDGIKASDLWRDYL---APIGLDM
QDF17535.1        ARLRE--RVV--GGSIYHLTDRVDGGPLHSQVHAVVADGITASELWREVL---APIGLEM
URM86103.1        ARLRE--RVV--GGSIYHLTDRVDGGPLHSQVHAVVADGITASELWREVL---APIGLEM
QNL30130.1        ARLRE--RVV--GGSIYHLTDRVDGGPLHDQAHAVVPDGIKASDLWREYL---APIGLDM
QGH80014.1        ARLRE--RVV--GGSIYHLTDRVDGGPLHSQVHAVVPDGISASDLWRDYL---APIGLTM
CAB4130405.1      IRMRE--RVT--GGTVYRLSNRMDGGEILAQRHVFIRPDDDAQSLWRRDL---APLGIEM
YP_001111230.1    MHMRE--AVT--GGTVYWMDDGADSGPIALQDWCHIRPDDTPTSLWRREL---GPMGLRL
AYV83854.1        WLNG---SAQ--GVTIHYIDEGVDTGDIIAQKEVVLPTNITYNKSFDIILNQCSRLFRET
AYV80424.1        WLNK---TQQ--GVTVHYMNEGVDTGDIIAQKSVVLPTDITYSKSFDIIVKECANLFRMT
                              *

YP_010114479.1    WVDFLKN-----------------------------------------------------
```

```
CAB5221950.1     WSFFLREKGL--------------------------------------------------
CAB4142580.1     WCDLLRERL---------------------------------------------------
CAB5226463.1     WYQFFKDYEHRRNSKGN-------------------------------------------
CAH1079710.1     YPQVAEWLCNGQLT-----WKNGQAYFRNQ------------------------------
BCU93200.1       LPEAIKNVKYKIQTRVN-------------------------------------------
AGN12222.1       LPAAIDSLG---------------------------------------------------
YP_214283.1      LPAAIDSLG---------------------------------------------------
YP_004324978.1   LPQAINLCASSETV-----PRVAHSDLRLL------------------------------
YP_007674480.1   LPMVINNLFENIN-----------------------------------------------
AYV82304.1       ------------------------------------------------------------
CAH1078398.1     LARAVKWHLEDRIIV------DGNKTVVF-------------------------------
CAH1082857.1     ICAVLESEETLQKYLAEREVQDESLTVYAHKLVKSEARIDWSMNAVQVDRNIRAFNPWPV
UJE15658.1       LRRAAQAVE------------DGEVNYR--------------------------------
QDF17535.1       LVRAAQAVE------------DGEVNYH--------------------------------
URM86103.1       LVRAAQAVE------------DGEVNYH--------------------------------
QNL30130.1       LVRAAQAVE------------DGEVNYR--------------------------------
QGH80014.1       LVRAAQAVE------------DNEVNYR--------------------------------
CAB4130405.1     LASVVLRFA--------------EFGYQT-------------------------------
YP_001111230.1   FARALAMIE------------QGACPSS--------------------------------
AYV83854.1       WPLIRGGVNERR-------LQSGESSCHR--LADQAVLNDF-------------------
AYV80424.1       WPLIRAGMNERR-------VQSGEITSHR--LADQKVLNDF-------------------

YP_010114479.1   ------------------------------------------------------------
CAB5221950.1     ------------------------------------------------------------
CAB4142580.1     ------------------------------------------------------------
CAB5226463.1     ------------------------------------------------------------
CAH1079710.1     ---ILEHPIRFASW----------------------------------------------
BCU93200.1       ------------------------------------------------------------
AGN12222.1       ------------------------------------------------------------
YP_214283.1      ------------------------------------------------------------
YP_004324978.1   ------------------------------------------------------------
YP_007674480.1   ------------------------------------------------------------
AYV82304.1       ------------------------------------------------------------
CAH1078398.1     ------------------------------------------------------------
CAH1082857.1     AFIQLDENNALRVWNSTISNQNKADAQAGEIIAIDKQGVHVACGENTFICLTSVQWPGGK
UJE15658.1       -----PQDDSMATWEPSF---DSAPLYRPELL----------------------ELPRGA
QDF17535.1       -----PQDDALATWEPSF---DSAPLYRPELM----------------------ELPRGA
URM86103.1       -----PQDDALATWEPSF---DSAPLYRPELM----------------------ELPRGA
QNL30130.1       -----PQDDALATWEPSF---DSAPLYRPELV----------------------ELPRGS
QGH80014.1       -----PQDDALATWEPSF---DSAPLYRPELM----------------------ELPRGR
CAB4130405.1     ---GAPQDEELATWEPSI---DRPPAFRPDLM----------------------LLTDQR
YP_001111230.1   -----EQDSALATWEPAF---RSKSLSAAT------------------------------
AYV83854.1       ---IESNQNSMGILE---------------------------------FCEKARGILGGS
AYV80424.1       ---IRHNQNSMGILE---------------------------------FCEKARLILEGK

YP_010114479.1   --------------------
CAB5221950.1     --------------------
CAB4142580.1     --------------------
CAB5226463.1     --------------------
CAH1079710.1     --------------------
BCU93200.1       --------------------
AGN12222.1       --------------------
YP_214283.1      --------------------
YP_004324978.1   --------------------
YP_007674480.1   --------------------
```

```
AYV82304.1       --------------------
CAH1078398.1     --------------------
CAH1082857.1     ALNTQQIAQTQKLHVGQILP
UJE15658.1       S-------------------
QDF17535.1       AR------------------
URM86103.1       AR------------------
QNL30130.1       T-------------------
QGH80014.1       G-------------------
CAB4130405.1     ATGTKK--------------
YP_001111230.1   --------------------
AYV83854.1       AKKEKKRRKRKRK-------
AYV80424.1       DPNDVKMKLAKKKKKKN---
```

**Alignment S5: ArcS**

```
nt-1             --------------------MNS--VQDKSVL-------------------MGDELL---
STSV1            --------------------MSY-------------------------------------
STSV2            --------------------MSY-------------------------------------
S.solfataricus   --------------------MS--------------------------------------
I.hospitalis     --------------------MES-------------------------------------
A.fulgidus       M---FYPEKRDGFSRTGKFEVEGRTVQTPAILEVGEIPEWDFGLAPTSLKFISEELYS--
H.volcanii       MTDYFEVHARDGAARIGELRLSD-SVTTPAVVDDVLADAGSLWAAERELPDGSDDVLTVL
M.thermautotrop  M-----------------IKVDR-------------------------------------
M.smithii        ------------------------------------------------------------

nt-1             -----------------------------------------------------------F
STSV1            ------------------------------------------------------------
STSV2            ------------------------------------------------------------
S.solfataricus   ------------------------------------------------------------
I.hospitalis     ------------------------------------------------------------
A.fulgidus       --RLRPINEEVEILTSLHLLSPR-------------------------------QLVQVF
H.volcanii       PHRSLPAGSADEVRESFSVAYPDVDFPSAAVVTADTADDFGADAYVLSDAQGFVGHARAF
M.thermautotrop  ------------------------------------------------------------
M.smithii        ------------------------------------------------------------

nt-1             SE----------------------------------------------------------
STSV1            ------------------------------------------------------------
STSV2            ------------------------------------------------------------
S.solfataricus   ------------------------------------------------------------
I.hospitalis     ------------------------------------------------------------
A.fulgidus       EDLSKEGVSPKP----LYAASSALPSNVSLLIYLGADLVDNVLAIAKAYSGIYFLGEVEV
H.volcanii       RDNVIEAKENLPADTALVLSGVATPRNVSLLVYAGVDLVDEKLARARGLEGFYLTSDGEY
M.thermautotrop  ------------------------------------------------------------
M.smithii        ------------------------------------------------------------

nt-1             ------------------------------------------------------------
STSV1            -----------VESC---------------EHLEKYLGS---------NNEPIIKKP---
STSV2            -----------VESC---------------EHLEKYLGS---------NNEPIIKKP---
S.solfataricus   ------------SAC---------------EQLYEYLKP---------DFTRISKRD---
I.hospitalis     -----------LVACLNKKACRAELWRRAKETASKYKDELVNIAESLMNDERINERELDL
A.fulgidus       EISKLRRLPCNCIHCRNRVVDEVENLL---ETTAKHNTEMLRM-----EVEKCRRLILNE
H.volcanii       FLEDLDELPCACEACRKPA---SEFTR---ADAADHNANALRA-----ELARVRRRVRDG
M.thermautotrop  ------------------------------------------------------------
```

```
M.smithii       ------------------------------------------------------------

nt-1            ------------------------------------------SEWVP----FDTMNTEKK
STSV1           --------------------------------------------------------GFDP
STSV2           --------------------------------------------------------GFDP
S.solfataricus  --------------------------------------------------------NTDD
I.hospitalis    ALKALV-----VDPQYLHFKGIKVVTPIGFYLAREYDLDT--VYATP----GEVVLEGDW
A.fulgidus      ELRNYVEGKVKLNPEFTA------------ALRLSDSLRN--HSTFPRFRKSRCNFSALE
H.volcanii      RLRDYVEGQARHDQWLTA------------LFRRFDQQYSFMEQRVPVIRDSELTAASEE
M.thermautotrop ---------------------------------------------------MKVICSIEE
M.smithii       ---------------------------------------------------MKVICSSEE

nt-1            ILSNPIFDEGLNRIIKQYTPK---TDTAFISLCTTTRPYENSRKWKTFMETFGSSA----
STSV1           -FQHSVVRKWHEYFLNNWYSKK-EKEIALLLPCTSVKPYNRSATHKLAYSLLRKYKLEDK
STSV2           -FQHPVVRKWHEYFLNNWHSK---KEIALLLPCTSVKPYNRSATHKLAYSLLRKYKLEDK
S.solfataricus  LFRHPVVMRWHRYFLENWSSN---KEIALLLPCTVVKPYSSSPTHKIAYAILQKYNLEEK
I.hospitalis    LFLHPTVKDYHEFLFSYLLDKMGPGEVALITPCSKVKPYRDSFMYKKIESIINKYG--ND
A.fulgidus      SSSRFEVRYFFERALECYKPF---SDTVLLLPCTARKPYLTSRTHRALRSKVKVNV----
H.volcanii      SIDRVEIQRFADRVTKRYRNRF--DNPLVLLPCSAKKPYSESQSHRQFQEAVQYRA----
M.thermautotrop SLHRPEAVRWRER-MRLMRPL---GDVVVVLPCSMKKPYSTSRSHRKFTGVTRGFQ----
M.smithii       SLYRPEAVRWRQR-MELMKPL---GDTVVLLPCSMKKPYSNSKSHQKFRKVSRSFQ----
                   .                     :  .:  *:  :**  *   :

nt-1            --DLIVISNGGVIPQQFWHSYPYLNYDGDPK-VNPQTELYCEICEERVYRFFS--QVKFK
STSV1           VQVYSVSEPMLLVPRELENCYPFNSYDYPPS-LMTEEEK--QEFIRLLSIALSHIHKIHK
STSV2           VQVYSVSEPMLLVPRELENCYPFNSYDYPPS-LMTEEEK--QEFISLLSIALSHVHNIHK
S.solfataricus  VQVYSVSEPMLLVPRELEECYPFNNYDYPPA-LMSKEEK--EEFVELLTKPLIKISKLHE
I.hospitalis    VWRFVMSEPLVIVPRFFDVYFPSAHYDYPPE-RVAPEEV--PIYVDLVRKALEVIATRFE
A.fulgidus      -NEIIISSPL-VVPREFELLYPAVNYDTPVTGHWSEEEV--SFVAGWLKRFIE--KGGFR
H.volcanii      -HMVSMTSPIGVVPQELELTYPAQHYDSVVTGDWSEDEK--SFVAEVLRRYLE--RNDYP
M.thermautotrop --ELILTSPFGICPRELENTYPISSYDVSTIGEWSHEEK--KLVGDVLREYVG-----DL
M.smithii       --ELIITSPFGICPRELENTFPIQSYDVAVTGSWSQDEI--DESGKLLEKYVK-----GK
                     : .   : *: :   :*   **       .  *         :   .

nt-1            NIVANFKHVQRNTEAIKSALKRLKDDGHIENYVVLP----TVEQYEELQSRGFPKGKVFP
STSV1           RIVAVLP------KHHYKIVYEASK--NIPIE-LYPYGKLAFKTIANVIMKI--------
STSV2           RIIAVLP------KHHYKIVYEASK--NIPIE-LYPYGKLAFKTIANVIMKI--------
S.solfataricus  RLIGILP------RHHYEIVKGAAEISHLKIT-IYPYGRLAFKTIANVIRNLS-------
I.hospitalis    RIVYTLP------RKHKRIFEMAAEQANLSAN-YTPYNVYFFPRLKEAIV----------
A.fulgidus      KVVAHVT------GGYRKVVERVEDEVEAEVVYTAEKDVLSDESIERLKQEIESKGKV--
H.volcanii      RIIAHLPP-----GAYTDIVERVADDLDLDVEFTVSEHPTTTESIGNLMRTLDGEPKF--
M.thermautotrop DVIAHVD------GGYLEVCMEYLDSFTAT---SMEARPTSPEALERLKRELESYERI--
M.smithii       NVIANVH------GGYEEVCRQYLDECTYT---CKDGRPTSPDSIYNLRMELKEHKRN--
                 ::  .                  .                     .

nt-1            DVDDKILEHLRS-AVHSYSPPKENSLA---------------------------------
STSV1           ------------------------------------------------------------
STSV2           ------------------------------------------------------------
S.solfataricus  ------------------------------------------------------------
I.hospitalis    ------------------------------------------------------------
A.fulgidus      DLYRRILEHMLS---YQFGITWSG--------KVAGRYPELELL-EGKKRLARVDRIYGM
H.volcanii      TREEREHNVVKALADYQLGPDAGDALFSDVALEMTSRYPKLQVWNDAGVQLATMVPQYGV
M.thermautotrop PPRERLLHMLRSVAVYQFGPGGERIIPEDC--RVTGRYHR-KILKSDGTQIGTFMMDRGL
M.smithii       NRRDKVLNELKSIATYQFGKEGSKLIPEDV--KTKGRYHR-KII-SNGKQLALLNQDTGL
```

```
nt-1              -----------------------NFF----------------------------------
STSV1             ------------------------------------------------------------
STSV2             ------------------------------------------------------------
S.solfataricus    ----------------------------------------------------LSNPRSLI
I.hospitalis      ------------------------------------------------------------
A.fulgidus        LDIYEKIAAYLLEKN--IYTVEIGDFEVKGTIFAGGVLRADEKIRPNDV-VVFHNSRIFG
H.volcanii        LSFTLEGAKVWRDSDAPTKTVEIDGFVPHGSVLAPGVVDADEDIRPGDE-VVVEGPKAFA
M.thermautotrop ISLSPEGGRSLYMLG--VKWVEIDFNLESNTLFAPGVSDADTGIIPGDEVVIVQGGEVAG
M.smithii         YRLNLAGGEILKDLN--INVVNIDFNLETNTVFAPGIRKADHNIIPNDEVVVVKDDEVVG

nt-1              -----------------------------
STSV1             -----------------------------
STSV2             -----------------------------
S.solfataricus    V----------------------------
I.hospitalis      -----------------------------
A.fulgidus        VGLAAMSGKEMAGSEKGIAINVKRKFS-F
H.volcanii        IGRAEMGGRELVESTRGIGVEIRHVEE-R
M.thermautotrop VGRAVLSGDEMVRAERGVAVRVRRRTG--
M.smithii         VGKAILTGREMEQCTNGIGVKIKHRVKNN
```

**Supplementary References**

1. Baba,T., Ara,T., Hasegawa,M., Takai,Y., Okumura,Y., Baba,M., Datsenko,K.A., Tomita,M., Wanner,B.L. and Mori,H. (2006) Construction of Escherichia coli K-12 in-frame, single-gene knockout mutants: the Keio collection. *Mol. Syst. Biol.*, **2**, 2006.0008.

2. Moore,S.D. (2011) Assembling New Escherichia coli Strains by Transduction Using Phage P1. In Williams,J.A. (ed), *Strain Engineering: Methods and Protocols*. Humana Press, Totowa, NJ, pp. 155–169.

3. Holmfeldt,K., Solonenko,N., Shah,M., Corrier,K., Riemann,L., VerBerkmoes,N.C. and Sullivan,M.B. (2013) Twelve previously unknown phage genera are ubiquitous in global oceans. *Proc. Natl. Acad. Sci. U. S. A.*, **110**, 12798–12803.

4. Nilsson,E., Li,K., Fridlund,J., Šulčius,S., Bunse,C., Karlsson,C.M.G., Lindh,M., Lundin,D., Pinhassi,J. and Holmfeldt,K. (2019) Genomic and seasonal variations among aquatic phages infecting the Baltic Sea gammaproteobacterium Rheinheimera sp. strain BAL341. *Appl. Environ. Microbiol.*, **85**.

5. Lee,Y. and Weigele,P.R. (2021) Detection of Modified Bases in Bacteriophage. In *DNA Modifications Methods and Protocols*.Vol. 2198, pp. 53–66.

6. Erdmann,S., Chen,B., Huang,X., Deng,L., Liu,C., Shah,S. a., Le Moine Bauer,S., Sobrino,C.L., Wang,H., Wei,Y., *et al.* (2014) A novel single-tailed fusiform Sulfolobus virus STSV2 infecting model Sulfolobus species. *Extremophiles*, **18**, 51–60.

7. Zillig,W., Kletzin,A., Schleper,C., Holz,I., Janekovic,D., Hain,J., Lanzendörfer,M. and Kristjansson,J.K. (1993) Screening for Sulfolobales, their Plasmids and their Viruses in Icelandic Solfataras. *Syst. Appl. Microbiol.*, **16**, 609–628.

8. Skliros,D., Kalatzis,P.G., Katharios,P. and Flemetakis,E. (2016) Comparative functional genomic analysis of two vibrio phages reveals complex metabolic interactions with the host cell. *Front. Microbiol.*, **7**, 1–13.

9. Moineau,S., Pandian,S. and Klaenhammer,T.R. (1994) Evolution of a lytic bacteriophage via DNA acquisition from the Lactococcus lactis chromosome. *Appl. Environ. Microbiol.*, **60**, 1832–1841.

10. Vander Meersche,Y., Cretin,G., de Brevern,A.G., Gelly,J.C. and Galochkina,T. (2021) MEDUSA: Prediction of Protein Flexibility from Sequence. *J. Mol. Biol.*, **433**, 166882.

11. Pettersen,E.F., Goddard,T.D., Huang,C.C., Meng,E.C., Couch,G.S., Croll,T.I., Morris,J.H., Ferrin,T.E. and Thomas Ferrin,C.E. (2021) UCSF ChimeraX: Structure visualization for researchers, educators, and developers. *Protein Sci.*, **30**, 70–82.

12. Phillips,G., Chikwana,V.M., Maxwell,A., El-Yacoubi,B., Swairjo,M.A., Iwata-Reuyl,D. and De Crécy-Lagard,V. (2010) Discovery and characterization of an amidinotransferase involved in the modification of archaeal tRNA. *J. Biol. Chem.*, **285**, 12706–12713.

13. Blattner,F.R., Plunkett,G., Bloch,C.A., Perna,N.T., Burland,V., Riley,M., Collado-Vides,J., Glasner,J.D., Rode,C.K., Mayhew,G.F., *et al.* (1997) The Complete Genome Sequence of Escherichia coli K-12. *Science (80-. ).*, **277**, 1453–1468.

14. Phillips,G., Grochowski,L.L., Bonnett,S., Xu,H., Bailly,M., Blaby-Haas,C., El Yacoubi,B., Iwata-Reuyl,D., White,R.H. and De Crécy-Lagard,V. (2012) Functional promiscuity of the COG0720 family. *ACS Chem. Biol.*, **7**, 197–209.

15. Chakravartty,V. and Cronan,J.E. (2015) A series of medium and high copy number arabinose-inducible Escherichia coli expression vectors compatible with pBR322 and pACYC184. *Plasmid*, 10.1016/j.plasmid.2015.03.001.

**Supplementary Figures**

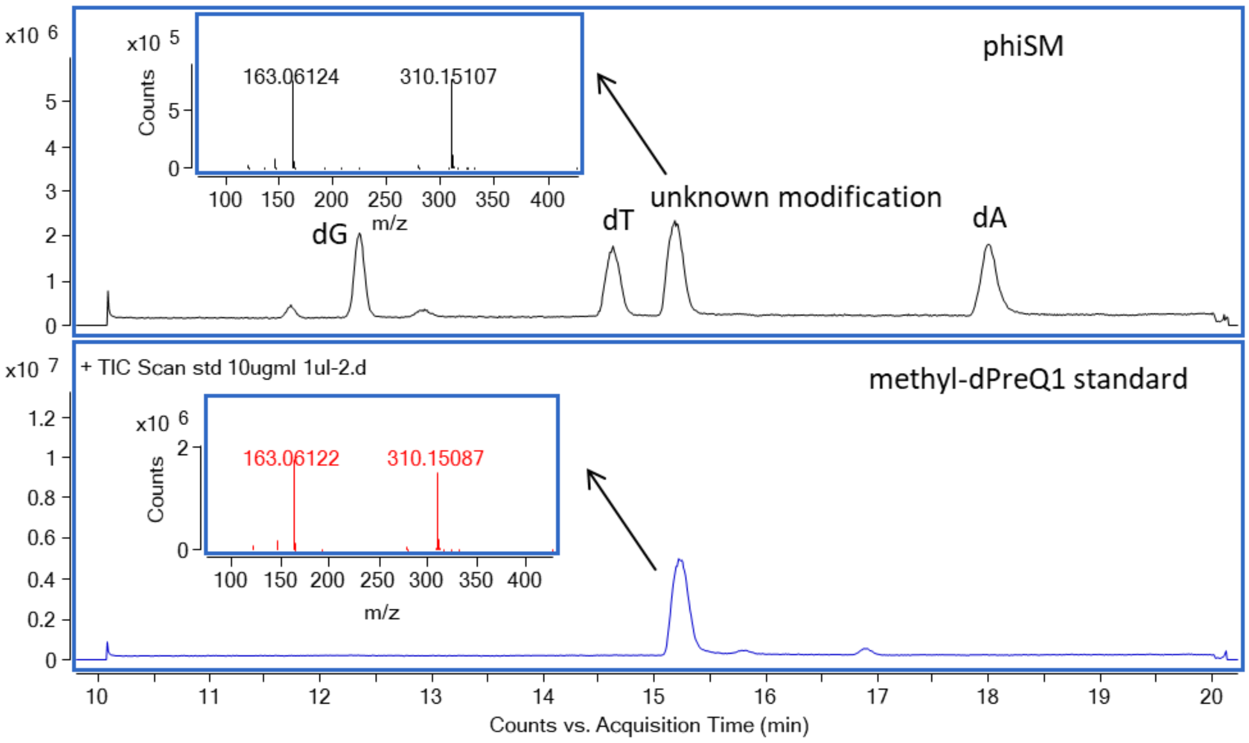

**Fig. S1. LC-MS analysis of digested Cellulophaga phage phiSM DNA sample and methyl-dPreQ1 (mdPreQ1) standard**

The retention time and MS/MS spectra of the unknown non-canonical nucleoside in phiSM DNA (upper) were identical to those of the mdPreQ$_1$ standard (lower).

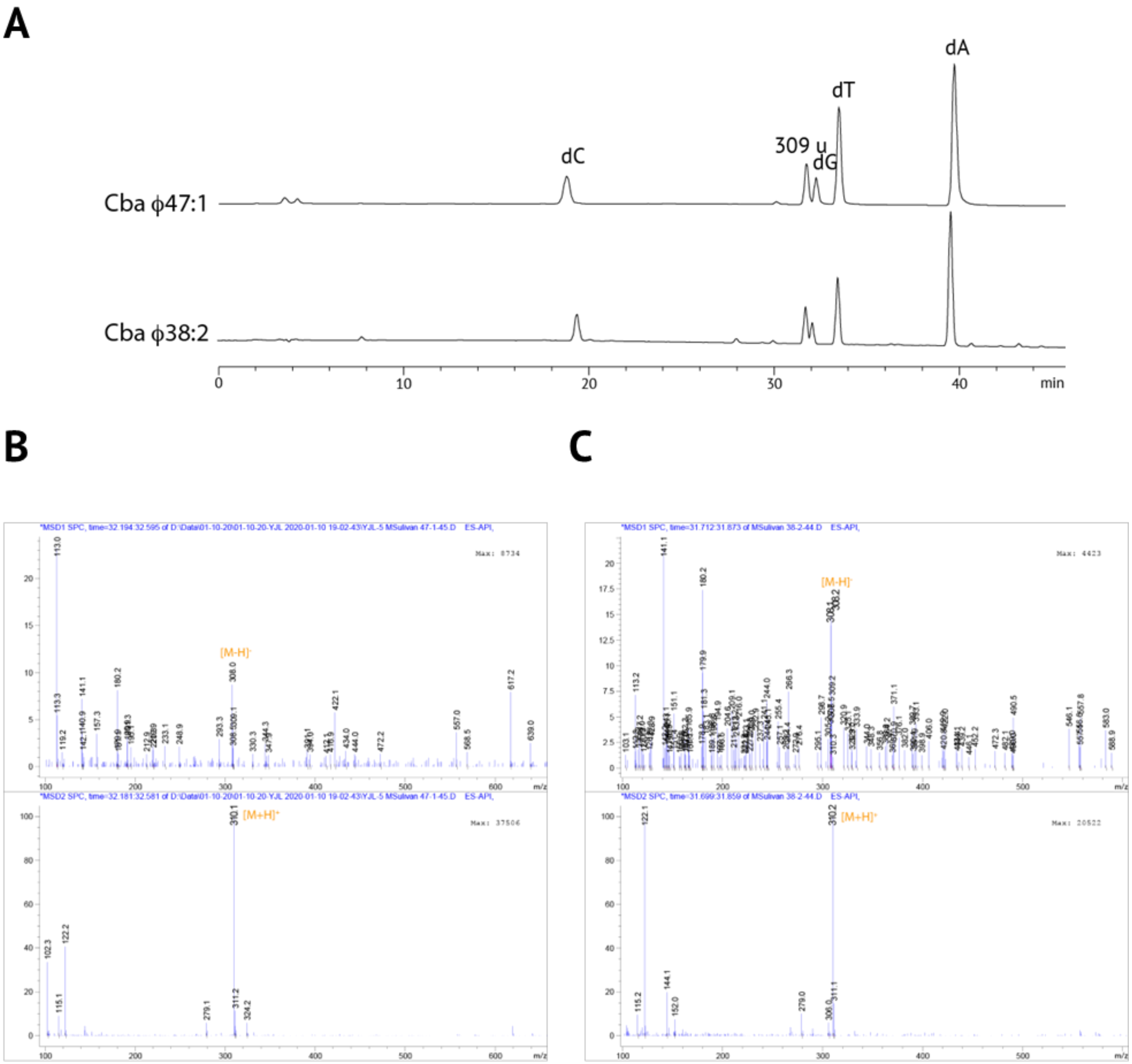

**Fig. S2. LC-MS nucleoside analysis of genomic DNA isolated from Cellulophaga phages phi47:1 and phi38:2.**

(A) HPLC traces of Cellulophaga phages phi47:1 and phi38:2 genomic DNA enzymatically digested to free nucleosides. (B) MS signals from the non-canonical nucleoside species observed at 32.1 min in the LC chromatogram of phi47:1 genomic DNA digest. The deconvoluted mass of the species is 309 g/mol. (C) MS signals from the non-canonical nucleoside species observed at 31.7 min in the LC chromatogram of phi38:2 genomic DNA digest. The deconvoluted mass of the species is 309 g/mol.

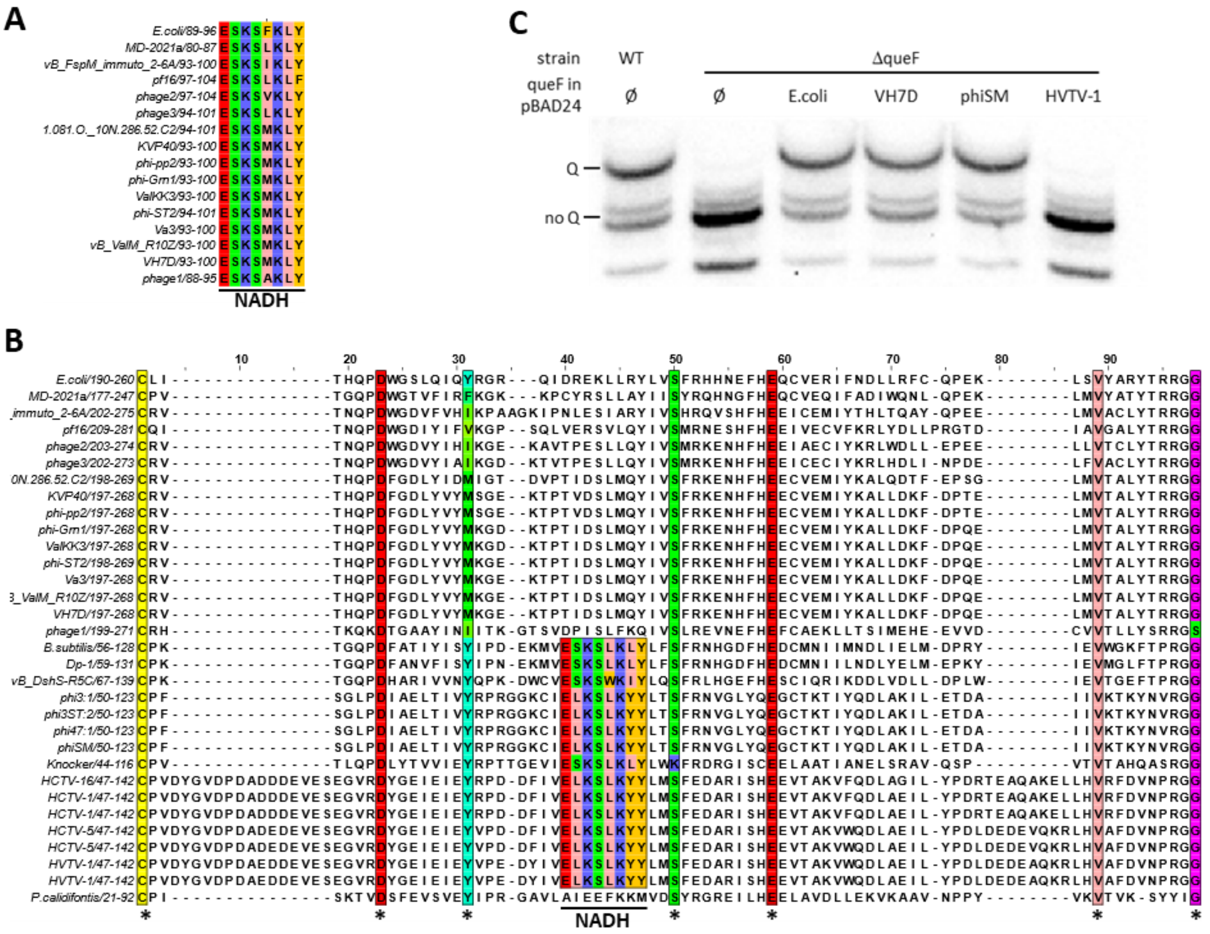

**Fig. S3. Phage QueF are NADPH-dependent 7-cyano-7-deazaguanine reductase.** (A) Alignment of the NADH binding site (E(S/L)K(S/A)hK(L/Y)(Y/F/W)) of the phage bimodular QueF compared to *E. coli* bimodular QueF (NP_417274). (B) Alignment the reactive amino acids (stared residues) of QueF compared to *E. coli* bimodular QueF, *B. cereus* unimodular QueF (NP_389258), and *P.calidifonti* QueF-L (WP_011848915), in addition to the NADH binding site of the unimodular QueF. (C) Northern blot of acrylamide-APB gel electrophoresis of tRNA extracted from different strains of *E. coli* MG1655 (WT and Δ*queF* strains) transformed with pBAD24 (empty vector control) or *queF* derivatives expressing the respective homologous genes from *E. coli*, *Vibrio* phage VH7D, Cellulophaga phage phiSH, and Halovirus HVTV-1. tRNA lacking Queuosine (no Q) migrates faster than tRNA containing Queuosine (Q).

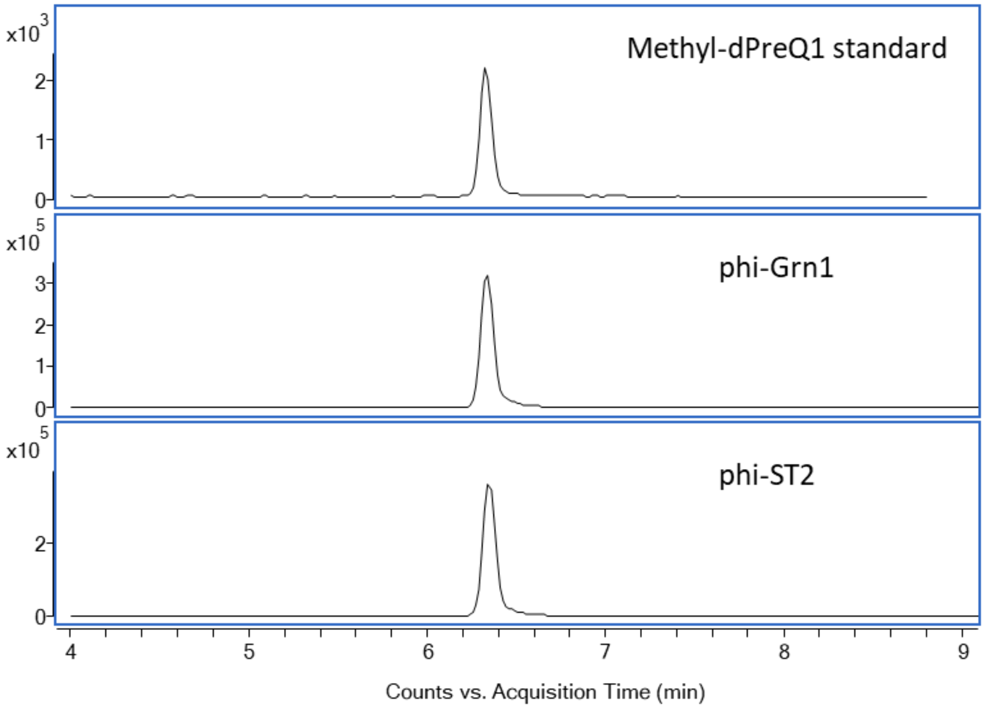

**Fig. S4. LC-MS analysis of digested Vibrio phage phi-Grn1 and phi-ST2 DNA samples and methyl-dPreQ1 standard**

The retention time and multiple reaction monitoring (MRM) transition (m/z 310→163) of the unknown non-canonical nucleoside in phi-Grn1 DNA (middle) and phi-ST2 DNA (lower) were identical to those of the mdPreQ$_1$ standard (upper).

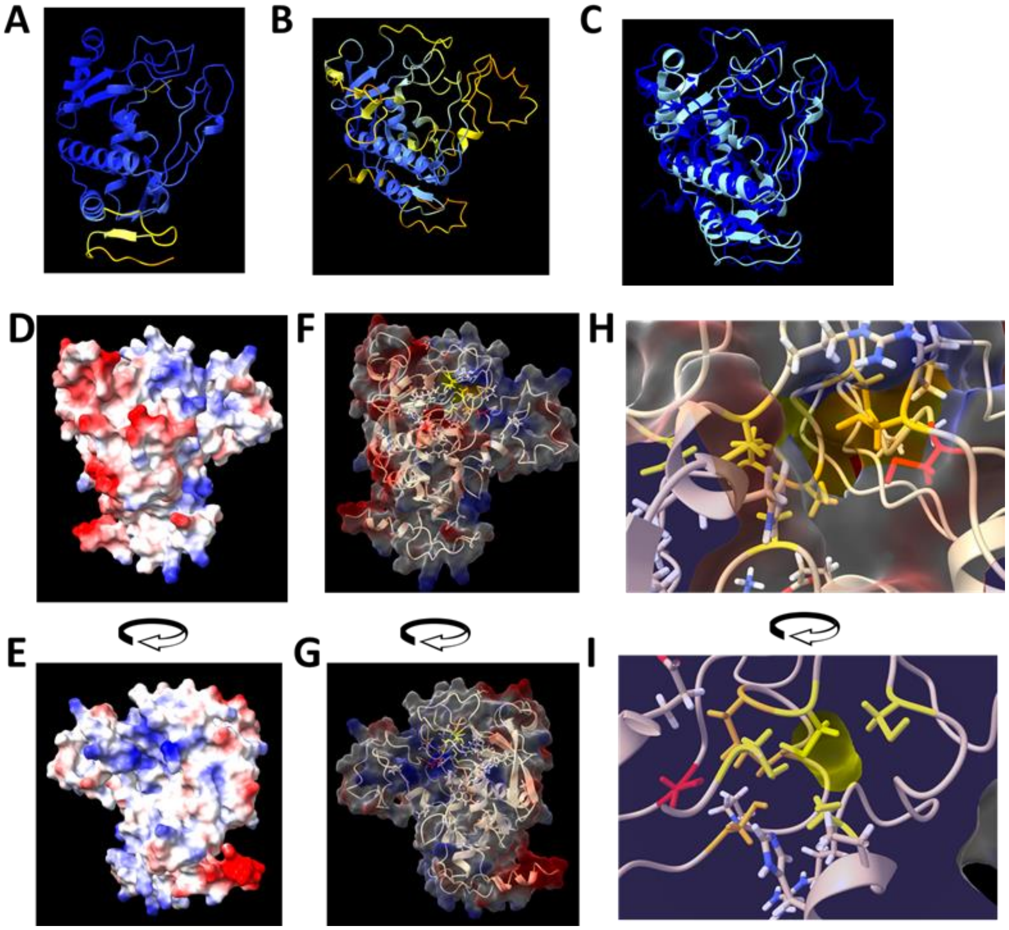

**Fig. S5. DpdM structure prediction**

AlphaFold2 predictions of Cellulophaga phage phiSM DpdM (A, YP_007675730.1) and Vibrio phage VH7D (B, YP_009006155.1), colored by confidence score, with yellow being low confidence and blue high confidence, and overlayed structures (C). Electromagnetic surface charges of VH7D DpdM proteins were visualized on two opposite faces of the proteins matching with phiSM DpdM in figure 3 (D and E,), blue are positive charges and red negative charges. Conserved residues were added to the structures with a 60 % transparency on the electromagnetic surface (F and G). Conserved cysteines predicted to bind metal are colored in yellow, orange, and red as described in the text. Visualization of these cysteines were zoomed in to view both metal pockets (H and I).

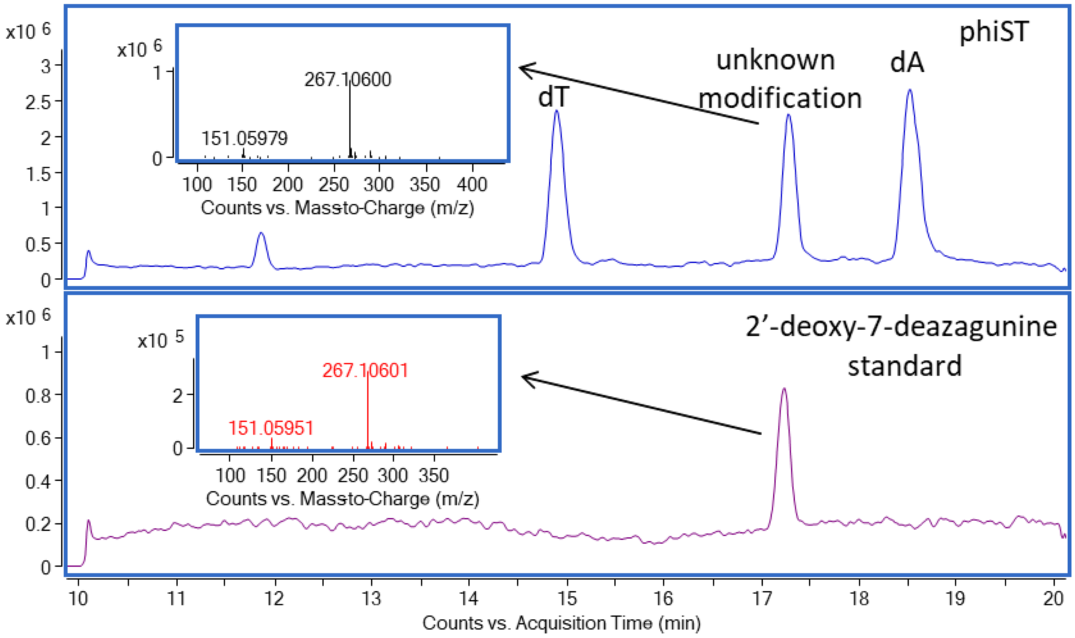

**Fig. S6. LC-MS analysis of digested Cellulophaga phage phiST DNA sample and methyl-2'-deoxy-7-deazagunine (dDG) standard**

The retention time and MS/MS spectra of the unknown non-canonical nucleoside in phiST DNA (upper) were identical to those of the dDG standard (lower).

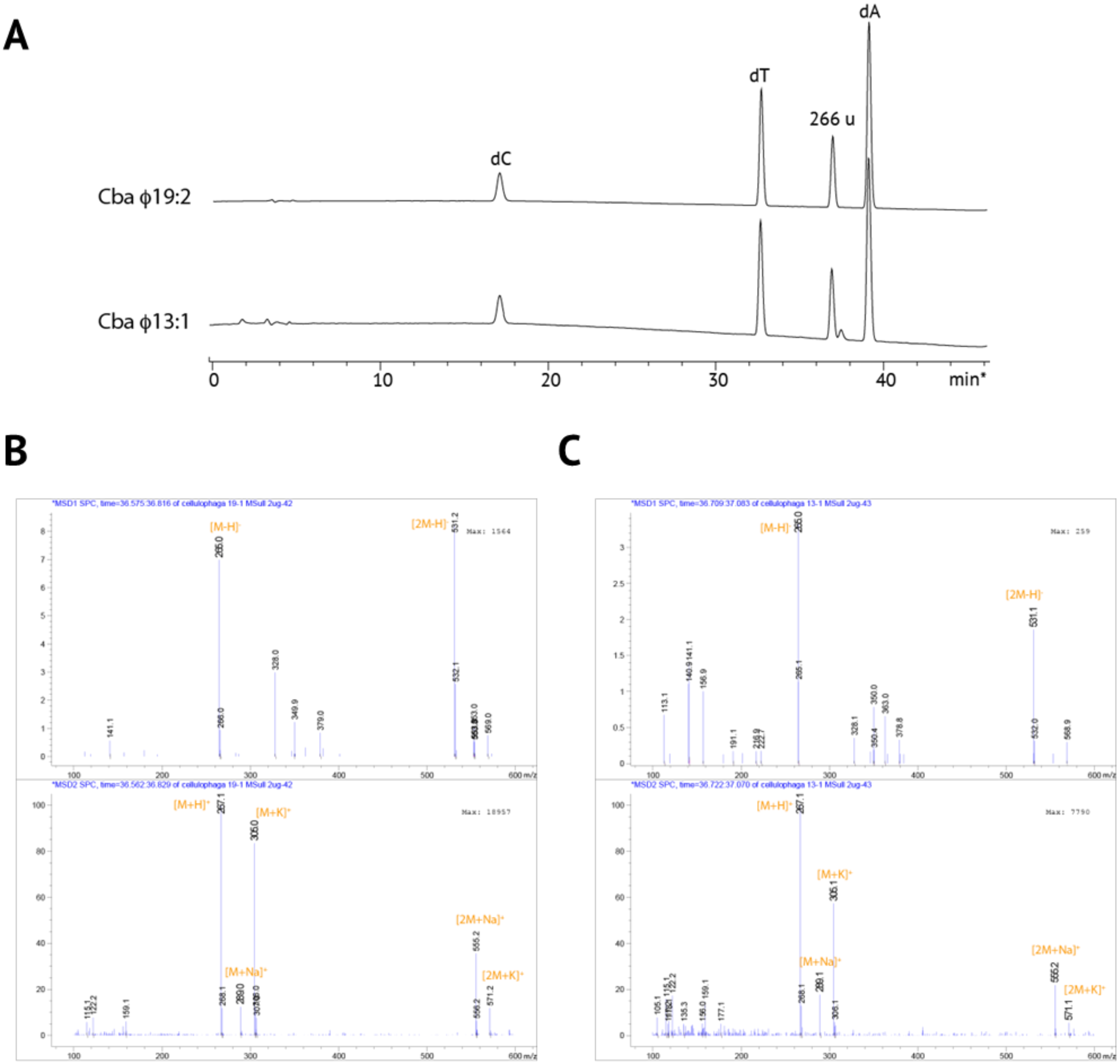

**Fig. S7. LC-MS nucleoside analysis of genomic DNA isolated from Cellulophaga phages phi19:2 and phi13:1.**

(A) HPLC traces of Cellulophaga phages phi19:2 and phi13:1 genomic DNA enzymatically digested to free nucleosides. (B) MS signals of the non-canonical nucleoside species observed at 36.5 min in the LC chromatogram of phi19:2 genomic DNA digest. The deconvoluted mass of the species is 266 g/mol. (C) MS signals of the none-canonical nucleoside species observed at 36.7 min in the LC chromatogram of phi13:1 genomic DNA digest. The deconvoluted mass of the species is 266 g/mol.

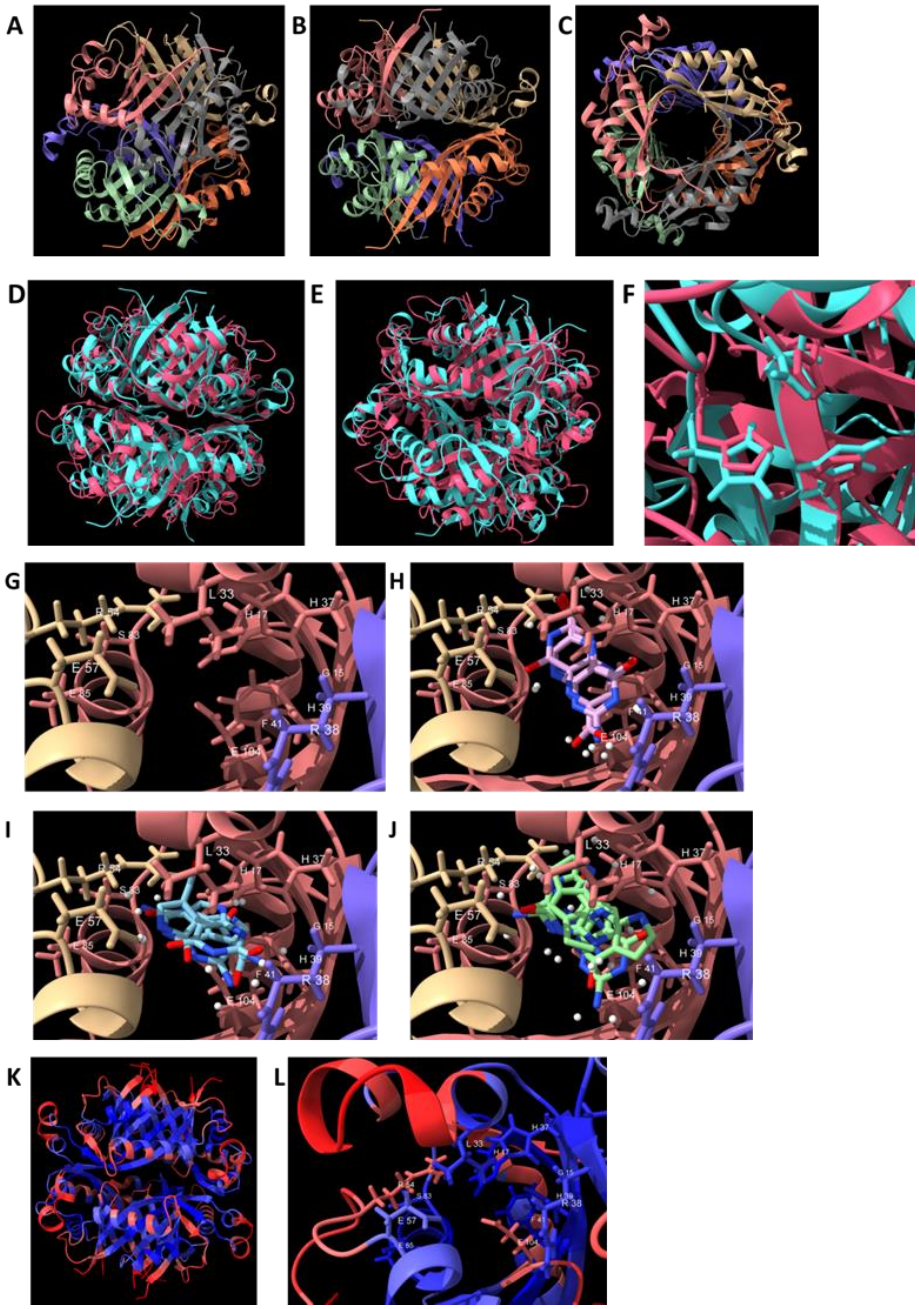

**Fig. S8. DpdL structure prediction**

Multimer AlphaFold2 prediction of Cellulophaga phage phiST DpdL (YP_009006155.1) colored by monomer and visualized from three different points of view (A, B, and C). DpdL hexamer (light blue) and *E. coli* QueD (PDB:4NTK, pink) were overlayed for comparison, and view from two sides (D and E) and zoomed in the zinc binding pocket with the conserved cysteines (F). The substrate binding pocket was investigated showing annotated conserved residues (G), colored by protein monomer. $CPH_4$ (H), CDG (I), and DG (J) were predicted to fit in the substrate pocket using Vina. Flexibility of the protein was analyzed by Medusa (K) and zoomed in the substrate binding pocket (L). Blue represents high confidence of rigidity and red high confidence of flexibility.

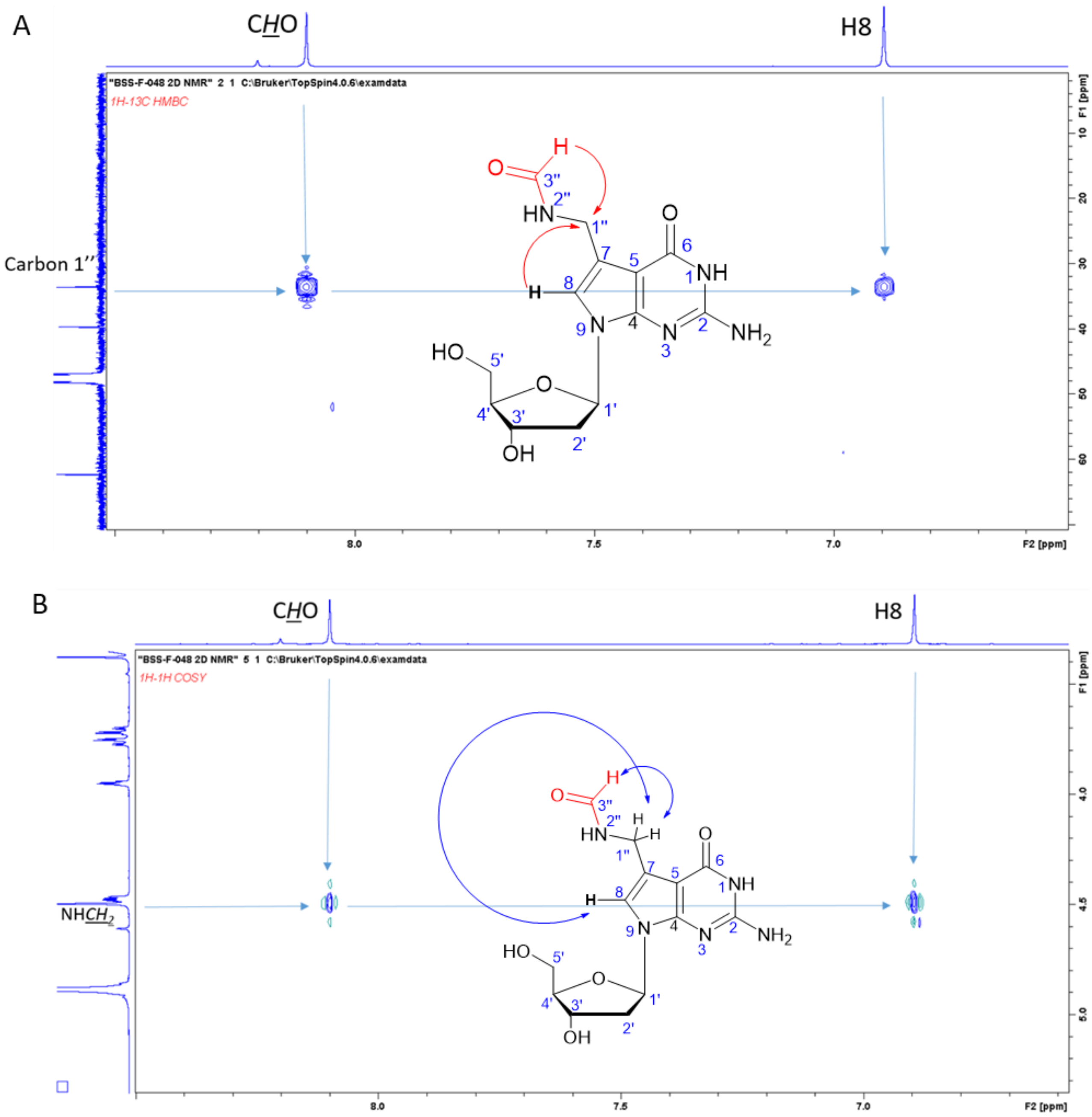

**Fig. S9. 2D NMR spectra of formyl-dPreQ1 (fdPreQ1) standard**

2D NMR confirms that formyl group is on nitrogen at C1''. (A) In the HMBC spectrum, the proton on formyl group (8.1 ppm) has a clear correlation with carbon of C1'' (33.6 ppm), the aromatic proton at C8 (6.89 ppm) has correlation to carbon of C1'' (33.6 ppm). (B) In the COSY spectrum, the proton on formyl group (8.1 ppm) has a clear correlation with singlet of NH*CH*$_2$ at C1'' (4.49 ppm), and the aromatic proton at C8 (6.89 ppm) has correlation to carbon of NH*CH*$_2$ at C1'' (4.49 ppm).

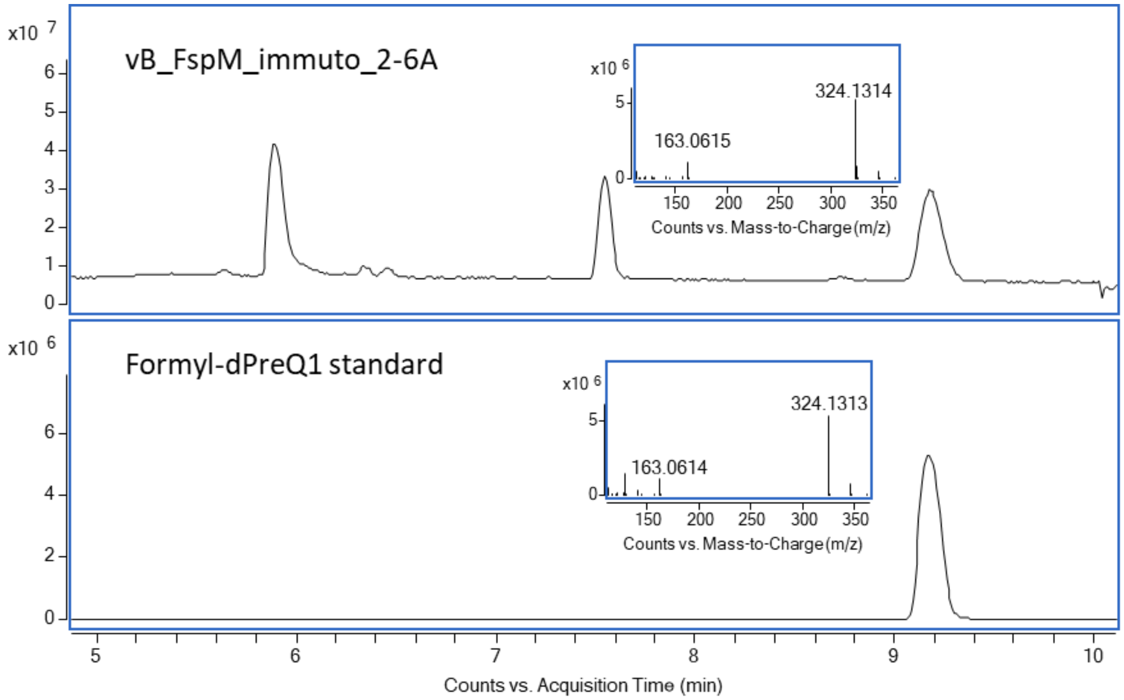

**Fig. S10. LC-MS analysis of digested vB_FspM_immuto_2-6A DNA sample and formyl-dPreQ1 standard**

The retention time and MS/MS spectra of the unknown non-canonical nucleoside in vB_FspM_immuto_2-6A (upper) were identical to those of the formyl-dPreQ$_1$ standard (lower)

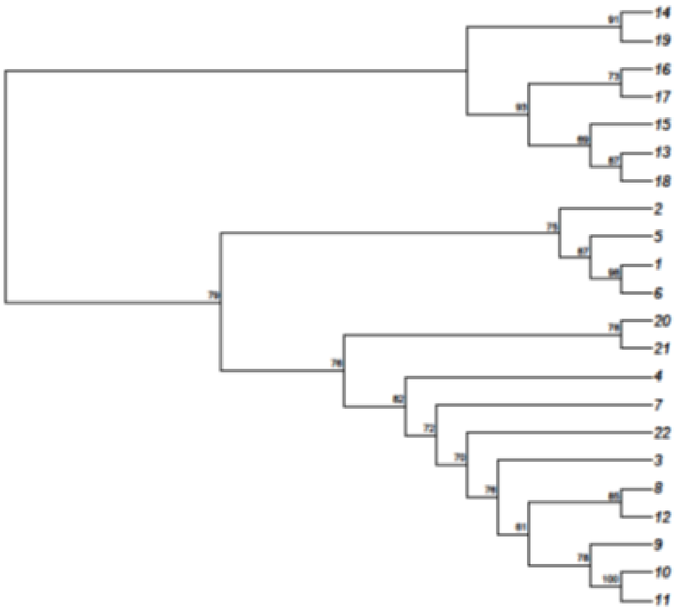

| ID | Name |
|---|---|
| 1 | YP_010114479.1 |
| 2 | CAB5226463.1 |
| 3 | CAH1079710.1 |
| 4 | CAH1082857.1 |
| 5 | CAB5221950.1 |
| 6 | CAB4142580.1 |
| 7 | CAH1078398.1 |
| 8 | BCU93200.1 |
| 9 | YP_004324978.1 |
| 10 | AGN12222.1 |
| 11 | YP_214283.1 |
| 12 | YP_007674480.1 |
| 13 | UJE15658.1 |
| 14 | CAB4130405.1 |
| 15 | QGH80014.1 |
| 16 | QDF17535.1 |
| 17 | URM86103.1 |
| 18 | QNL30130.1 |
| 19 | YP_001111230.1 |
| 20 | AYV83854.1 |
| 21 | AYV80424.1 |
| 22 | AYV82304.1 |

**Fig. S11. DpdN clustering tree**

Tree generated using Graph Splitting with all the DpdN sequences in Data S1. Values on the tree are Edge Perpetuation value and represent a confidence score in percent.

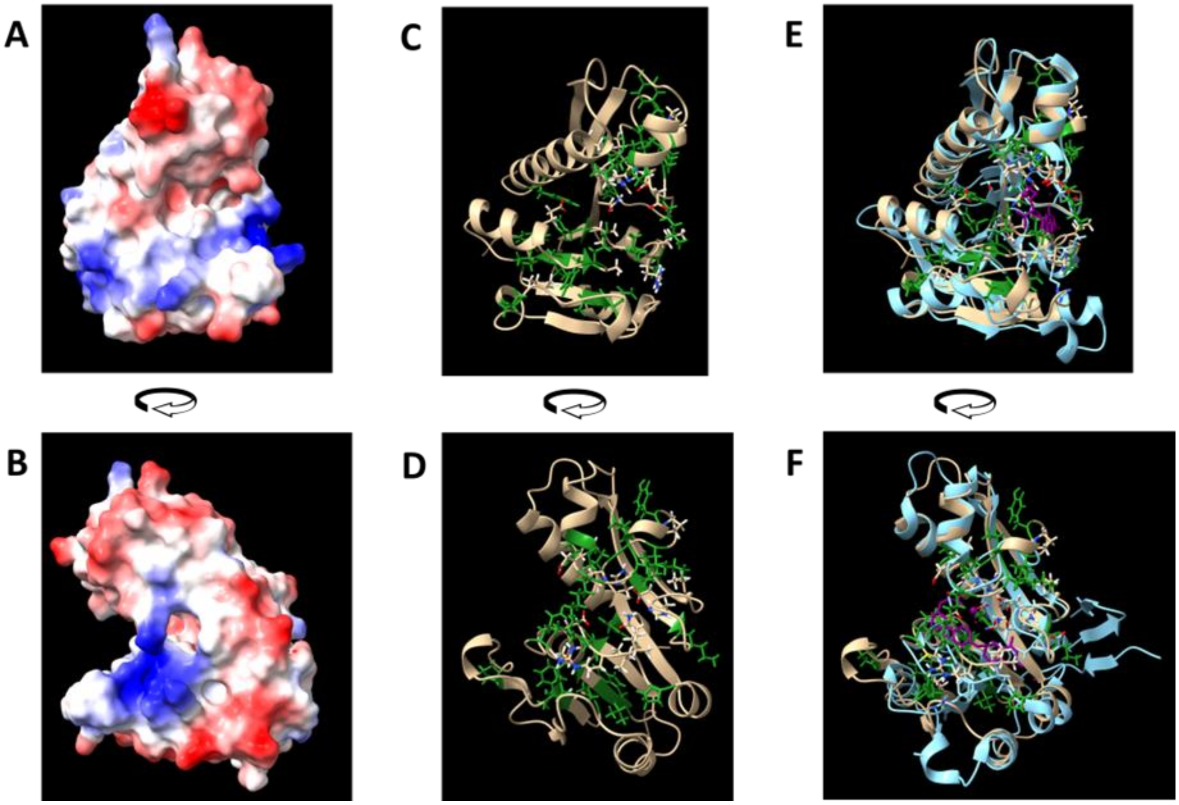

**Fig. S12. DpdN structure analysis**

The structure of Flavobacterium phage vB_FspM_immuto_2-6A DpdN was predicted with AlphaFold2 and visualized by electromagnetic surface charges on the face of the protein (A) and the side (B), blue are positive charges and red negative charges. Conserved residues were visualized with the ribbon representation (C and D), green residues are conserved residues in the four proposed true DpdN (Flavobacterium phage vB_FspM_immuto_2-6A DpdN numbering: W5, V6, A7, F9, Q11, P27, T32, N33, P44 H78, L81, P85, G100, I102, Y105, L108, K109, K111, P113, Q114, V128, K131, and W169). vB_FspM_immuto_2-6A DpdN was overlayed with *Mycobacterium tuberculosis* PurN (PDB:3DCJ, light blue in E and F). The co-crystalized formyl donor of PurN, 5-methyl-5,6,7,8-tetrahydrofolate, is colored in purple.

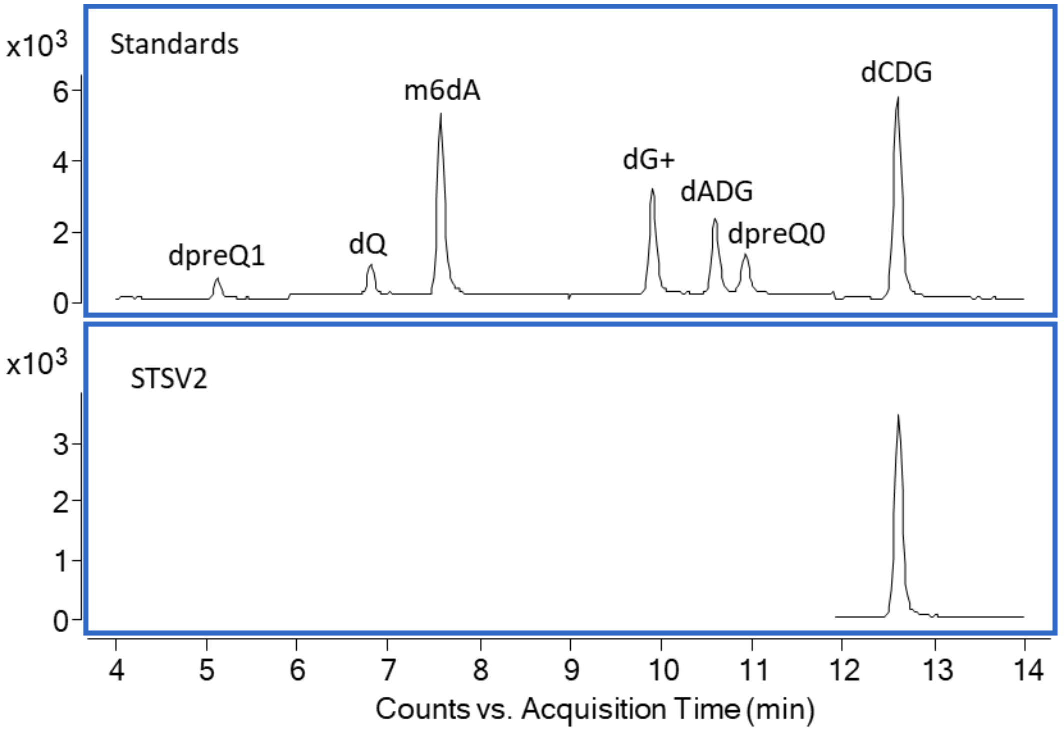

Fig. S13. **LC-MS analysis of digested Sulpholobus phage STSV-2 DNA sample and 2'-deoxy-7-deazaguanosine derivative standards**

The retention time and MRM transition (311→177) of the unknown non-canonical nucleoside in STSV-2 DNA (lower) were identical to those of the 2'-deoxy-7-carboxy-7-deazaguanine (dCDG) standard (upper).

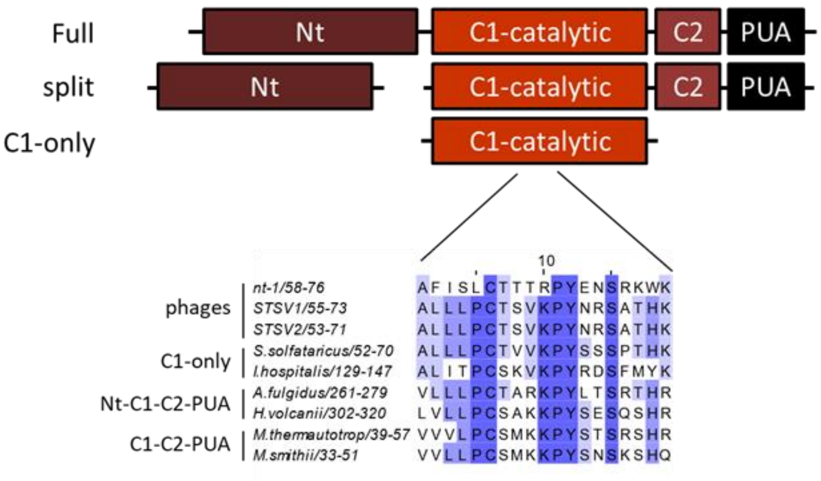

**Fig. S14. ArcS2 of phages contain only the core catalytic domain.**

ArcS contains four domains (nt, C1-catalytic, C2 and PUA), and has been found encoded either as a full enzyme or split between the Nt domain and the three others (12). Some only have the C1-catalitic domain, as in the phages. This catalytic domain is characterized by the PC-X3-KPY-X2-S-X2-H motif shared by all ArcS in the alignment.

#### Supplemental schemes

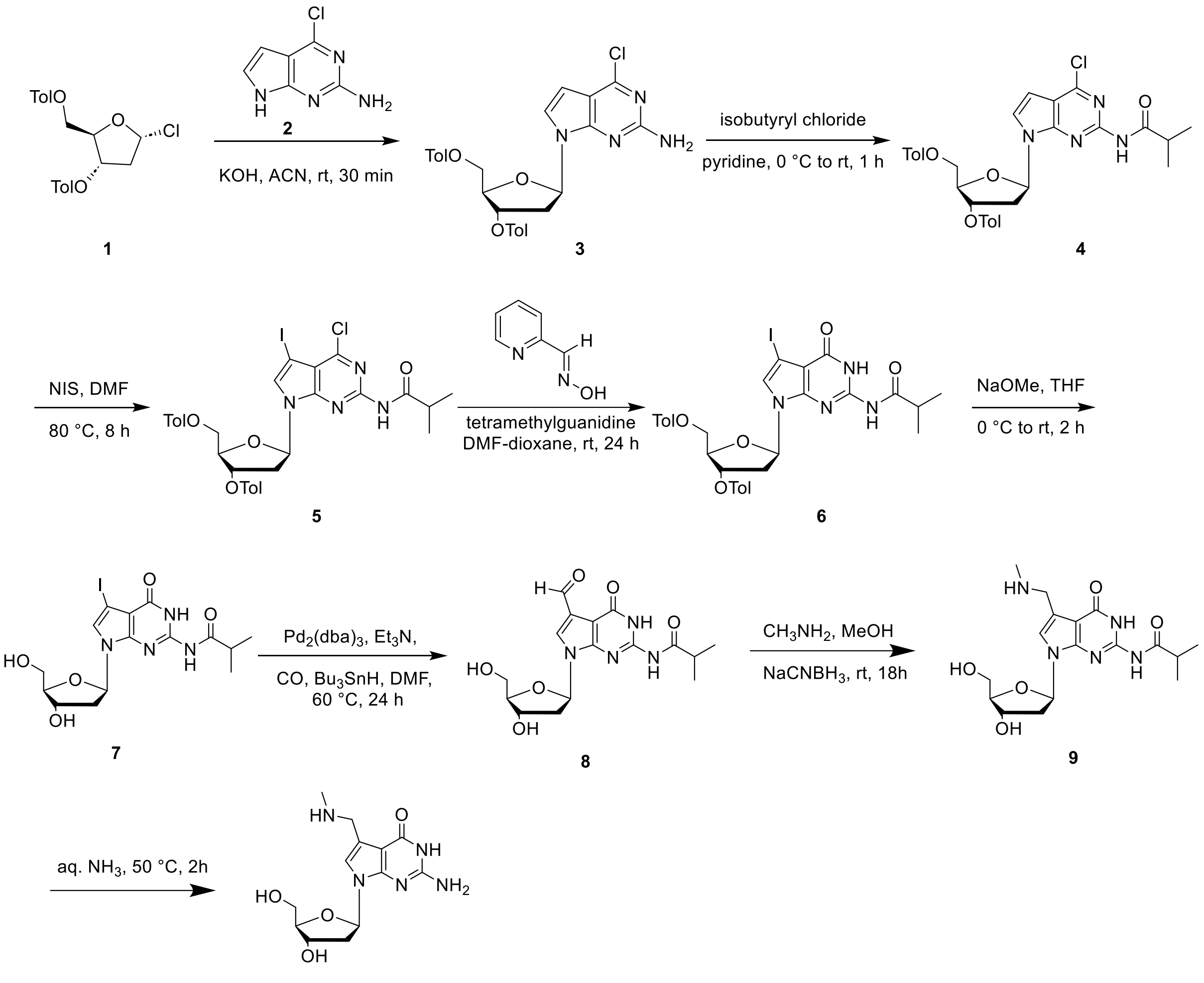

**Methyl-dPreQ$_1$ (10)**

**Scheme S1. Synthesis of methyl-dPreQ1.**

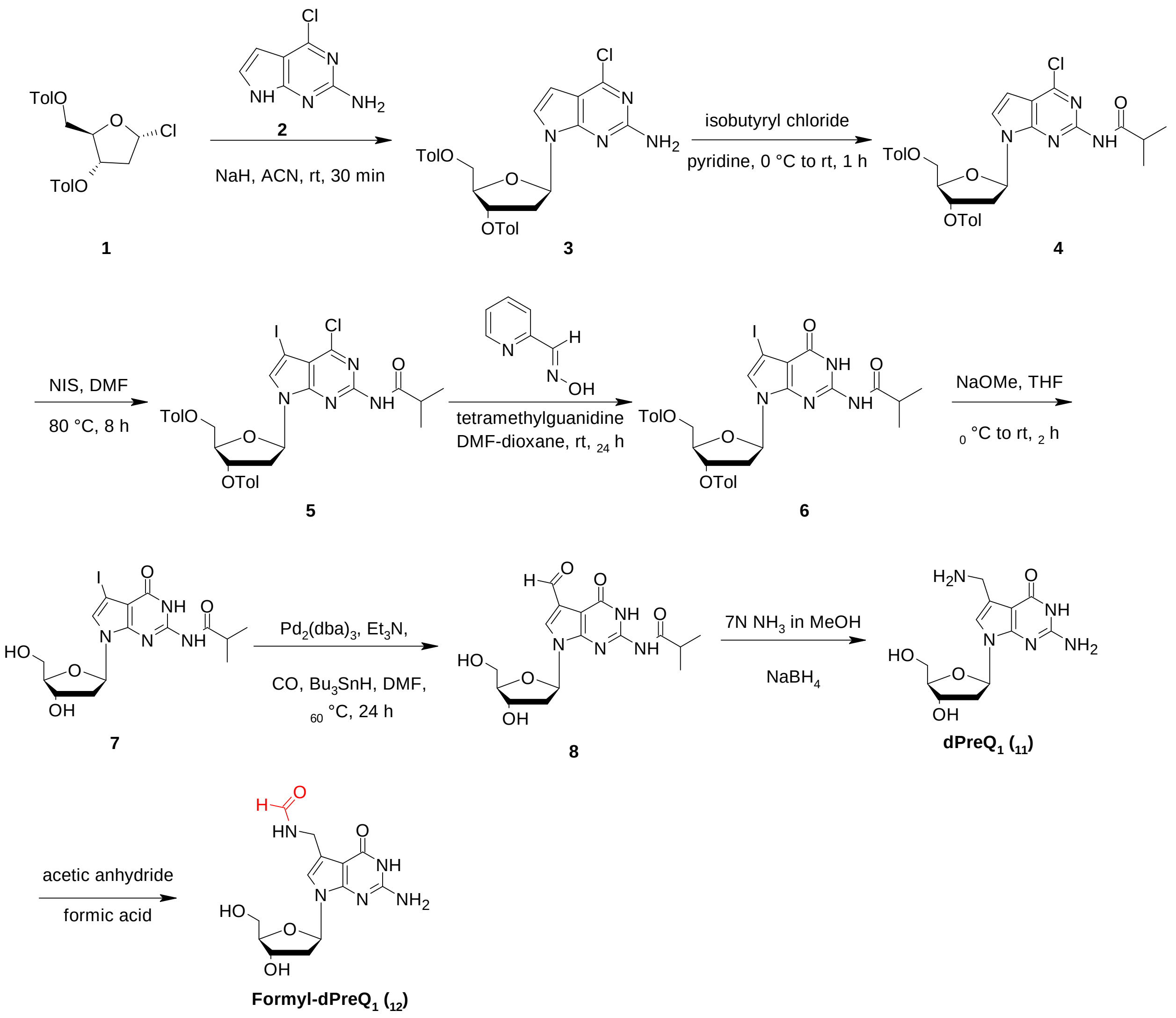

**Scheme S2. Synthesis of formyl-dPreQ1.**

**Supplemental tables**

| strain name | background | plasmid(s) | reference |
|---|---|---|---|
| **GC10** | F- mrcA D(mrr-hsdRMS-mrcBC) F80dlacZDM15 DlacX74 endA1 recA1 D(ara, leu)7697 araD139 galU galK nupG rpsL l T1R | none | Genesee Scientific Cat #: 42-658 |
| **MG1655** | F- λ- ilvG- rfb-50 rph-1 | none | (13) |
| **VDC2043** | MG1655 queD::FRT | none | (14) |
| **VDC7012** | MG1655 queC::FRT | none | natcoomm |
| **GJH2179** | MG1655 queF::kan | none | this study |
| **GJH1132** | MG1655 | pBAD24 | natcoomm |
| **GJH1510** | MG1655 | pBAD24/pBAD33 | natcoomm |
| **GJH2113** | GC10 | pGH116 | this study |
| **GJH2136** | GC10 | pGH126 | this study |
| **GJH2138** | GC10 | pGH128 | this study |
| **GJH2141** | GC10 | pGH118 | this study |
| **GJH2144** | GC10 | pGH120 | this study |
| **GJH2148** | GC10 | pGH129 | this study |
| **GJH2152** | GC10 | pGH121 | this study |
| **GJH2175** | GC10 | pGH127 | this study |
| **GJH2176** | GC10 | pGH119 | this study |
| **GJH2204** | MG1655 | pGH118/pBAD33 | this study |
| **GJH2207** | MG1655 | pBAD24/pGH121 | this study |
| **GJH2210** | MG1655 | pGH118/pGH121 | this study |
| **GJH2213** | MG1655 | pGH120/pBAD33 | this study |
| **GJH2216** | MG1655 | pBAD24/pGH119 | this study |
| **GJH2219** | MG1655 | pGH120/pGH119 | this study |
| **GJH2222** | MG1655 | pGH126/pBAD33 | this study |
| **GJH2225** | MG1655 | pBAD24/pGH129 | this study |
| **GJH2228** | MG1655 | pGH126/pGH129 | this study |
| **GJH2231** | MG1655 | pGH128/pBAD33 | this study |
| **GJH2234** | MG1655 | pBAD24/pGH127 | this study |
| **GJH2237** | MG1655 | pGH128/pGH127 | this study |
| **GJH2283** | VDC2043 | pBAD24/pBAD33 | this study |
| **GJH2286** | VDC2043 | pGH118/pBAD33 | this study |
| **GJH2289** | VDC2043 | pBAD24/pGH121 | this study |
| **GJH2292** | VDC2043 | pGH118/pGH121 | this study |
| **GJH2295** | VDC2043 | pGH120/pBAD33 | this study |
| **GJH2298** | VDC2043 | pBAD24/pGH119 | this study |
| **GJH2301** | VDC2043 | pGH120/pGH119 | this study |

| | | | |
|---|---|---|---|
| **GJH2425** | MG1655 | pGH118 | this study |
| **GJH2494** | GC10 | pGH162 | this study |
| **GJH2496** | GC10 | pGH164 | this study |
| **GJH2498** | GC10 | pGH166 | this study |
| **GJH2501** | GC10 | pGH168 | this study |
| **GJH2502** | GC10 | pGH170 | this study |
| **GJH2504** | GC10 | pGH172 | this study |
| **GJH2507** | GC10 | pGH178 | this study |
| **GJH2514** | GC10 | pGH163 | this study |
| **GJH2516** | GC10 | pGH165 | this study |
| **GJH2519** | GC10 | pGH167 | this study |
| **GJH2522** | GC10 | pGH171 | this study |
| **GJH2524** | GC10 | pGH173 | this study |
| **GJH2528** | GC10 | pGH177 | this study |
| **GJH2534** | GC10 | pGH174 | this study |
| **GJH2540** | GC10 | pGH176 | this study |
| **GJH2546** | MG1655 | pGH166 | this study |
| **GJH2548** | MG1655 | pGH172 | this study |
| **GJH2562** | MG1655 | pGH162/pBAD33 | this study |
| **GJH2564** | MG1655 | pBAD24/pGH165 | this study |
| **GJH2566** | MG1655 | pGH162/pGH165 | this study |
| **GJH2568** | MG1655 | pGH164/pBAD33 | this study |
| **GJH2570** | MG1655 | pBAD24/pGH163 | this study |
| **GJH2572** | MG1655 | pGH164/pGH163 | this study |
| **GJH2574** | MG1655 | pGH166/pBAD33 | this study |
| **GJH2576** | MG1655 | pBAD24/pGH171 | this study |
| **GJH2578** | MG1655 | pGH166/pGH171 | this study |
| **GJH2580** | MG1655 | pGH170/pBAD33 | this study |
| **GJH2582** | MG1655 | pBAD24/pGH167 | this study |
| **GJH2584** | MG1655 | pGH170/pGH167 | this study |
| **GJH2586** | MG1655 | pGH172/pBAD33 | this study |
| **GJH2588** | MG1655 | pBAD24/pGH177 | this study |
| **GJH2590** | MG1655 | pGH172/pGH177 | this study |
| **GJH2592** | MG1655 | pGH176/pBAD33 | this study |
| **GJH2594** | MG1655 | pBAD24/pGH173 | this study |
| **GJH2596** | MG1655 | pGH176/pGH173 | this study |
| **GJH2626** | GJH2179 | pGH166/pBAD33 | this study |
| **GJH2628** | GJH2179 | pBAD24/pGH171 | this study |
| **GJH2630** | GJH2179 | pGH166/pGH171 | this study |
| **GJH2632** | GJH2179 | pGH170/pBAD33 | this study |
| **GJH2634** | GJH2179 | pBAD24/pGH167 | this study |

| | | | |
|---|---|---|---|
| **GJH2636** | GJH2179 | pGH170/pGH167 | this study |
| **JMH1050** | GJH2179 | pBAD24 | this study |
| **JMH1051** | GJH2179 | pGH178 | this study |
| **JMH1052** | GJH2179 | pGH168 | this study |
| **JMH1053** | GJH2179 | pGH174 | this study |
| **JMH1054** | GJH2179 | pGH116 | this study |
| **JMH1068** | VDC7012 | pBAD24/pGH167 | this study |
| **JMH1070** | VDC7012 | pGH170/pGH167 | this study |
| **JMH1074** | VDC7012 | pGH166/pBAD33 | this study |
| **JMH1076** | VDC7012 | pGH166/pGH171 | this study |

**Table S1. Strain list**

| plasmid name | backbone | inserted gene | reference |
|---|---|---|---|
| **pBAD24** | pBAD24 | none | (15) |
| **pBAD33** | pBAD33 | none | (15) |
| **pGH116** | pBAD24 | HVTV-1 *queF* | this study |
| **pGH118** | pBAD24 | nt-1 *dpdA2* | this study |
| **pGH119** | pBAD33 | nt-1 *dpdA2* | this study |
| **pGH120** | pBAD24 | nt-1 *arcS2* | this study |
| **pGH121** | pBAD33 | nt-1 *arcS2* | this study |
| **pGH126** | pBAD24 | phiST *dpdA3* | this study |
| **pGH127** | pBAD33 | phiST *dpdA3* | this study |
| **pGH128** | pBAD24 | phiST *dpdL* | this study |
| **pGH129** | pBAD33 | phiST *dpdL* | this study |
| **pGH162** | pBAD24 | ACP17 *dpdA2* | this study |
| **pGH163** | pBAD33 | ACP17 *dpdA2* | this study |
| **pGH164** | pBAD24 | ACP17 *dpdL* | this study |
| **pGH165** | pBAD33 | ACP17 *dpdL* | this study |
| **pGH166** | pBAD24 | VH7D *dpdA2* | this study |
| **pGH167** | pBAD33 | VH7D *dpdA2* | this study |
| **pGH168** | pBAD24 | VH7D *queF* | this study |
| **pGH170** | pBAD24 | VH7D *dpdM* | this study |
| **pGH171** | pBAD33 | VH7D *dpdM* | this study |
| **pGH172** | pBAD24 | phiSM *dpdA* | this study |
| **pGH173** | pBAD33 | phiSM *dpdA* | this study |
| **pGH174** | pBAD24 | phiSM *queF* | this study |
| **pGH176** | pBAD24 | phiSM *dpdM* | this study |
| **pGH177** | pBAD33 | phiSM *dpdM* | this study |
| **pGH178** | pBAD24 | *E. coli queF* | this study |

**Table S2. Plasmid list**

**Supplementary data**

**Data S1. (separate file)**

Protein involved in the 7-deazaguanine DNA modification pathway encoded in several phages

**Data S2. (separate file)**
psiBLAST results of CEPG_00048 against Viruses (taxid:10239)

**Data S3. (separate file)**
psiBLAST results of CEPG_00057 against Viruses (taxid:10239)

**Data S4. (separate file)**
psiBLAST results of CEPG_00056 against Viruses (taxid:10239)

**Data S5. (separate file)**
psiBLAST results of CEPG_00054 against Viruses (taxid:10239)

**Data S6. (separate file)**
HHpred results of CEPG_00054 against pFam

**Data S7. (separate file)**
psiBLAST results of CGPG_00064 against Viruses (taxid:10239)

**Data S8. (separate file)**
psiBLAST results of CGPG_00066 against Viruses (taxid:10239)

**Data S9. (separate file)**
HHpred results of CGPG_00068 against pFam

**Data S10. (separate file)**
HHpred results of CGPG_00065 against pFam

**Data S11. (separate file)**
HHpred results of CGPG_00067 against pFam

**Data S12. (separate file)**
psiBLAST results of CGPG_00067 against Viruses (taxid:10239)

**Data S13. (separate file)**
psiBLAST results of CGPG_00065 against Viruses (taxid:10239)

**Data S14. (separate file)**
HHpred results of HVTV1_69 against pFam

**Data S15. (separate file)**
psiBLAST results of HVTV1_69 against Viruses (taxid:10239)

**Data S16. (separate file)**

psiBLAST results of KNV73_gp067 against Viruses (taxid:10239)

**Data S17. (separate file)**

HHpred results of STSV2_16 against pFam

**Data S18. (separate file)**

HHpred results of VPFG_00169 against pFam
